# Reconstructing multi-scale tissue spatial architecture from single-cell RNA-seq with REMAP

**DOI:** 10.64898/2026.02.21.707167

**Authors:** Shunzhou Jiang, Kyle Coleman, Zihao Chen, Kaitian Jin, Yunhe Liu, Dong Heon Lee, Tae Hyun Hwang, Rui Xiao, Jin Jin, Christopher A. Walsh, Xuyu Qian, Linghua Wang, Mingyao Li

**Affiliations:** Statistical Center for Single-Cell and Spatial Genomics, Department of Biostatistics, Epidemiology and Informatics, Perelman School of Medicine, University of Pennsylvania, Philadelphia, PA, USA; Department of Computational Biomedicine, Cedars Sinai Medical Center, Los Angeles, CA, USA; Department of Genomic Medicine, University of Texas MD Anderson Cancer Center, Houston, TX, USA; Department of Surgery, Vanderbilt University Medical Center, Nashville, TN, USA; Division of Genetics and Genomics, Department of Pediatrics, Boston Children’s Hospital, Harvard Medical School, Boston, MA, USA; Howard Hughes Medical Institute, Boston Children’s Hospital, Harvard Medical School, Boston, MA, USA; Department of Pediatrics, Division of Neurology, University of Pennsylvania Perelman School of Medicine, Philadelphia, PA, USA; Center for Brain Research in Development, Genetics, and Engineering (BRIDGE), Children’s Hospital of Philadelphia, PA, USA; Department of Pathology and Laboratory Medicine, Perelman School of Medicine, University of Pennsylvania, Philadelphia, PA, USA

**Keywords:** Single-cell RNA-seq, Spatial transcriptomics, Cellular neighborhood, Deep learning

## Abstract

Understanding spatial organization of cells is critical for deciphering tissue function and disease. Single-cell RNA-sequencing (scRNA-seq) profiles transcriptomes at scale but loses spatial context, while spatial transcriptomics (ST) preserves spatial information but is constrained by cost and gene coverage. Here, we present REMAP, a deep learning framework that integrates gene expression with neighborhood-level gene-gene covariance to reconstruct multi-scale spatial organization of scRNA-seq data using one or multiple ST references. Across 2D and 3D mouse brain, human fetal cortex, and seven human cancer types, REMAP consistently outperformed existing approaches. Applied to a human multiple sclerosis atlas, REMAP resolved microglial neighborhood heterogeneity, and identified a rare pro-inflammatory microglia-astrocyte subpopulation. Across diverse cancers, REMAP recovered conserved spatially-defined cancer-associated fibroblast subtypes with known prognostic significance. By transforming cost-efficient single-cell datasets into spatially interpretable tissue maps, REMAP enables spatial hypothesis generation, microenvironment discovery, and population-scale inference of conserved and perturbed architectural principles in human disease.

## Introduction

Single-cell RNA-sequencing (scRNA-seq) has emerged as a powerful technology for dissecting the transcriptional landscape of complex tissues at single-cell resolution. A major limitation, however, is the loss of spatial context, as dissociated cells lack information on their original location. Spatial information is essential for understanding tissue organization, cellular microenvironments, and intercellular communications. More recently developed spatial transcriptomics (ST) technologies address this drawback by profiling gene expression while maintaining cell locations in their native tissue context^1-7^. However, the high costs decrease the accessibility of those technologies and limit their applications in population-level studies. Given the complementary nature of scRNA-seq and ST technologies, integrating these two technologies is essential to overcome their individual limitations^8-11^. Since scRNA-seq is more cost-effective and extensive scRNA-seq cell atlas datasets have already been generated, it is highly desirable to reconstruct the spatial locations of cells in scRNA-seq by leveraging ST data as a reference.

Several computational approaches have been developed to address this challenge. The first category of these methods aims to find a mapping between scRNA-seq and ST data, assigning each single cell to the location of a ST cell or spot with the highest mapping probability. Representative methods include Tangram^12^, SpaOTsc^13^, novoSpaRc^14^, and CellContrast^15^. While these methods are suitable for ST-centric tasks such as missing gene expression imputation and cell type deconvolution, when predicting cell locations for scRNA-seq, each single cell can only be mapped to one of the existing locations in the ST reference, which lacks flexibility and generalizability. Another category of methods directly predicts the spatial locations of scRNA-seq cells. Representative methods include CeLEry^16^, iSORT^17^, and LUNA^18^. While these methods have shown promising performance, a common limitation is that they fail to consider the local neighborhood context, which is crucial for capturing cell-cell interactions and accurate representation of cell states beyond individual gene expression alone^19-22^. Moreover, except for LUNA, all existing methods rely on a single ST reference, which is limiting given the increasing availability of diverse ST datasets. Furthermore, the physical size of many tissue samples exceeds the capture area of standard ST platforms. As a result, multiple subregions are often required to cover an entire sample^23^. Consequently, it is vital to integrate all ST captures as references so that the model can learn spatial relationships at the whole-tissue level.

To address the limitations of existing methods, we developed REMAP (**RE**constructing **M**ulti-scale tissue **A**rchitecture from single-cell **P**rofiles), a deep learning framework designed to reconstruct the spatial locations for cells in scRNA-seq. REMAP leverages single or multiple ST references and incorporates gene expression and cellular neighborhood context to guide spatial reconstruction. In addition to predicting absolute coordinates, REMAP is also able to infer the cellular neighborhood (CN) spatial network, which captures high-level spatial relationships of scRNA-seq cells and identifies which cells are spatial neighbors and which cell groups are colocalized. Through comprehensive benchmarking evaluations on data generated from diverse tissues and platforms, we show that REMAP accurately reconstructs global and local tissue spatial architecture, significantly surpassing existing tools visually and quantitatively. We further illustrate REMAP’s utility by analyzing a human multiple sclerosis atlas, where REMAP enabled comparisons of spatial niches across disease states and uncovered context-specific organization of microglia, including previously unrecognized patterns within inactive lesions. Extending further to cancer, REMAP revealed conserved spatial subtypes of cancer-associated fibroblasts across multiple cancer types that align with known functional states. Together, these results position REMAP as a general-purpose and robust framework for integrating ST and scRNA-seq data, supporting the development and biological discovery of the Human Cell Atlas^24^.

## Results

### Overview of REMAP

REMAP reconstructs the spatial locations of cells in scRNA-seq by integrating both first-order gene expression and second-order gene-gene covariance features, using ST as references (**Fig. 1**). First-order features capture the gene expression patterns of individual cells, while second-order features characterize gene-gene relationships derived from neighboring cells, which encode information about local tissue microenvironment^19-22^. During training, REMAP identifies spatial neighbors for each cell in the ST reference and computes gene-gene covariance within these neighboring cells to capture their local context. This neighborhood covariance is combined with gene expression to train a deep neural network that predicts cell locations. During the scRNA-seq inference stage, the spatial neighbors are unknown as scRNA-seq lacks location information. To enable neighborhood covariance estimation, REMAP first adopts ENVI^21^, which leverages ST data to infer such covariance in scRNA-seq, and then iteratively refines it by training a secondary model that predicts neighborhood covariance from gene expression and preliminary location estimates. This iterative updating procedure improves the neighborhood context accuracy for scRNA-seq cells and enhances the effectiveness of the location prediction model trained on ST.

**Fig. 1:**
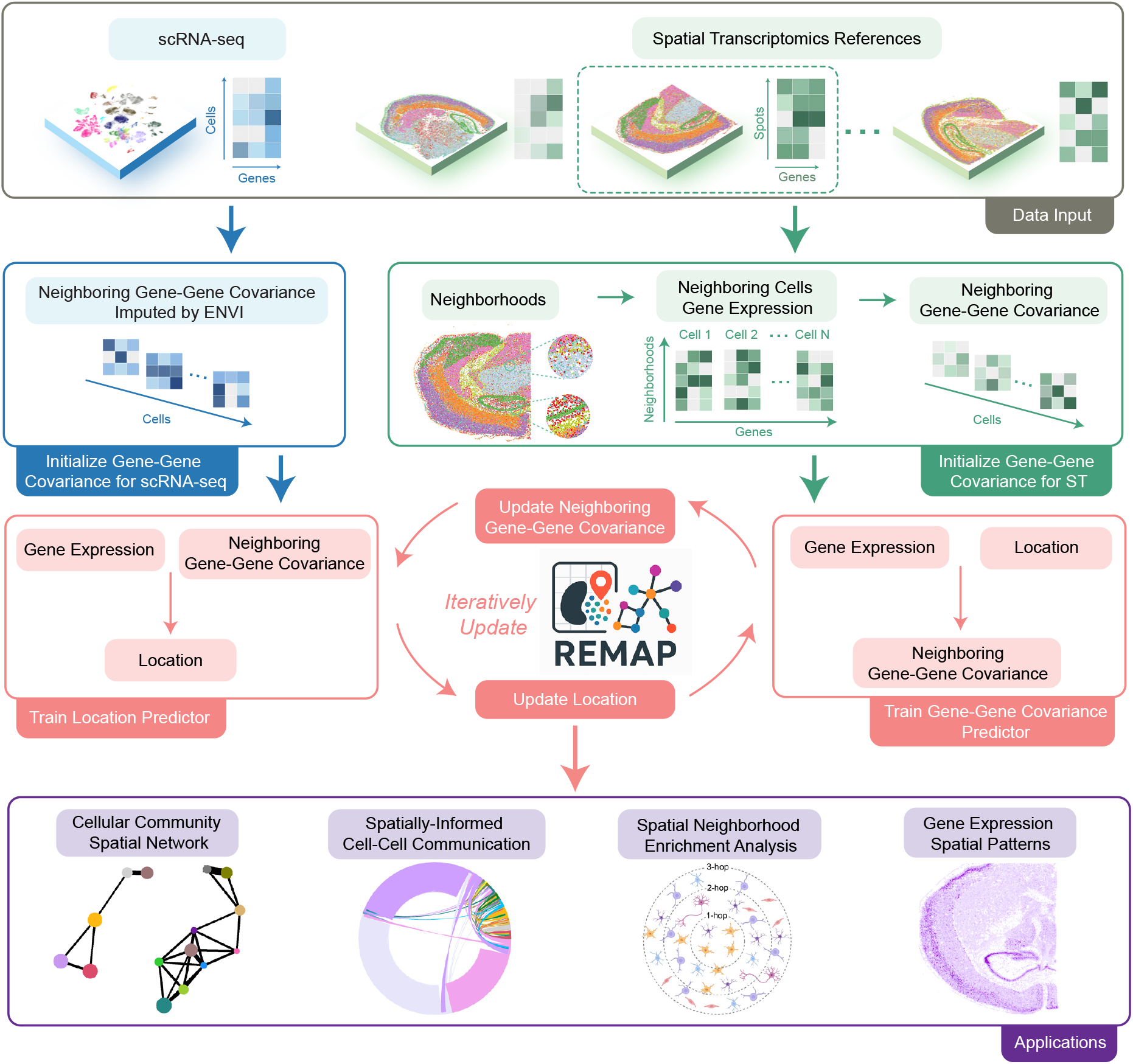
Overview of REMAP. REMAP takes single or multiple ST sections as input for training and reconstructs the spatial locations of cells in scRNA-seq data by integrating first-order gene expression and second-order cellular neighborhood information. To initialize the neighborhood representation for ST, REMAP identifies spatial neighbors and computes gene-gene covariance within neighbors for every ST cell. While for scRNA-seq, where cell locations are unknown, gene-gene covariance for neighboring cells is initialized using ENVI^21^. Gene expression and initialized neighborhood covariance are combined to train a location predictor, which predicts cell locations. Next, a secondary predictor refines neighborhood covariance estimates based on gene expression and predicted locations. These two predictors are iteratively trained to update location and neighborhood covariance estimates. After obtaining the final predicted locations for scRNA-seq, various downstream applications can be performed, including cellular community spatial network construction, spatially-informed cell-cell communication analysis, spatial neighborhood enrichment analysis, and gene expression spatial patterns visualization.

When multiple ST references are available, the individual references may differ in orientation or tissue coverage, making absolute spatial coordinates difficult to compare or align. In contrast, pairwise cell-to-cell distances remain comparable and therefore provide a reliable basis for integration across different ST references. REMAP leverages this property to learn the relationship between pairwise distances and combined first-order and second-order molecular information to predict a pairwise distance matrix for scRNA-seq cells. This matrix enables the identification of spatial neighbors, clustering of cells with similar neighborhood composition, and inference of CN spatial proximity. Multidimensional scaling (MDS) can be applied to the predicted pairwise distances to generate 2D visualization of cell locations. Moreover, when 3D location is available in the reference ST data, REMAP can predict 3D spatial origins, which better reflect the authentic tissue architecture. The predicted cell locations from REMAP enable a wide range of downstream applications, including spatially informed inference of cell-cell communication, spatial neighborhood enrichment analysis, and visualization of spatial expression patterns for genes not measured in ST.

### REMAP recovered fine-grained shapes of the mouse brain

To benchmark REMAP against existing methods, we analyzed a mouse brain dataset generated with 10x Visium HD, masking spatial coordinates, and applied REMAP alongside competing methods to recover cell locations using a reference mouse brain dataset profiled with 10x Xenium (**Fig. 2a**). Because the test data covered only half of the brain, we split the reference into two hemispheres and restricted training to the left hemisphere. Notably, the training and test datasets originated from distinct mice and were generated by different ST platforms. To mitigate gene expression batch effects, we first applied CarDEC^25^ and used the batch-corrected expressions as input for all methods. **Fig. 2b** shows that REMAP faithfully reconstructed the tissue architecture in the test data, uniquely recovering the hippocampal shape and delineating its three subregions. In contrast, CeLEry, iSORT, and LUNA localized cells only to approximate regions without capturing delicate spatial structures, while CellContrast collapsed multiple cells into identical coordinates, producing sparse and distorted spatial maps. More detailed cluster-specific comparisons are provided in **Supplementary Fig. 1**.

**Fig. 2:**
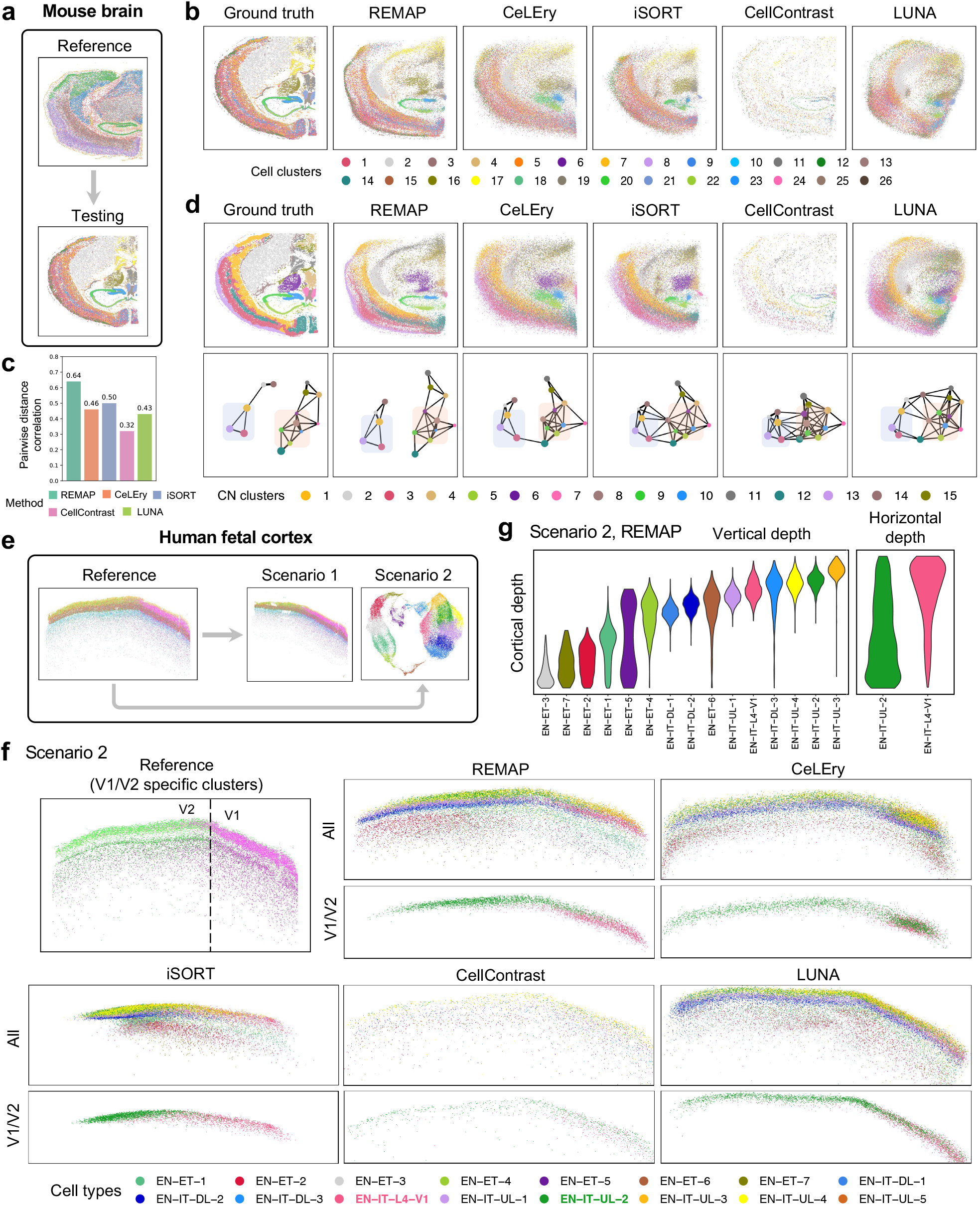
Application to mouse brain and human fetal cortex data. **a**, Study design for the mouse brain data benchmarking evaluation, where 10x Xenium data serves as the reference and 10x Visium HD data serves as the test dataset. **b**, Ground-truth and predicted cell locations by REMAP and other methods. **c**, Pearson correlations between true and predicted pairwise distances. **d**, CN clusters and spatial networks constructed by REMAP and other methods. In the top panel, cells were plotted under predicted locations and colored by ground-truth CN clusters; in the bottom panel, nodes represent the centroids of CN clusters with edges drawn based on spatial proximity. Edge length corresponds to the Euclidean distance between centroids, and edge width was scaled inversely to distance, such that thicker edges indicate closer proximity. Node size is proportional to the standard deviation of within-cluster pairwise distances among cells. Two regions of interest were highlighted with blue and orange shaded boxes. **e**, Study design for the human fetal cortex data evaluation, where one MERFISH slice was used as the reference, with either an adjacent MERFISH slice (Scenario 1) or paired snRNA-seq data (Scenario 2) as the test data. **f**, Predicted cell locations by REMAP and other methods in Scenario 2, with the top panel showing all cell types and the bottom panel displaying V1/V2 specific cell types. **g**, Vertical cortical depth distributions of all cell types and horizontal cortical depth distributions of V1/V2 specific cell types based on REMAP recovered locations in Scenario 2.

We further quantified prediction accuracy using the Pearson correlation between true and predicted pairwise distances across all cell pairs (**Fig. 2c**). REMAP achieved the highest correlation of 0.64. To assess REMAP’s robustness under imperfect ST references, we removed subsets of cortical and hippocampal cells from the ST reference and reran all methods (**Supplementary Fig. 2a**). Even under this challenging scenario, REMAP maintained strong performance with a pairwise distance correlation of 0.61, and correctly placed cells from the missing regions in their approximate locations (**Supplementary Fig. 2b**,**c**).

Next, we evaluated how well each method preserved global tissue architecture. Using the ground-truth cell coordinates, we first identified spatial neighborhoods for each cell and clustered them by cell composition within the neighborhood to detect CN clusters (see **Methods**). As shown in **Fig. 2d**, REMAP faithfully maintained the CN cluster structure; for example, REMAP was the only method capable of distinguishing Cluster 12 as a distinct cortical layer structure that did not intermix with Clusters 1 and 3. To characterize inter-cluster proximity, we constructed CN spatial networks (see **Methods**). The network generated by REMAP most closely resembled the ground truth (**Fig. 2d**); for instance, REMAP separated cortical layer-associated CN clusters (blue shaded box) and avoided spurious connections in the hippocampal surrounding regions (orange shaded box), whereas competing methods produced noisier networks with excessive edges.

We then computed Euclidean distances between centroids of all CN cluster pairs and compared predicted and true pairwise distances. As shown in **Supplementary Fig. 3a**, REMAP consistently yielded the lowest mean discrepancy across varying numbers of CN clusters, confirming its superior retention of global tissue structure. When CN clustering was performed directly on the predicted locations, REMAP again produced the most accurate CN networks (**Supplementary Fig. 3c**) and achieved the highest normalized mutual information (NMI) scores between predicted and ground-truth CN labels (**Supplementary Fig. 3b**,**d**). We further assessed whether REMAP-predicted locations were sufficient for downstream analyses of spatial cell-cell communication using SpatialDM^26^, which requires pairwise distance between cells (see **Methods**). REMAP-recovered positions maintained the majority of ligand-receptor communication signals (**Supplementary Fig. 4**), demonstrating robust preservation of biologically relevant spatial context.

Additionally, we evaluated REMAP’s performance using a scRNA-seq mouse primary motor cortex dataset^27^ as test data and a MERFISH dataset comprising multiple tissue slices^28^ as ST reference (**Supplementary Figs. 5-8**). REMAP-predicted locations in the scRNA-seq dataset exhibited clear cortical laminar separation and an even spatial distribution, faithfully recapitulating layer-specific marker gene expression and reconstructing laminar structures with multiple reference slices. Remarkably, in leave-one-out experiments excluding one cell type from the reference at a time (L4/5 IT, L5 IT, L6 IT, or L6 CT), REMAP accurately reassigned the omitted cell types to their correct cortical layers, whereas competing methods produced blurred or misplaced patterns.

### REMAP identified distinct cortical areal borders in the human fetal cortex

Encouraged by REMAP’s superior performance in mouse studies, we next applied it to a human cerebral cortex dataset^29^ comprising extensive MERFISH and paired single-nucleus RNA-sequencing (snRNA-seq) data from consecutive tissue sections collected during mid-gestation. This dataset captures both continuous six-layer laminar gradients and a sharp boundary between primary (V1) and secondary (V2) visual cortices, marked by V1- and V2-enriched clusters at gestational week 20. For this dataset, we designed two evaluation scenarios (**Fig. 2e**).

Scenario 1 involved two adjacent MERFISH slices, with one serving as the reference and the other as the test. REMAP sharply delineated the V1/V2 border and achieved the highest pairwise distance correlation (**Supplementary Fig. 9**). To summarize the complex spatial organization of cell types, we performed CN clustering to group cells with similar spatial context (**Supplementary Fig. 10**). Reducing ∼40 raw cell types into CN clusters produced clearer spatial patterns that captured both vertical laminar gradients and horizontal V1-V2 areal distinctions. Among all methods, REMAP best retained these CN structures, accurately resolving the vertical laminar transitions within both V1 and V2 territories. While other methods distinguished the V1/V2 split, they intermixed CN clusters within each side. Quantitatively, REMAP produced the lowest error in the pairwise distances between CN centroids and the highest concordance between true and predicted CN cluster assignments (**Supplementary Fig. 11**), consistent across cluster resolutions. Together, these results show that REMAP most effectively preserves the high-level spatial architecture of tissues.

Scenario 2 evaluated a real-world application in which the spatial locations of snRNA-seq cells were inferred. REMAP accurately delineated the V1/V2 border, localizing EN-IT-L4-V1 and EN-IT-UL-2 cells exclusively to their respective territories (**Fig. 2f**). In contrast, CeLEry, CellContrast, and LUNA failed to recover the V2-enriched spatial pattern of EN-IT-UL-2, while iSORT erroneously expanded the V1 region. To assess cortical depth patterns, we selected three cortical regions from the reference and quantified both vertical and horizontal depth distributions (**Supplementary Fig. 12a**). REMAP-recovered locations recapitulated progressive laminar specificity and preserved horizontal separation of V1- and V2-enriched clusters (**Fig. 2g**). Competing methods failed to capture both features (**Supplementary Fig. 12b**). CN spatial networks further highlighted REMAP’s accuracy: only REMAP eliminated spurious connections between V1- and V2-specific CN clusters (**Supplementary Fig. 13a**). Transcriptomic analysis of CN clusters confirmed biological relevance, with REMAP derived clusters strongly enriched for V1 and V2 markers (*ABI3BP, PDZRN4, NPY, IGFBPL1, and FLRT2*), as shown by mean expression and fold change analyses (**Supplementary Fig. 13b**). Moreover, heatmaps of additional genes not included in the MERFISH gene panel revealed distinct V1- or V2-enriched patterns (**Supplementary Fig. 14**), demonstrating REMAP’s ability to uncover biologically meaningful spatial architecture beyond the measured gene set.

### REMAP reconstructed curved and layered structures in human colorectal cancer

Building on REMAP’s success in brain tissue, we next tested its utility in cancer, a far more challenging setting. Unlike the brain, which exhibits highly conserved laminar and areal organization, tumors are characterized by pronounced heterogeneity, irregular structures, and complex microenvironments. These features complicate spatial reconstruction and demand greater model robustness. We applied REMAP to a colorectal cancer (CRC) sample from Patient 5, profiled with 10x Visium HD^30^, treating the spatial coordinates as unobserved. Given the substantial heterogeneity of cancer tissues across patients, we used the 10x Xenium data from an adjacent section of the same patient as the spatial reference (**Fig. 3a**). REMAP accurately reconstructed the complex tissue architecture (**Fig. 3b**) and achieved the highest preservation of pairwise distance relationships with a correlation of 0.64 (**Fig. 3c**). Notably, REMAP faithfully recovered the two curved structures and the layered organization of the goblet cell region. Competing methods failed to resolve these fine-grained patterns: CeLEry and iSORT collapsed curved structures into straight lines, while LUNA produced erroneous placements for several clusters (clusters 7, 20, and 21; **Supplementary Fig. 15**). Vertical depth quantification further confirmed REMAP’s accuracy as violin plots closely matched the reference and maintained area specificity for clusters 17, 6, 20, and 21. By contrast, competing methods lost area specificity or produced overly fragmented spatial patterns (**Supplementary Fig. 16**).

**Fig. 3:**
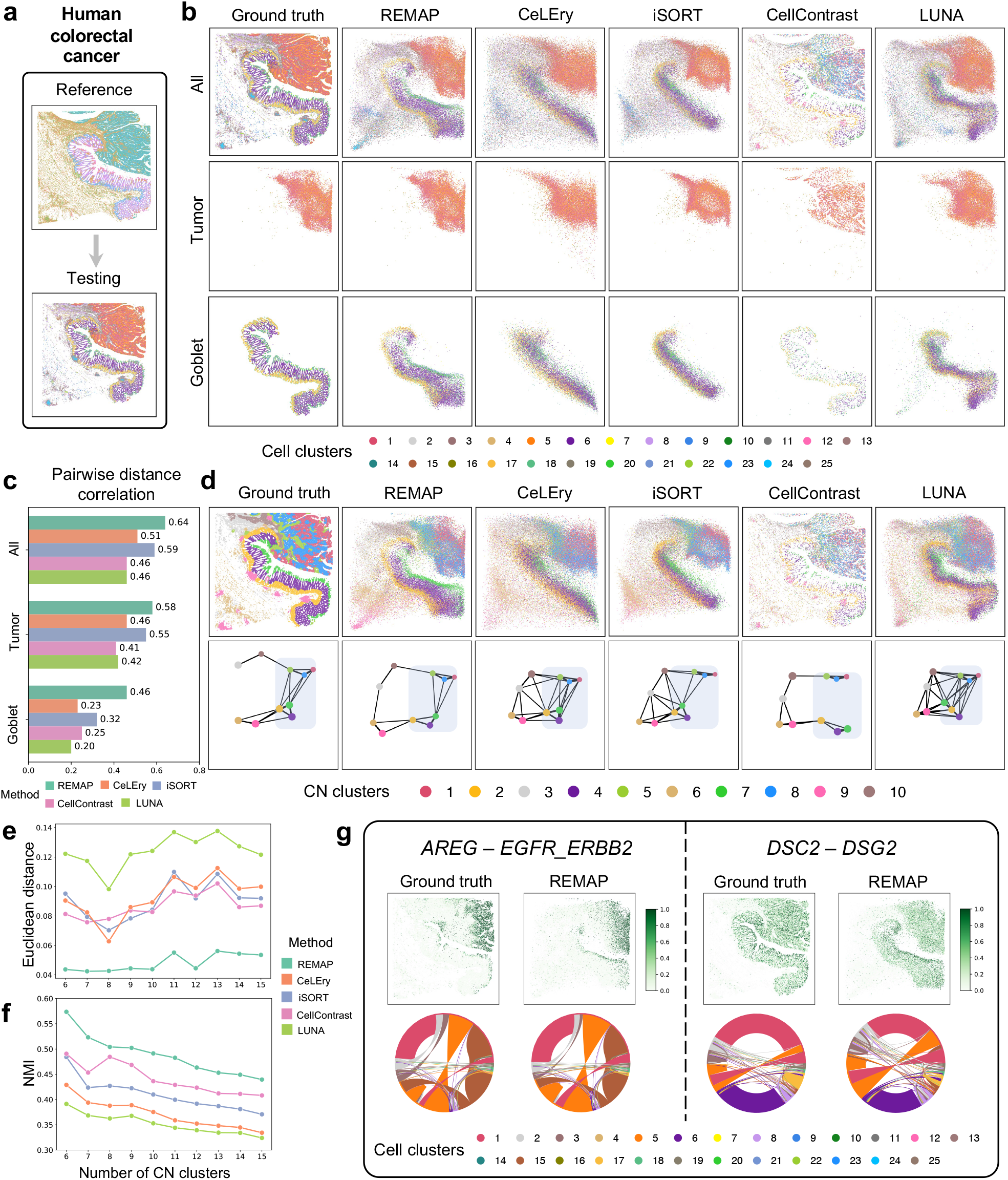
Application to human colorectal cancer data. **a**, Study design for the human colorectal cancer data benchmarking evaluation. 10x Xenium data served as the reference, and 10x Visium HD data from an adjacent slice served as the test data. **b**, Ground-truth and predicted cell locations by REMAP and other methods for all regions, tumor region, and goblet cell region. **c**, Pearson correlation between true and predicted pairwise distances for all regions, tumor region, and goblet cell region, separately. **d**, CN clusters and spatial networks constructed by REMAP and other methods. In the top panel, cells were plotted under predicted locations and colored by ground-truth CN clusters; in the bottom panel, nodes represent the centroids of CN clusters and were connected based on their spatial proximity. Edge length corresponds to the Euclidean distance between centroids, and edge width was scaled inversely to distance, such that thicker edges indicate closer proximity. Node size is proportional to the standard deviation of within-cluster pairwise distances among cells. The region of interest was highlighted using blue shaded boxes. **e**, Mean error between true and predicted pairwise Euclidean distances among CN cluster centroids, as a function of CN cluster number. **f**, NMI between true and predicted CN clusters as a function of CN cluster number. **g**, Cell-cell communication results for REMAP. For each ligand-receptor pair, the scatterplot shows the interaction strengths (i.e., 1 – *p*-value) for all cells inferred by SpatialDM, and the chord diagram represents the interaction patterns at the cell-cluster level, with node colors indicating cell types and edge colors denoting sender cell types.

CN clustering and spatial network analysis further highlighted REMAP’s ability to recover high-level architecture, as it precisely reconstructed the “double triangular” structures in these regions (**Fig. 3d, Supplementary Fig. 17**). In contrast, CeLEry, iSORT, and LUNA introduced spurious inter-cluster edges, resulting in artificially reduced inter-cluster distances relative to the ground truth. Across varying numbers of CN clusters, REMAP consistently produced the lowest pairwise cluster centroid errors (**Fig. 3e**) and the highest NMI scores (**Fig. 3f**).

Finally, we evaluated downstream applications by conducting spatially informed cell-cell communication analysis. REMAP-predicted locations uncovered complex and diverse ligand-receptor interactions (**Fig. 3g; Supplementary Fig. 18**), and the results matched those observed with true coordinates. We further analyzed a human breast cancer dataset^31^ (**Supplementary Figs. 19-21**), where high heterogeneity and irregular tumor boundaries complicate spatial reconstruction. Although REMAP could not fully recover every structural detail, it robustly reconstructed both neighborhood-level context and tissue-scale organization, and captured intricate cell-cell communication patterns. Collectively, these results demonstrate that REMAP remains accurate and robust in cancer tissues, despite its greater heterogeneity and structural irregularity, extending beyond well-organized brain tissues to highly variable pathological contexts.

### REMAP resolved neighborhood relationships and high-level spatial structures with multiple ST captures

All previous evaluations relied on full-tissue ST references. In practice, however, many tissue samples exceed the capture capacity of standard ST platforms, and researchers often profile multiple regions of interest (ROIs). Each capture represents only a subregion, and no single capture provides complete spatial information. To mimic this realistic setting, we adopted an evaluation strategy inspired by iSCALE^23^, where ROIs were manually selected from the whole tissue and treated as independent captures. To faithfully reconstruct the spatial organization of scRNA-seq data, these captures must be jointly leveraged as a combined ST reference.

We considered two scenarios using mouse brain and CRC datasets, each divided into overlapping subregions spanning the full tissue (**Fig. 4a**). Since global tissue architecture was no longer available in the ST reference, full-shape reconstruction was infeasible. Nevertheless, local neighborhood relationships were retained, which enabled the inference of pairwise relationships and spatial context for cells in scRNA-seq. We benchmarked REMAP against LUNA, the only existing method capable of handling multi-slice references. In both datasets, REMAP achieved substantially higher pairwise distance correlations (0.41 vs. 0.16 for mouse brain; 0.45 vs. 0.28 for CRC), demonstrating superior preservation of local spatial relationships (**Fig. 4b**). From predicted pairwise distance matrices, REMAP performed CN clustering and quantified spatial proximity among CN clusters by their neighborhood compositions. In mouse brain, REMAP-derived CN clusters closely resembled the ground truth, preserving spatial affinities such that clustering on the proximity matrix revealed distinct and coherent spatial niches, including cortical layers and hippocampal regions (**Fig. 4c**). LUNA failed to produce meaningful CN clusters or capture their spatial relationships, notably missing hippocampal-specific niches. Quantitatively, REMAP also achieved higher NMI scores across varying numbers of CN clusters (**Fig. 4d, Supplementary Fig. 22a**). Remarkably, for hippocampal cell types, MDS of the REMAP-predicted pairwise distance matrix recovered the overall hippocampal shape and its three subregions, despite no single ST capture spans the entire hippocampus (**Fig. 4e**). For the CRC data, REMAP’s CN clusters aligned with the ground truth, and consistently reached higher NMI scores (**Fig. 4g, Supplementary Fig. 22b**). As demonstrated by the CN spatial proximity analysis, REMAP also accurately reconstructed intra- and inter-CN spatial proximities within tumor and goblet regions (**Fig. 4f**). In contrast, LUNA produced distorted spatial proximities of the CN clusters.

**Fig. 4:**
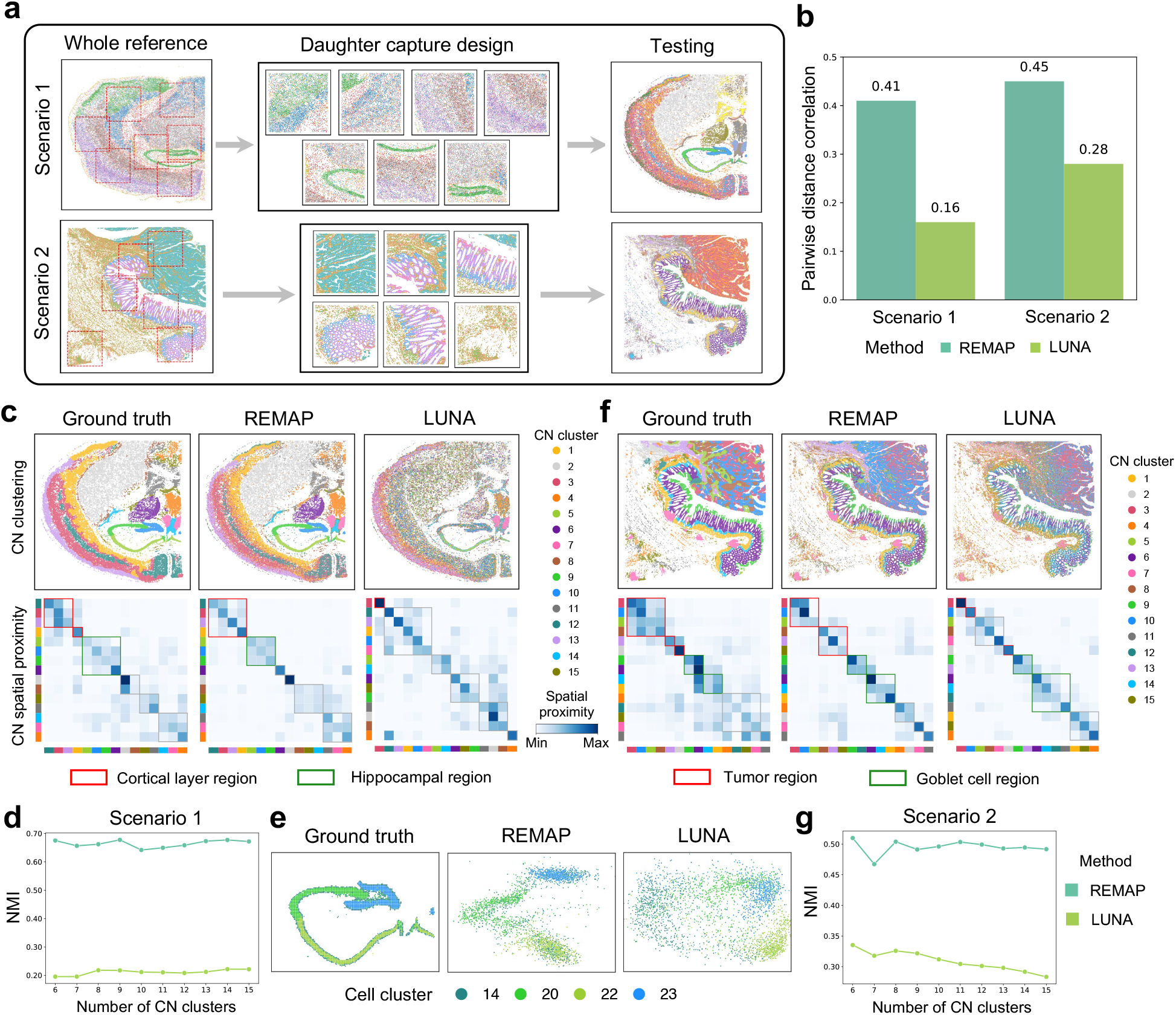
Application to multiple ST captures. **a**, Overview of the benchmarking design. Multiple equal-sized daughter captures were selected from the whole tissue and served as the reference set for model input. These designs were applied to the mouse brain data (Scenario 1) and the human colorectal cancer data (Scenario 2). **b**, Pearson correlation between true and predicted pairwise distances. **c**, CN clusters and CN spatial proximity heatmaps constructed by REMAP and LUNA in Scenario 1. In the top panel, cells were plotted under true locations and colored by predicted CN clusters; in the bottom panel, spatial proximities among CN clusters were shown as heatmaps. Clustering was performed on the predicted proximity matrices, and boxes indicated cluster memberships. Red boxes: cortical layer-specific niches; green boxes: hippocampal-specific niches; grey boxes: others. **d**, NMI between true and predicted CN clusters as a function of CN cluster number in Scenario 1. **e**, Ground-truth and recovered cell locations in the hippocampus by REMAP and LUNA. The recovered locations were manually rotated to match the orientation of the ground truth. **f**, CN clusters and spatial proximity heatmaps constructed by REMAP and LUNA in Scenario 2, similar to **c**. Red boxes: tumor-specific niches; green boxes: goblet cell-specific niches; grey boxes: others. **g**, NMI between true and predicted CN clusters as a function of CN cluster number in Scenario 2.

We further applied REMAP to a normal human gastric tissue profiled by 10x Xenium (**Supplementary Fig. 23a)**. Multiple ROIs were selected and partitioned, with half of the cells designated as the reference and the remainder, including cells outside ROIs, used as the test. Even with fragmented references, REMAP still effectively recovered the CN cluster organization and their spatial proximities, and achieved substantially higher NMI scores than LUNA (**Supplementary Fig. 23b-e**).

Finally, REMAP uniquely extends to 3D spatial reconstruction when z-axis information is available. Using two mouse whole brain datasets^32^ (147 consecutive 2D slices in Mouse 1 as reference, and 66 consecutive 2D slices in Mouse 2 as test), REMAP achieved a pairwise distance correlation of 0.73 and accurately reconstructed complex 3D structures, including DG-IMN glutamatergic neurons (**Supplementary Fig. 24**). These findings demonstrate REMAP’s capacity to generalize across diverse settings, from partial 2D ROIs to large-scale 3D reconstruction.

### REMAP uncovered spatial neighborhood heterogeneity in human multiple sclerosis

Encouraged by benchmarking results, we next applied REMAP to a human multiple sclerosis (MS) atlas^33^ to demonstrate its utility for biological discovery in real-world settings. The dataset comprised 15 samples profiled with paired Visium and snRNA-seq, spanning varying levels of demyelination and inflammation: five controls (Samples 1-5), six active MS (Samples 6-11), and four inactive MS (Samples 12-15). For each sample, we aimed to recover the spatial locations of snRNA-seq cells using the paired Visium data as a reference. Because Visium lacks single-cell resolution, we first applied iStar^34^ to enhance Visium to near single-cell resolution using histology, and then used Harmony^35^ to correct for batch effects between reference and query datasets prior to REMAP (**Fig. 5a**).

**Fig. 5:**
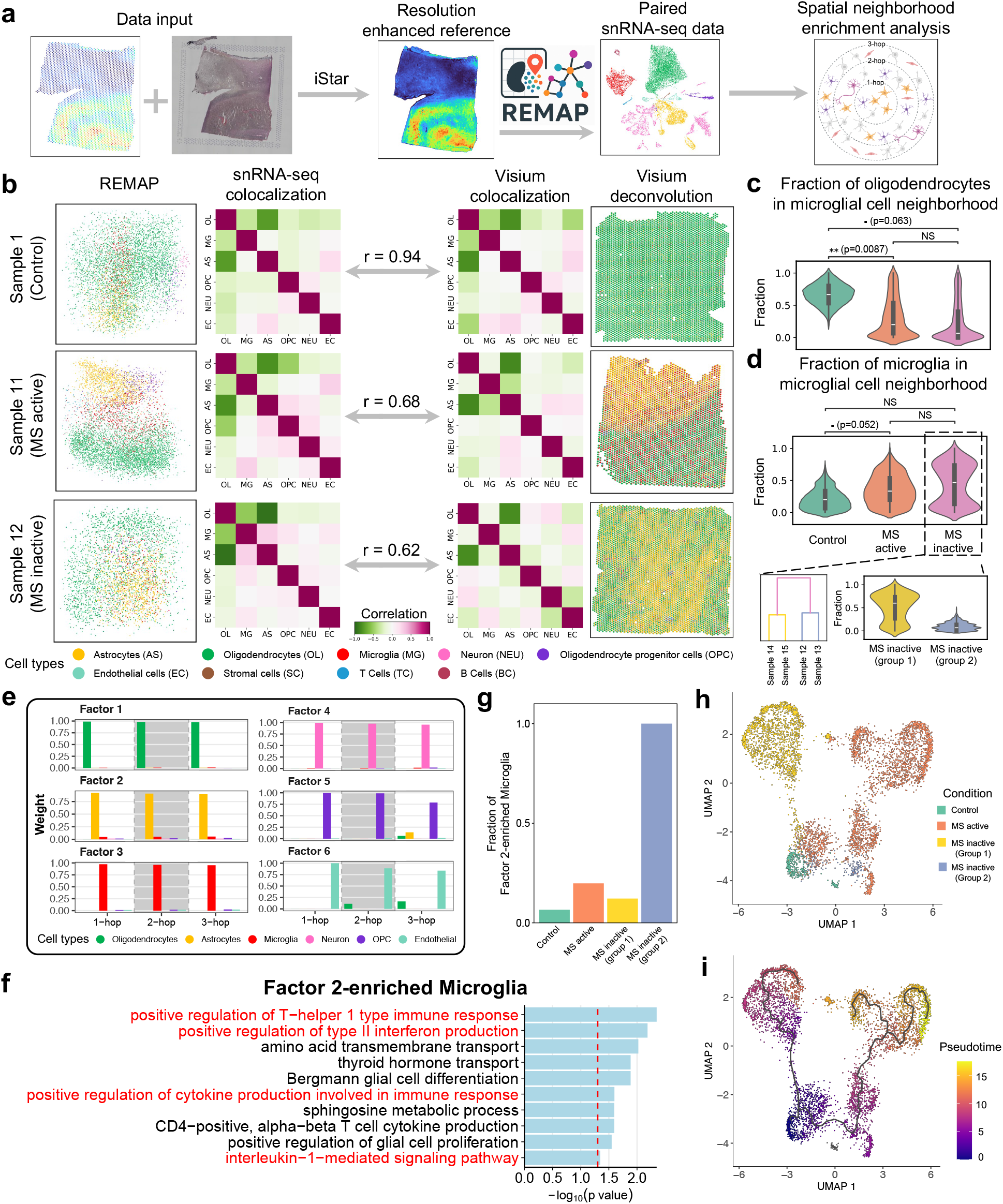
Application to MS atlas to uncover microglial spatial neighborhood variation. **a**, Overview of the study design. Visium data with paired histology images were used as input to iStar to enhance spatial resolution. The enhanced ST reference was then used as input for REMAP to infer cell locations in paired snRNA-seq. After obtaining the predicted locations, spatial neighborhood enrichment analysis was performed. **b**, Left: REMAP predicted cell locations and cell type colocalization matrices for three example samples. Middle: correlations between REMAP-inferred and Visium-derived colocalization matrices. Right: scatter pie plots displaying inferred cell type fractions from deconvolution and cell type colocalization matrices for the corresponding Visium samples. **c**, Fraction of oligodendrocytes in microglial cell neighborhoods, stratified by control, active MS, and inactive MS samples. P values were computed using a two-sided Wilcoxon rank-sum test comparing mean distributions between conditions. **d**, Top: fraction of microglia among microglial cell neighborhood, as in panel **c**. Bottom: dendrogram of inactive MS samples based on microglial cell neighborhood composition, with a violin plot further separated by the two identified subgroups. **e**, Bar plots of NMF-derived factors based on 3-hop neighborhood compositions. **f**, Top enriched pathways with adjusted p-values for upregulated genes in Factor 2-enriched microglia compared with Factor 3-enriched microglia. **g**, Fraction of Factor 2-enriched microglia among Factor 2- and Factor 3-enriched microglia in each condition. **h**,**i**, Trajectory and pseudotime inference of microglia across all conditions, shown as UMAP plots colored by conditions (**h**) and pseudotime (**i**).

REMAP reliably recovered spatial locations for snRNA-seq cells across all 15 samples (**Fig. 5b, Supplementary Fig. 25**). We validated these predictions by comparing cell type colocalization patterns inferred from REMAP with those derived from Visium deconvolution using cell2location^36^. For snRNA-seq, the cell type colocalization matrix was computed by identifying the fractions of each cell type among the 15 nearest neighbors per cell and computing Pearson correlations between the cell type fractions across all cells. Visium’s colocalization matrix was obtained by calculating the Pearson correlations between cell type proportions across all deconvoluted spots. The similarity between the two colocalization matrices was quantified by vectorizing each matrix and computing the Pearson correlation between the two vectors. We focused our analyses on six major cell types: astrocytes, oligodendrocytes, microglia, neurons, oligodendrocyte progenitor cells, and endothelial cells. Overall, REMAP predictions showed strong concordance with Visium-derived colocalization patterns, with discrepancies likely resulting from differences in cell type composition between Visium and snRNA-seq data, particularly in MS disease samples (**Supplementary Fig. 26**).

We next focused on microglia, the primary immune cells of the central nervous system and key drivers of lesion progression in MS^37^. For each microglial cell, we identified its 15 nearest spatial neighbors and computed the cell type composition within its neighborhood. Across disease states, oligodendrocytes and microglia showed the most striking differences (**Fig. 5c,d**). In controls, microglia were frequently surrounded by oligodendrocytes, whereas in MS, they increasingly colocalized, indicating increased microglia-microglia interactions^38^. Notably, the inactive MS samples exhibited a bimodal distribution in neighborhood composition (**Fig. 5d**), consistent with prior snRNA-seq studies reporting that patient-level variability can exceed lesion-level signatures in shaping MS gene expression profiles^39^.

To further explore this heterogeneity, we performed hierarchical clustering on the four inactive MS samples based on their mean neighborhood cell type composition. The resulting dendrogram revealed two distinct subgroups: Samples 14 and 15 formed inactive group 1, and Samples 12 and 13 formed inactive group 2. Violin plots showed that the microglial neighborhoods in inactive group 2 resembled controls, with minimal self-colocalization. In contrast, inactive group 1 mirrored the features of active MS, with stronger microglial self-colocalization (**Fig. 5d**). Therefore, we retained this stratification of inactive samples for subsequent analyses.

To obtain a global view of tissue microenvironments, we defined three tiers of spatial neighbors for each cell: 1-hop (top 15), 2-hop (top 16-50), and 3-hop (top 51-100). For each cell, cell type fractions were computed within each hop, yielding a multi-scale neighborhood representation. Aggregating across all cells and samples, we applied non-negative matrix factorization (NMF) to this cell type fraction matrix, specifying six factors to match the number of primary cell types. The resulting factors (**Fig. 5e**) captured distinct neighborhood contexts, each enriched for a cell type. For instance, Factor 3 corresponded to a microglia-enriched context, while Factor 2 reflected astrocyte-enriched neighborhoods. Although most microglia loaded strongly on Factor 3, a subset showed notable contributions from Factor 2, suggesting localized microglia-astrocyte colocalization. We therefore defined two subpopulations: Factor 3-enriched microglia and Factor 2-enriched microglia. Differential expression (DE) analysis between these two subpopulations (**Supplementary Table 2**) revealed that Factor 2-enriched microglia expressed genes consistent with a transitioning and more reactive state. These included *CHIT1*, a biomarker of early microglial activation and disability progression in MS^40^, and *SIGLEC1*, a marker of active neuroinflammation^41^. Gene Ontology (GO) enrichment analysis further confirmed enrichment for pathways related to interferon signaling, immune activation, cytokine production, and interleukin mediation (**Fig. 5f; Supplementary Table 3**), supporting a proinflammatory microglial phenotype and reinforcing the hypothesis of enhanced microglia-astrocytes crosstalk in this rare subpopulation of microglia^42^.

Next, we quantified the proportion of Factor 2-enriched microglia among all Factor 2- and Factor 3-enriched microglia across conditions (**Fig. 5g**). These cells were most prevalent in inactive group 2. Trajectory inference and pseudotime analysis with Monocle 3^43^ revealed that microglia from inactive group 2 possessed an intermediate state between control samples and active MS, suggesting a transition from homeostatic toward a more reactive phenotype (**Fig. 5h,i**).

### REMAP revealed conserved spatial subtypes of cancer-associated fibroblasts across human cancers

Having established REMAP’s real-world applicability in brain studies, we extended its use to multiple cancer types to demonstrate its broad utility in identifying spatial subtypes. We analyzed four large human cancer datasets with full tissue coverage: cervical and ovarian cancers profiled by Xenium 5K, melanoma profiled by MERSCOPE, and lung cancer profiled by CosMx. We first assessed REMAP for within-subject location prediction, where cervical, ovarian cancers, and melanoma datasets were randomly divided into two halves, with one half as the reference and the other as the testing set. For the lung cancer dataset, adjacent tissue sections were used as reference and testing data. REMAP accurately reconstructed tissue shapes across all cancer types (**Fig. 6a-d**). Among existing tools, only CeLEry could process all of these large-scale datasets on an NVIDIA A100 GPU with 80 GB memory, but it produced less refined spatial architectures (**Supplementary Fig. 27**). Quantitatively, REMAP consistently achieved higher pairwise distance correlations than other methods (**Fig. 6e**).

**Fig. 6:**
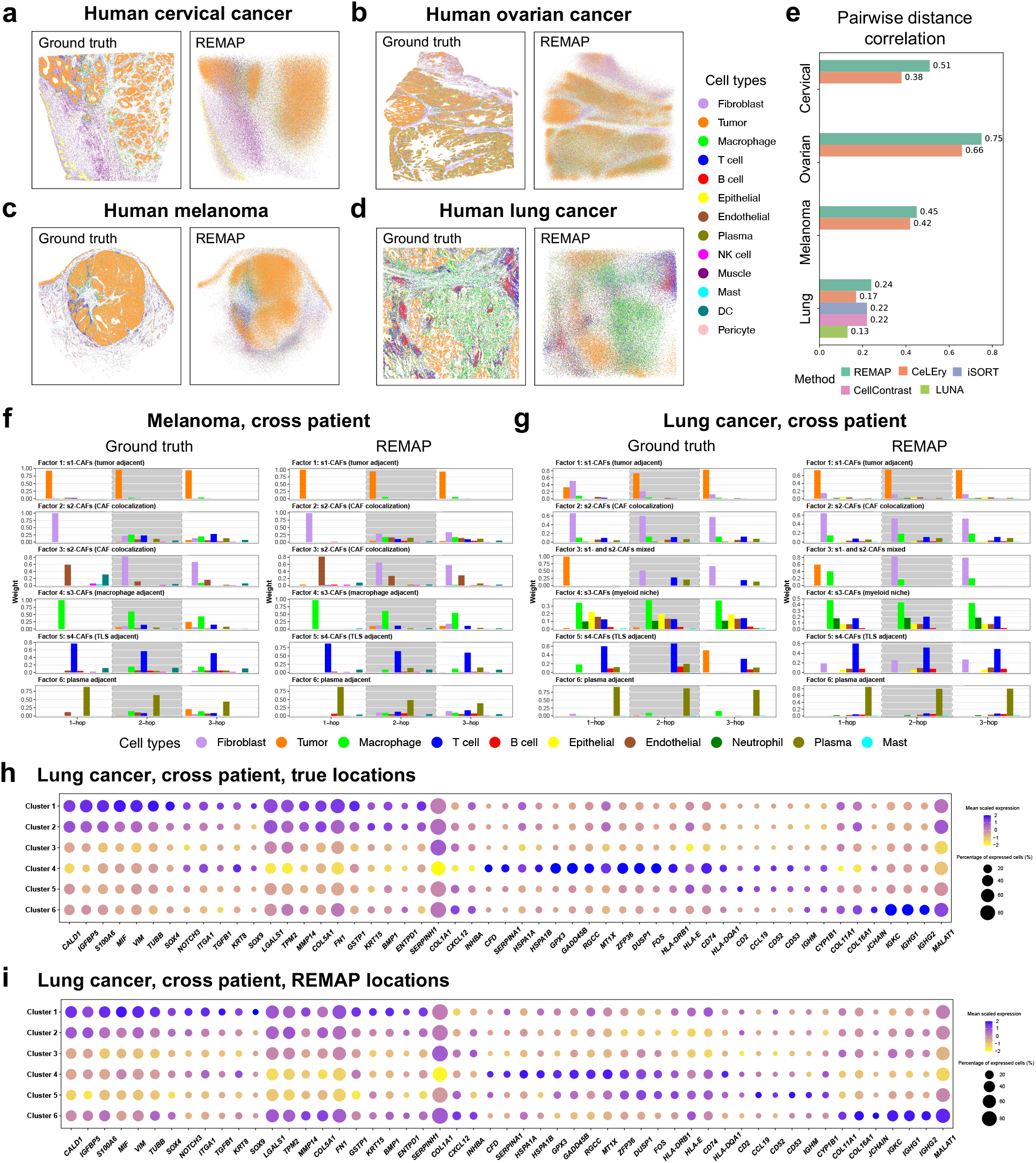
Application to multiple human cancers to identify conserved spatial subtypes of CAFs. **a-d**, Ground-truth and REMAP-predicted cell locations for within-subject location prediction across multiple human cancers, including cervical cancer (**a**), ovarian cancer (**b**), melanoma (**c**), and lung cancer (**d**). **e**, Pearson correlation between true and predicted pairwise distances. For cervical cancer, ovarian cancer, and melanoma datasets, besides REMAP, only CeLEry could process the full datasets. **f-g**, Bar plots of NMF-derived factors based on 3-hop neighborhood compositions of CAFs for cross-subject location prediction using ground-truth and REMAP-predicted locations in melanoma (**f**) and lung cancer (**g**). **h-i**, Bubble plots showing the expression of differentially expressed genes across NMF-derived CAF clusters for cross-subject location prediction in lung cancer, using ground-truth (**h**) and REMAP-predicted locations (**i**).

Next, we tested REMAP’s performance for an even more challenging situation that involves cross-subject prediction, in which reference and testing samples were obtained from different cancer patients. Since tissue morphology varies substantially across individuals, direct reconstruction of tissue shape was infeasible; however, it remained possible to uncover cellular neighborhood compositions and spatial niches. Cancer-associated fibroblasts (CAFs), a multifaceted cell population known to shape the tumor microenvironment and influence therapeutic responses^44^, exhibit conserved spatial subtypes with unique neighborhood profiles across cancer types^45^. We therefore focused on the CAF neighborhood organization. After location prediction, each CAF cell was treated as a centroid, and concentric circular regions with radii of 40 *μm*, 80 *μm*, and 120 *μm* were defined to capture its three-tier neighborhoods. Cells located within these regions were identified as their three-tier neighbors, and their cell type compositions were computed. Across all CAFs, NMF was then applied to the resulting composition matrix.

Applying this framework to melanoma, we used a second MERSCOPE dataset from a different patient as the testing sample. REMAP-derived factors closely recapitulated the ground truth and revealed distinct CAF neighborhood profiles, aligning with the s1-to s4-CAF subtypes reported by Liu et al.^45^ (**Fig. 6f**). Factor 1 corresponded to s1-CAFs, which localized within tumor regions and, importantly, have been shown to possess prognostic value in human tumors^45^. In contrast, CAFs enriched for Factors 2 and 3 showed self-colocalization patterns, recapitulating s2-CAFs. Factor 4- and Factor 5-enriched CAFs, representing s3-CAFs and s4-CAFs, were predominantly adjacent to macrophages and tertiary lymphoid structures (TLSs), respectively. Factor 6-enriched CAFs were proximal to plasma cells. In lung cancer, using a second CosMx slice from a different patient as testing data, REMAP again successfully recovered diverse spatial patterns of CAFs (**Fig. 6g**), demonstrating its robustness in decoding complex and subtle microenvironment variability within the same cell type.

To dissect the transcriptional programs underlying these spatial subtypes of CAFs, we conducted K-Means clustering on the NMF-derived cell-by-factor matrix and performed DE analysis for each cluster. REMAP revealed distinct and biologically interpretable expression profiles that reliably matched the ground truth in both melanoma and lung cancer (**Fig. 6h,i**; **Supplementary Fig. 28**). For example, in lung cancer, Cluster 1 and 2 exhibited elevated expressions on genes associated with tumor progression (*CALD1*^46^, *LGALS1*^47^, *MMP14*^48^), immune suppression (*TGFB1, MYH11*)^49^, collagen production (*NOTCH3*^50^, *SERPINH1*^51^), and ECM remodeling (*FN1*^52^, *BMP1*^53^), promoting tumor invasion and consistent with s1-CAF identity. Cluster 4 displayed increased expression of stress response genes (*DUSP1, GADD45B, HSPA1A, HSPA1B*)^54^, reflecting s3-CAF features. In contrast, Cluster 5 showed enriched expression of immune-modulatory cytokines (*CCL19, CXCL9*) and antigen presentation genes (*CD74*, HLA class II molecules)^55^, characteristics of immunoactive s4-CAF (**Fig. 6h,i; Supplementary Table 4**). These results validated that REMAP-derived CAF spatial subtypes corresponded to transcriptionally distinct and functionally meaningful states.

We also repeated the CAF neighborhood analysis framework on four within-subject predictions that involve cervical cancer, ovarian cancer, melanoma, and lung cancer. REMAP consistently reconstructed all CAF spatial subtypes (**Supplementary Fig. 29**). Importantly, these neighborhood patterns, especially tumor adjacent s1-CAFs and self-colocalized s2-CAFs, were recurrently detected across multiple cancer types, highlighting REMAP’s ability to capture conserved CAF spatial subtypes across diverse cancer types. We further applied the same workflow to a prostate cancer MERSCOPE dataset (**Supplementary Fig. 30**). REMAP again reconstructed tissue structures under within-subject prediction and identified distinct and meaningful CAF spatial subtypes under cross-subject prediction. Together, these findings demonstrated REMAP’s capacity to robustly decode the spatial organizational heterogeneity of CAFs and classify them into distinct spatial subtypes, directly from dissociated single-cell data across diverse human cancers.

## Discussion

In this paper, we have presented REMAP, a deep learning method that leverages single or multiple ST references to reconstruct multi-scale spatial information for cells in scRNA-seq. By integrating gene expression with neighborhood context encoded through gene-gene covariance, REMAP consistently outperformed existing approaches across tissues and platforms. When global spatial information was available, REMAP accurately reconstructed tissue-level architectures, such as the hippocampal structure in the mouse brain and tumors in human cancers. Besides, REMAP uniquely extended to 3D spatial reconstruction when the z-axis information was available in reference ST. In scenarios where tissue architecture is fragmented across multiple ST captures, REMAP’s multi-slice training framework inferred CN clusters and their spatial proximities to characterize global tissue organization. REMAP is computationally fast (**Supplementary Fig. 31**) and scales efficiently to large datasets containing hundreds of thousands of cells on a single NVIDIA A100 GPU with 80 GB memory.

Beyond benchmarking, we demonstrated REMAP’s potential for biological discovery in neuroinflammatory and cancer contexts. In a human MS atlas, REMAP uncovered differences in microglial neighborhoods between control and MS and revealed heterogeneity among inactive MS samples. Strikingly, REMAP identified a rare microglial subpopulation colocalizing with astrocytes, marked by a transitioning, activated, immune-responsive state enriched for proinflammatory pathways. These insights could not be obtained from Visium data, due to its limited spatial resolution and imprecise annotations, or from snRNA-seq data alone, which lacks spatial information. Across multiple human cancer types, including lung, melanoma, cervical, ovarian, and prostate, REMAP uncovered conserved spatial subtypes of CAFs characterized by distinct neighborhood compositions and transcriptional profiles, robust across both within- and cross-subject predictions. These CAF subtypes occupied specialized spatial niches, ranging from tumor proximal to TLS adjacent, suggesting diverse functional roles and interaction patterns within the same cell type. Together, these findings establish REMAP as a generalizable framework for identifying spatial subtypes and decoding tissue organization and disease biology directly from snRNA-seq data.

While REMAP has achieved strong performance in recovering spatial coordinates, this task remains particularly challenging in cancer tissues, which exhibit complex and fine-grained microenvironments. In such cases, REMAP is limited in its ability to fully recover tissue shapes. Although REMAP employs an iterative training strategy to mitigate this limitation, the estimated neighborhood gene-gene covariance inevitably lacks the high resolution available in reference ST data. Nevertheless, perfect recovery of tissue shapes may not be necessary for many downstream applications. In analyses such as cell-cell communication, pairwise spatial distances among cells are typically used to constrain inference and ensure reliability of detected communication signals. Because REMAP excels in preserving pairwise spatial relationships, it provides a robust foundation for such tasks, even when fine-grained tissue structures cannot be fully recovered.

We further highlighted REMAP’s unique ability to reconstruct CN spatial networks and performed NMF-based spatial neighborhood enrichment analysis directly from scRNA-seq. CN spatial networks provide a high-level, structured, and interpretable characterization of tissue organization by summarizing how groups of cells, defined by shared expression and spatial context, colocalize and interact within the tissue. This framework is particularly valuable in population-scale scRNA-seq studies, where samples from different conditions (e.g., health vs. disease) are collected and compared. By comparing CN networks across individuals, researchers can identify conserved spatial modules, quantify inter-individual variation, and detect condition-specific alterations in tissue organization. Additionally, applying NMF to multi-scale neighborhood composition matrices enabled the discovery of shared tissue microenvironments across samples. Comparable analysis of factor loadings revealed phenotype-associated spatial structures, while factor patterns dissected heterogeneous spatial configurations within a given cell type and uncovered previously unrecognized spatially informed subpopulations. Importantly, both the CN network and the NMF-based approaches are robust to minor prediction errors in absolute cell positions. As long as neighborhood compositions are reliably predicted, these methods can generate meaningful insights.

Together, these advantages highlight REMAP’s ability to uncover functional tissue units that persist across biological states and to detect rewiring events in disease and inflammation. As large-scale initiatives such as the Human Cell Atlas^24,56,57^ increasingly adopt paired ST and scRNA-seq designs, we anticipate REMAP will have broad applications, particularly in population-level single-cell studies.

## Data availability

(1) 10x Visium HD mouse brain data (https://www.10xgenomics.com/datasets/visium-hd-cytassist-gene-expression-libraries-of-mouse-brain-he); (2) 10x Xenium mouse brain data (https://www.10xgenomics.com/datasets/fresh-frozen-mouse-brain-for-xenium-explorer-demo-1-standard); (3) Processed MERFISH and snRNA-seq human cerebral cortex data (https://doi.org/10.5281/zenodo.14422018); (4) 10x Visium HD and Xenium human colorectal cancer data (https://www.10xgenomics.com/platforms/visium/product-family/dataset-human-crc); (5) Visium and snRNA-seq human multiple sclerosis data reported in Lerma-Martin et al.^33^ (GEO GSE279183). (6) 10x Xenium 5K human cervical cancer data (https://www.10xgenomics.com/datasets/xenium-prime-ffpe-human-cervical-cancer); (7) 10x Xenium 5K human ovarian cancer data (https://www.10xgenomics.com/datasets/xenium-prime-ffpe-human-ovarian-cancer); (8) MERSCOPE human melanoma data (https://vizgen.com/applications/tumor-profiling, Melanoma 1 and Melanoma 2). (9) CosMx human non-small cell lung cancer data (https://nanostring.com/products/cosmx-spatial-molecular-imager/ffpe-dataset/nsclc-ffpe-dataset/). (10) MERSCOPE human prostate cancer data (https://vizgen.com/applications/tumor-profiling, Prostate cancer 1 and Prostate cancer 2). (11) Processed MERFISH and scRNA-seq mouse primary motor cortex data (https://dp-lab-data-public.s3.amazonaws.com/ENVI/st_data.h5ad, https://dp-lab-data-public.s3.amazonaws.com/ENVI/sc_data.h5ad); (12) 10x Xenium human breast cancer data (https://www.10xgenomics.com/products/xenium-in-situ/preview-dataset-human-breast); (13) 10x Xenium human gastric data (https://doi.org/10.5281/zenodo.15164978); (14) MERFISH whole mouse brain data (https://alleninstitute.github.io/abc_atlas_access/descriptions/Zhuang-ABCA-1.html, https://alleninstitute.github.io/abc_atlas_access/descriptions/Zhuang-ABCA-2.html). Details of the datasets analyzed in this paper are described in **Supplementary Table 1**.

## Code availability

REMAP was implemented in Python and is available on GitHub at https://github.com/ShunzhouJiang/REMAP.

## Acknowledgments

M.L. was partly supported by the following NIH grants R01HG013185, R01LM014592, U19NS135582, R01HL171595, and U01CA294518. X.Q. was partially supported by the NIH Pathway to Independence Award R00NS135123. C.A.W. was supported by R01 NS035129 and R01 NS032457 from the NINDS, the Allen Discovery Center program, a Paul G. Allen Frontiers Group advised program of the Paul G. Allen Family Foundation, and John Templeton Foundation research grant (#62587). C.A.W. is an Investigator of the Howard Hughes Medical Institute. J.J. was partly supported by NIH grants R00HG012223 and R35GM157133. T.H.H. was partly supported by NIH grants R01CA276690 and U01CA294518, DOD grant CA190578, the Eric and Wendy Schmidt Foundation’s AI Innovation Award through the Mayo Clinic Foundation, and the Torrey Coast Foundation.

## Author contributions

This study was conceived of and led by M.L. S.J. designed the model and algorithm with input from M.L., implemented the software, and led data analyses. K.C. performed the cortical depth analysis. Z.C., R.X., and J.J. provided input on REMAP algorithms. K.J. and D.L. helped with figure design. Y.L. and L.W. annotated cell types and provided suggestions for the CAF applications. T.H.H. provided the gastric cancer Xenium data. C.A.W. and X.Q. provided the human fetal cortex MERFISH data and helped with interpretation and analysis. S.J. and M.L. wrote the paper with feedback from other co-authors.

## Methods

### Data preprocessing

REMAP takes as input one or more ST reference dataset(s) and a target scRNA-seq (or snRNA-seq) dataset, for which cell locations are unknown. Depending on the ST platform, spatial coordinates may be represented in either 2D or 3D. We first subset the common genes between ST and scRNA-seq datasets. For each cell in ST or scRNA-seq, the raw gene counts are normalized as

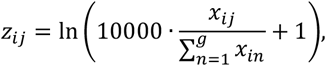

where *x*_*ij*_ is the raw count of gene *j* in cell *i*, and *g* is the number of common genes. Then gene-level z-score standardization is applied,

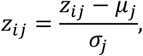

where *μ*_*j*_ and *σ*_*j*_ are the mean and standard deviation of gene *j* expression across all cells. For spatial references, cell locations in each ST slice are scaled relative to the capture area and normalized to the range of [0, 1], facilitating comparisons across ST slices. To ensure comparability across datasets, we recommend removing gene expression batch effects between ST and scRNA-seq data before running REMAP. State-of-the-art methods such as Harmony^35^ and CarDEC^25^ can be applied for this purpose.

### Cellular neighborhood representation

To represent the local neighborhood context, REMAP uses the gene-gene covariance matrix between neighboring cells. We first select the top *m* highly variable genes for gene-gene covariance calculation. For a given ST slice with *N*_1_ cells, and for each cell *i*, we identify its *k* nearest spatial neighboring cells and extract their gene expression matrix *E*_*i*_ ∈ *R*^*k*×*m*^. Next, the shifted gene-gene covariance matrix ∑_*i*_ ∈ *R*^*m*×*m*^ is computed as:

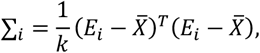

where 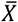 is the mean expression across all cells. By default, we select *k* = 100 for single-cell resolution ST data, and *m* = min(100, *g*), where *g* is the number of shared genes between ST and scRNA-seq datasets. Across all cells, this procedure yields a tensor of dimension an *N*_1_ × *m* × *m*. Since covariance matrices are symmetric, we extract and flatten their lower triangular parts into 1D vectors. To reduce dimensionality, principal component analysis (PCA) is applied with 98% of the variance retained by default, yielding dimension-reduced vectors 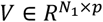, where *p* is the number of retained components. For scRNA-seq data, as cell locations are unknown, neighboring gene-gene covariance matrices cannot be directly computed. To address this, we use ENVI^21^, a conditional variational autoencoder-based integration tool for ST and scRNA-seq, to initialize estimates of neighborhood gene-gene covariance vectors 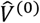 for scRNA-seq cells.

### Location prediction model with a single ST reference

To recover spatial locations for scRNA-seq cells, REMAP integrates gene expression features with neighboring gene-gene covariance features and iteratively updates the neighboring gene-gene covariance and cell location predictions. At initialization, neighboring gene-gene covariance vectors are computed from ground truth locations in ST data but inferred using ENVI for scRNA-seq, which results in a mismatch in accuracy. While ST-derived covariance captures fine-grained tissue microenvironment, this level of granularity is not directly attainable in scRNA-seq due to the absence of spatial information. Thus, directly applying a model trained on high-resolution ST features to scRNA-seq can therefore degrade performance. To address this, REMAP employs an iterative framework.

1. Location predictor: A deep neural network is trained using concatenated features (gene expression plus dimension-reduced gene-gene covariance vectors) to predict spatial coordinates. The loss is defined by mean squared error (MSE):

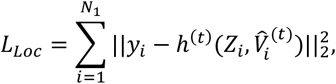

where *y*_*i*_ is the true coordinates of cell *i, Z*_*i*_ is its gene expression, and 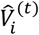 is the estimated covariance vector at iteration *t*, and *h*^(*t*)^ denotes the *t*^*th*^ round of location prediction neural network. At *t* = 0, 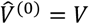 for ST cells.
2. Covariance predictor: Using the locations predicted in Step 1, a separate neural network is trained to refine covariance estimates. The predicted coordinates are scaled as

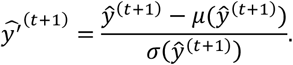

We infer 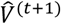 by minimizing the following MSE loss function:

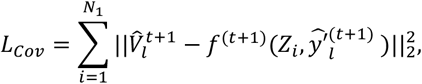

where *f*^(*t* + 1)^ denotes the (*t* + 1)^*th*^ round covariance prediction neural network.
3. Iteration: The updated covariance vector 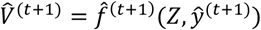 is then fed back into the location predictor for the next round. The previous two steps are iteratively trained, with the output of each iteration serving as input features for the next round.

REMAP allows users to specify the number of hidden layers and nodes. By default, both the location and covariance prediction networks use three fully connected layers (400, 200, 100 nodes). Each layer applies ReLU activation with a dropout probability of 0.05. Output layer dimensions match prediction target: *p* for covariance prediction, 2 for 2D coordinates, and 3 for 3D coordinates. The location predictor’s output uses a Sigmoid activation function to scale the output to the [0, 1] range. During training, both networks are optimized with stochastic gradient descent using the Adam optimizer. The network architectures remain consistent across iterations.

### Location prediction model with multiple ST references

ST slices from the same tissue often differ in orientation or shape, making their absolute coordinates not directly comparable and difficult to align. However, pairwise distances between cells remain comparable across slices. Thus, when multiple ST references are available, REMAP predicts pairwise distances rather than absolute coordinates.

Let *R* denote the number of reference ST slices. Because a test cell may be more similar to some reference slices than others, we assign slice-specific weights based on transcriptomic similarity. A fixed number of *K* cells are sampled from each ST slice and combined with the test data. For each test cell *i*, we identify 0.1 × *K* nearest neighbors based on expression similarity. The distribution of neighbor sources across slices defines the weight vector *w*_*i*·_, which reflects the relative similarity of the cell *i* to each slice. The overall slice-specific weight for the slice *r* is obtained by averaging across all test cells:

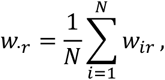

where *w*_*ir*_ is the contribution of the slice *r* to cell *i*, and *N* is the total number of test cells considered.

To predict pairwise distances, REMAP constructs a neural network that takes two cells from the same ST slice at a time. For each cell pair, their features (gene expression and neighboring gene-gene covariance vectors) are concatenated and passed independently through a fully connected network. The absolute difference between the two embeddings of the cell pair is then computed and further processed through two additional fully connected networks with ReLU activation and dropout (*p* = 0.05) to learn a joint representation. The network outputs a single scalar representing the predicted pairwise distance between the two cells. A Sigmoid activation function is applied to the output, followed by scaling with 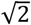, so that the prediction lies within the normalized Euclidean distance range 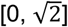. The neural network is trained by minimizing the following loss function:

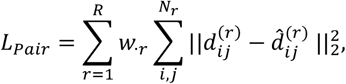

where *w*_·*r*_ denotes the mean weight for the slice *r*, and 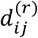 and 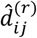 represent the true and predicted pairwise distances between cells *i* and *j* in the slice *r*, respectively. After training, the model can be applied to scRNA-seq data to generate a predicted pairwise distance matrix across all test cells. For visualization, Multidimensional Scaling (MDS) can be applied to the predicted distance matrix to embed cells into 2D or 3D spatial coordinates.

In REMAP, each training observation corresponds to a cell pair, and the total number of observations is the number of cell pairs across all ST slices. For single-cell resolution ST datasets, this number can be prohibitively large. To address this, we implement a grid-based downsampling strategy to reduce the computational burden while preserving spatial diversity. For each ST slice, the spatial capture area is divided into a *d* × *d* grid. From each grid, we select the cell with the highest UMI count, forming a representative subset of cells that provides balanced tissue coverage. Pairwise training samples are then constructed using only these selected cells. By default, *d* = 50.

During the inference phase, predicting the full pairwise distance matrix for all scRNA-seq cells can be computationally expensive for large datasets. However, when the primary goal is to identify spatial neighbors and infer CN clusters and spatial niches, constructing the complete pairwise distance matrix is often unnecessary. Therefore, REMAP introduces an optional filtering step. For each cell, the top 10% feature-based neighbors are first identified and treated as spatial neighbor candidates. After model training, pairwise distances are predicted only for those candidate pairs, and spatial neighbors are inferred among these candidates. This filtering reduces unnecessary computations for cell pairs that are unlikely to be neighbors and ensures that the running time scales linearly with the number of cells.

### Cellular Neighborhood (CN) clustering and CN spatial network construction

To define cellular neighborhoods, we first identify each cell’s top *K* nearest spatial neighbors and compute its neighborhood cell type composition. Based on this composition matrix, cells are grouped into CN clusters using K-Means clustering. By default, *K* = 100. CN spatial networks are constructed as follows. For each CN cluster, the centroid of its member cells is computed. Pairwise distances between centroids are then calculated across all CN cluster pairs. In the network visualization, each CN cluster is represented as a node placed at its centroid. Edges are drawn between spatially proximal CN clusters, with edge thickness proportional to spatial proximity (i.e., inverse of the centroid distance). Node size is proportional to the standard deviation of within-cluster pairwise distances among cells, such that clusters with more dispersed member cells appear as larger nodes. For the multi-capture reference analysis, since absolute locations are not observed, we quantify the spatial proximities among CN clusters as follows. For each cell, its top *K*_*CN*_ spatial neighbors are identified based on predicted pairwise distances, and the CN cluster composition within the neighbors is computed. Averaging these fractions across all cells within one CN cluster yields the CN spatial proximity heatmap. By default, *K*_*CN*_ = 2,000.

### Spatially informed cell-cell communication analysis

We performed spatially informed cell-cell communication analysis on both ground truth and REMAP-predicted locations using SpatialDM^26^. SpatialDM first computes the global Moran’s R statistic to detect spatially co-expressed ligand-receptor pairs. A pair is considered significant if its FDR-adjusted *p*-value <0.1. For each significant ligand-receptor pair, local Moran’s R is applied to detect locally interacting cells, with significance defined at *p*-value <0.1. Following the visualization strategies in the original SpatialDM paper, we generated interaction intensity maps for each ligand-receptor pair, where scatterplots display cells colored by interaction strength (defined as 1 – *p*-value), with darker colors indicating stronger involvement for a given ligand-receptor. In addition, chord diagrams were used to summarize cell type-level communication patterns, where node colors represent cell types and edge colors indicate the sender cell types. For analyses conducted in this paper, ligand-receptor pair lists were extracted from CellChatDB^58^.

### Differential expression (DE) and Gene Ontology (GO) enrichment analysis

For the human MS data, we performed DE analysis between Factor 2-enriched and Factor 3-enriched microglia using the *sc*.*tl*.*rank_genes_groups* function in Scanpy^59^ with a t-test. A gene was considered significantly upregulated if its Benjamini-Hochberg (BH) adjusted *p*-value <0.05 and log fold change>0. GO enrichment analysis of the DE genes was conducted using the *enrichGO* function in R package clusterProfiler^60^. Pathways with a BH-adjusted *p*-value <0.05 were deemed significantly enriched. To reduce redundancy among enriched terms, we applied the *simplify* function in clusterProfiler.

### Pseudotime inference

For the human MS data, microglia cells were extracted from all samples and analyzed with Monocole 3^43^ for trajectory inference. The raw count matrix was preprocessed by *preprocess_cds* function with PCA (num_dim=800). Trajectories were then constructed following the default Monocole 3 workflow, specifically with the *learn_graph* function, and pseudotime was estimated using the *order_cells* function. The control group was designated as the root for pseudotime assignment.

## References

1. Crosetto, N., Bienko, M. & van Oudenaarden, A. Spatially resolved transcriptomics and beyond. Nat Rev Genet 16, 57–66 (2015). 10.1038/nrg3832

2. Chen, K. H., Boettiger, A. N., Moffitt, J. R., Wang, S. & Zhuang, X. RNA imaging. Spatially resolved, highly multiplexed RNA profiling in single cells. Science 348, aaa6090 (2015). 10.1126/science.aaa6090

3. Burgess, D. J. Spatial transcriptomics coming of age. Nat Rev Genet 20, 317 (2019). 10.1038/s41576-019-0129-z

4. Asp, M., Bergenstrahle, J. & Lundeberg, J. Spatially Resolved Transcriptomes-Next Generation Tools for Tissue Exploration. Bioessays 42, e1900221 (2020). 10.1002/bies.201900221

5. Rao, A., Barkley, D., Franca, G. S. & Yanai, I. Exploring tissue architecture using spatial transcriptomics. Nature 596, 211–220 (2021). 10.1038/s41586-021-03634-9

6. Moses, L. & Pachter, L. Museum of spatial transcriptomics. Nat Methods 19, 534–546 (2022). 10.1038/s41592-022-01409-2

7. Hu, J. et al. Statistical and machine learning methods for spatially resolved transcriptomics with histology. Comput Struct Biotechnol J 19, 3829–3841 (2021). 10.1016/j.csbj.2021.06.052

8. Longo, S. K., Guo, M. G., Ji, A. L. & Khavari, P. A. Integrating single-cell and spatial transcriptomics to elucidate intercellular tissue dynamics. Nat Rev Genet 22, 627–644 (2021). 10.1038/s41576-021-00370-8

9. Zeng, Z., Li, Y., Li, Y. & Luo, Y. Statistical and machine learning methods for spatially resolved transcriptomics data analysis. Genome Biol 23, 83 (2022). 10.1186/s13059-022-02653-7

10. Coleman, K., Schroeder, A. & Li, M. Unlocking the power of spatial omics with AI. Nat Methods 21, 1378–1381 (2024). 10.1038/s41592-024-02363-x

11. Guo, B. et al. Integrating Spatially-Resolved Transcriptomics Data Across Tissues and Individuals: Challenges and Opportunities. Small Methods 9, e2401194 (2025). 10.1002/smtd.202401194

12. Biancalani, T. et al. Deep learning and alignment of spatially resolved single-cell transcriptomes with Tangram. Nat Methods 18, 1352–1362 (2021). 10.1038/s41592-021-01264-7

13. Cang, Z. & Nie, Q. Inferring spatial and signaling relationships between cells from single cell transcriptomic data. Nat Commun 11, 2084 (2020). 10.1038/s41467-020-15968-5

14. Nitzan, M., Karaiskos, N., Friedman, N. & Rajewsky, N. Gene expression cartography. Nature 576, 132–137 (2019). 10.1038/s41586-019-1773-3

15. Li, S. et al. CellContrast: Reconstructing spatial relationships in single-cell RNA sequencing data via deep contrastive learning. Patterns (N Y) 5, 101022 (2024). 10.1016/j.patter.2024.101022

16. Zhang, Q. et al. Leveraging spatial transcriptomics data to recover cell locations in single-cell RNA-seq with CeLEry. Nat Commun 14, 4050 (2023). 10.1038/s41467-023-39895-3

17. Tan, Y. et al. Transfer learning of multicellular organization via single-cell and spatial transcriptomics. PLoS Comput Biol 21, e1012991 (2025). 10.1371/journal.pcbi.1012991

18. Yu, T. et al. Tissue reassembly with generative AI. bioRxiv (2025). 10.1101/2025.02.13.638045

19. Wu, Z. et al. Graph deep learning for the characterization of tumour microenvironments from spatial protein profiles in tissue specimens. Nat Biomed Eng 6, 1435–1448 (2022). 10.1038/s41551-022-00951-w

20. Singhal, V. et al. BANKSY unifies cell typing and tissue domain segmentation for scalable spatial omics data analysis. Nat Genet 56, 431–441 (2024). 10.1038/s41588-024-01664-3

21. Haviv, D. et al. The covariance environment defines cellular niches for spatial inference. Nat Biotechnol 43, 269–280 (2025). 10.1038/s41587-024-02193-4

22. Varrone, M., Tavernari, D., Santamaria-Martinez, A., Walsh, L. A. & Ciriello, G. CellCharter reveals spatial cell niches associated with tissue remodeling and cell plasticity. Nat Genet 56, 74–84 (2024). 10.1038/s41588-023-01588-4

23. Schroeder, A. et al. Scaling up spatial transcriptomics for large-sized tissues: uncovering cellular-level tissue architecture beyond conventional platforms with iSCALE. bioRxiv (2025). 10.1101/2025.02.25.640190

24. Rood, J. E. et al. The Human Cell Atlas from a cell census to a unified foundation model. Nature 637, 1065–1071 (2025). 10.1038/s41586-024-08338-4

25. Lakkis, J. et al. A joint deep learning model enables simultaneous batch effect correction, denoising, and clustering in single-cell transcriptomics. Genome Res 31, 1753–1766 (2021). 10.1101/gr.271874.120

26. Li, Z., Wang, T., Liu, P. & Huang, Y. SpatialDM for rapid identification of spatially coexpressed ligand-receptor and revealing cell-cell communication patterns. Nat Commun 14, 3995 (2023). 10.1038/s41467-023-39608-w

27. Yao, Z. et al. A transcriptomic and epigenomic cell atlas of the mouse primary motor cortex. Nature 598, 103–110 (2021). 10.1038/s41586-021-03500-8

28. Zhang, M. et al. Spatially resolved cell atlas of the mouse primary motor cortex by MERFISH. Nature 598, 137–143 (2021). 10.1038/s41586-021-03705-x

29. Qian, X. et al. Spatial transcriptomics reveals human cortical layer and area specification. Nature (2025). 10.1038/s41586-025-09010-1

30. Oliveira, M. F. et al. High-definition spatial transcriptomic profiling of immune cell populations in colorectal cancer. Nat Genet 57, 1512–1523 (2025). 10.1038/s41588-025-02193-3

31. Janesick, A. et al. High resolution mapping of the tumor microenvironment using integrated single-cell, spatial and in situ analysis. Nat Commun 14, 8353 (2023). 10.1038/s41467-023-43458-x

32. Zhang, M. et al. Molecularly defined and spatially resolved cell atlas of the whole mouse brain. Nature 624, 343–354 (2023). 10.1038/s41586-023-06808-9

33. Lerma-Martin, C. et al. Cell type mapping reveals tissue niches and interactions in subcortical multiple sclerosis lesions. Nat Neurosci 27, 2354–2365 (2024). 10.1038/s41593-024-01796-z

34. Zhang, D. et al. Inferring super-resolution tissue architecture by integrating spatial transcriptomics with histology. Nat Biotechnol 42, 1372–1377 (2024). 10.1038/s41587-023-02019-9

35. Korsunsky, I. et al. Fast, sensitive and accurate integration of single-cell data with Harmony. Nat Methods 16, 1289–1296 (2019). 10.1038/s41592-019-0619-0

36. Kleshchevnikov, V. et al. Cell2location maps fine-grained cell types in spatial transcriptomics. Nat Biotechnol 40, 661–671 (2022). 10.1038/s41587-021-01139-4

37. Distefano-Gagne, F., Bitarafan, S., Lacroix, S. & Gosselin, D. Roles and regulation of microglia activity in multiple sclerosis: insights from animal models. Nat Rev Neurosci 24, 397–415 (2023). 10.1038/s41583-023-00709-6

38. Davalos, D. et al. Fibrinogen-induced perivascular microglial clustering is required for the development of axonal damage in neuroinflammation. Nat Commun 3, 1227 (2012). 10.1038/ncomms2230

39. Macnair, W. et al. snRNA-seq stratifies multiple sclerosis patients into distinct white matter glial responses. Neuron 113, 396–410 e399 (2025). 10.1016/j.neuron.2024.11.016

40. Belien, J. et al. CHIT1 at diagnosis predicts faster disability progression and reflects early microglial activation in multiple sclerosis. Nat Commun 15, 5013 (2024). 10.1038/s41467-024-49312-y

41. Ostendorf, L. et al. SIGLEC1 (CD169): a marker of active neuroinflammation in the brain but not in the blood of multiple sclerosis patients. Sci Rep 11, 10299 (2021). 10.1038/s41598-021-89786-0

42. Liddelow, S. A. et al. Neurotoxic reactive astrocytes are induced by activated microglia. Nature 541, 481–487 (2017). 10.1038/nature21029

43. Cao, J. et al. The single-cell transcriptional landscape of mammalian organogenesis. Nature 566, 496–502 (2019). 10.1038/s41586-019-0969-x

44. Caligiuri, G. & Tuveson, D. A. Activated fibroblasts in cancer: Perspectives and challenges. Cancer Cell 41, 434–449 (2023). 10.1016/j.ccell.2023.02.015

45. Liu, Y. et al. Conserved spatial subtypes and cellular neighborhoods of cancer-associated fibroblasts revealed by single-cell spatial multi-omics. Cancer Cell 43, 905–924 e906 (2025). 10.1016/j.ccell.2025.03.004

46. Du, Y. et al. The cancer-associated fibroblasts related gene CALD1 is a prognostic biomarker and correlated with immune infiltration in bladder cancer. Cancer Cell Int 21, 283 (2021). 10.1186/s12935-021-01896-x

47. Wu, M. H. et al. Targeting galectin-1 in carcinoma-associated fibroblasts inhibits oral squamous cell carcinoma metastasis by downregulating MCP-1/CCL2 expression. Clin Cancer Res 17, 1306–1316 (2011). 10.1158/1078-0432.CCR-10-1824

48. Makutani, Y. et al. Contribution of MMP14-expressing cancer-associated fibroblasts in the tumor immune microenvironment to progression of colorectal cancer. Front Oncol 12, 956270 (2022). 10.3389/fonc.2022.956270

49. Grout, J. A. et al. Spatial Positioning and Matrix Programs of Cancer-Associated Fibroblasts Promote T-cell Exclusion in Human Lung Tumors. Cancer Discov 12, 2606–2625 (2022). 10.1158/2159-8290.CD-21-1714

50. Xiang, H. et al. Single-Cell Analysis Identifies NOTCH3-Mediated Interactions between Stromal Cells That Promote Microenvironment Remodeling and Invasion in Lung Adenocarcinoma. Cancer Res 84, 1410–1425 (2024). 10.1158/0008-5472.CAN-23-1183

51. Xiong, G. et al. Hsp47 promotes cancer metastasis by enhancing collagen-dependent cancer cell-platelet interaction. Proc Natl Acad Sci U S A 117, 3748–3758 (2020). 10.1073/pnas.1911951117

52. Spada, S., Tocci, A., Di Modugno, F. & Nistico, P. Fibronectin as a multiregulatory molecule crucial in tumor matrisome: from structural and functional features to clinical practice in oncology. J Exp Clin Cancer Res 40, 102 (2021). 10.1186/s13046-021-01908-8

53. Rafi, J. H. et al. High expression of bone morphogenetic protein 1 (BMP1) is associated with a poor survival rate in human gastric cancer, a dataset approaches. Genomics 113, 1141–1154 (2021). 10.1016/j.ygeno.2020.11.012

54. Cretu, A., Sha, X., Tront, J., Hoffman, B. & Liebermann, D. A. Stress sensor Gadd45 genes as therapeutic targets in cancer. Cancer Ther 7, 268–276 (2009).

55. Chen, X. et al. Single-cell resolution spatial analysis of antigen-presenting cancer-associated fibroblast niches. Cancer Cell (2025). 10.1016/j.ccell.2025.09.001

56. Amit, I. et al. The commitment of the human cell atlas to humanity. Nat Commun 15, 10019 (2024). 10.1038/s41467-024-54306-x

57. Park, J. et al. The Spatial Atlas of Human Anatomy (SAHA): A Multimodal Subcellular-Resolution Reference Across Human Organs. bioRxiv (2025). 10.1101/2025.06.16.658716

58. Jin, S. et al. Inference and analysis of cell-cell communication using CellChat. Nat Commun 12, 1088 (2021). 10.1038/s41467-021-21246-9

59. Wolf, F. A., Angerer, P. & Theis, F. J. SCANPY: large-scale single-cell gene expression data analysis. Genome Biol 19, 15 (2018). 10.1186/s13059-017-1382-0

60. Xu, S. et al. Using clusterProfiler to characterize multiomics data. Nat Protoc 19, 3292–3320 (2024). 10.1038/s41596-024-01020-z

