## Supplementary Material for "Reconstructing multi-scale tissue spatial architecture from single-cell RNA-seq with REMAP"

**Supplementary Fig. 1 | Cluster-specific location prediction plot for the mouse brain data.** Scatterplot displaying the ground-truth and predicted locations by REMAP, CeLEry, and iSORT for each cluster.

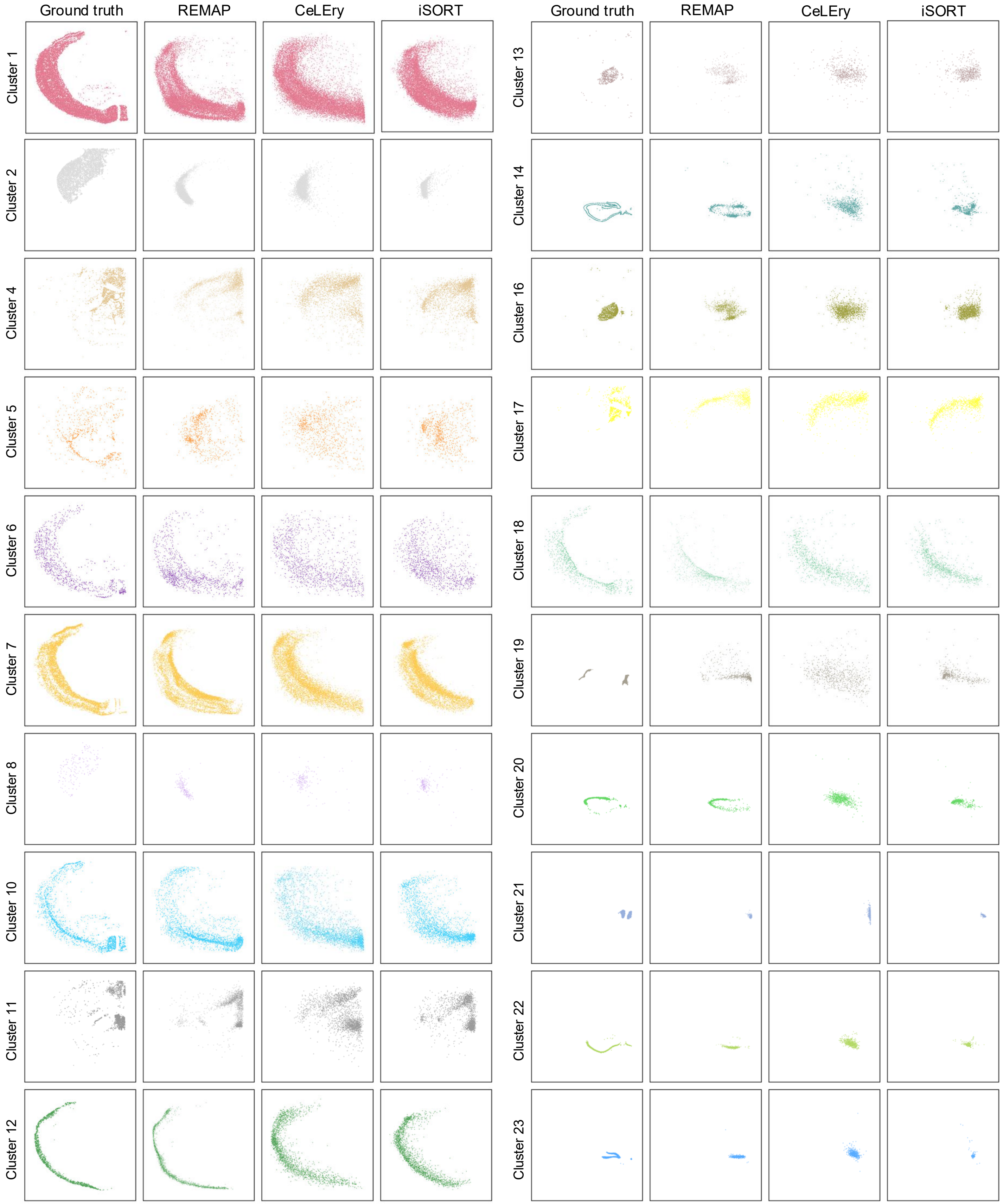

**Supplementary Fig. 2 | Evaluation for the setting with missing cell types in the ST reference for the mouse brain data. a**, Overview of study design. From the original ST reference (left), subsets of cells from both the cortical layer and the hippocampus regions were removed, forming the ST data with missing cell types (middle). We used this as the reference data for model input for all methods and tested on the 10x Visium HD mouse brain data (right). **b**, Ground-truth and predicted cell locations by REMAP and other methods for all regions, cortical layer regions, and hippocampus regions. **c**, Barplot of Pearson correlation between true and predicted pairwise distances.

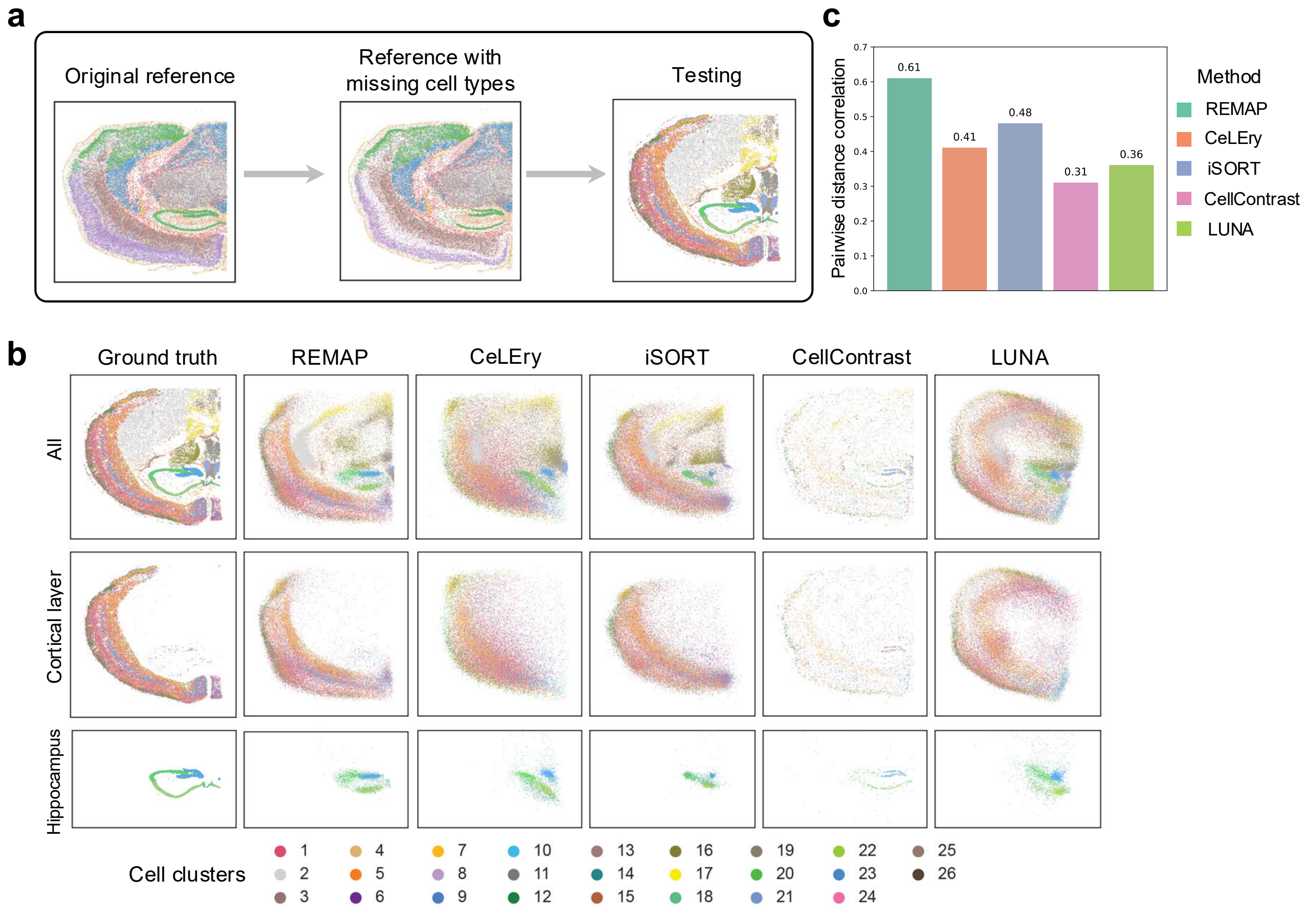

**Supplementary Fig. 3 | CN clustering and spatial networks in the mouse brain data. a**, Mean error between true and predicted pairwise Euclidean distances among CN cluster centroids as a function of CN cluster number. **b**, NMI between true and predicted CN clusters as a function of CN cluster number. **c**, Here, we performed CN clustering based on predicted locations for each method separately, and constructed spatial networks based on predicted CN clusters. In the top panel, cells are plotted under predicted locations and colored by predicted CN clusters; in the bottom panel, nodes represent the centroids of CN clusters and are connected based on their spatial proximity. Edge length corresponds to the Euclidean distance between centroids, and edge width was scaled inversely to distance, such that thicker edges indicate closer proximity. Node size is proportional to the standard deviation of within-cluster pairwise distances among cells. **d**, Sankey plots showing the correspondence between ground-truth and predicted CN clusters for each method, with NMI between two sets of cluster memberships indicated.

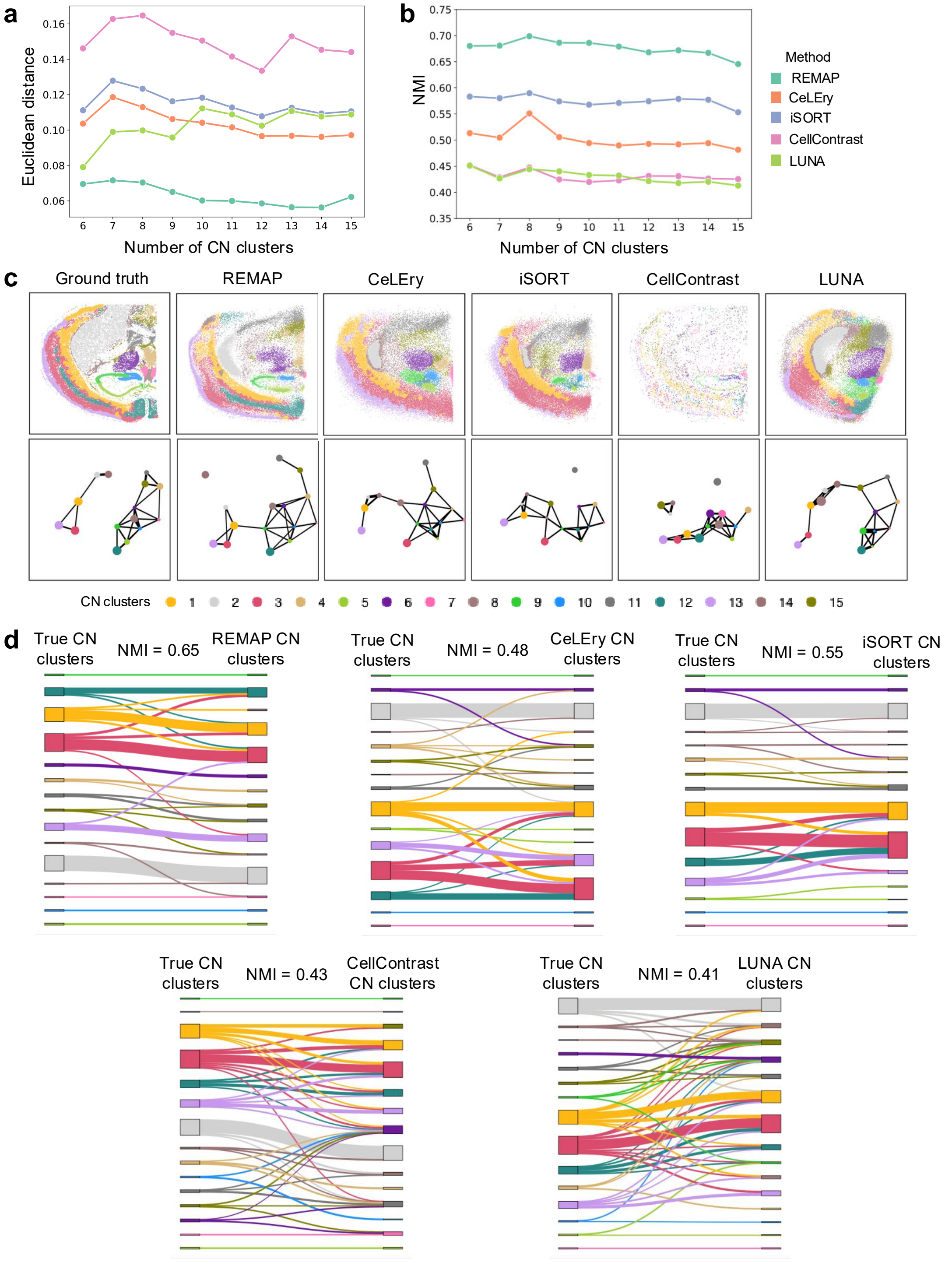

**Supplementary Fig. 4 | Cell-cell communication results under ground truth and REMAP locations in the mouse brain data.** For each ligand-receptor pair, the scatterplot shows the interacting strengths (i.e., 1 – *p*-value) for all cells inferred by SpatialDM, and the chord diagram represents the interaction patterns at the cell-cluster level, with node colors indicating cell types and edge colors denoting sender cell types.

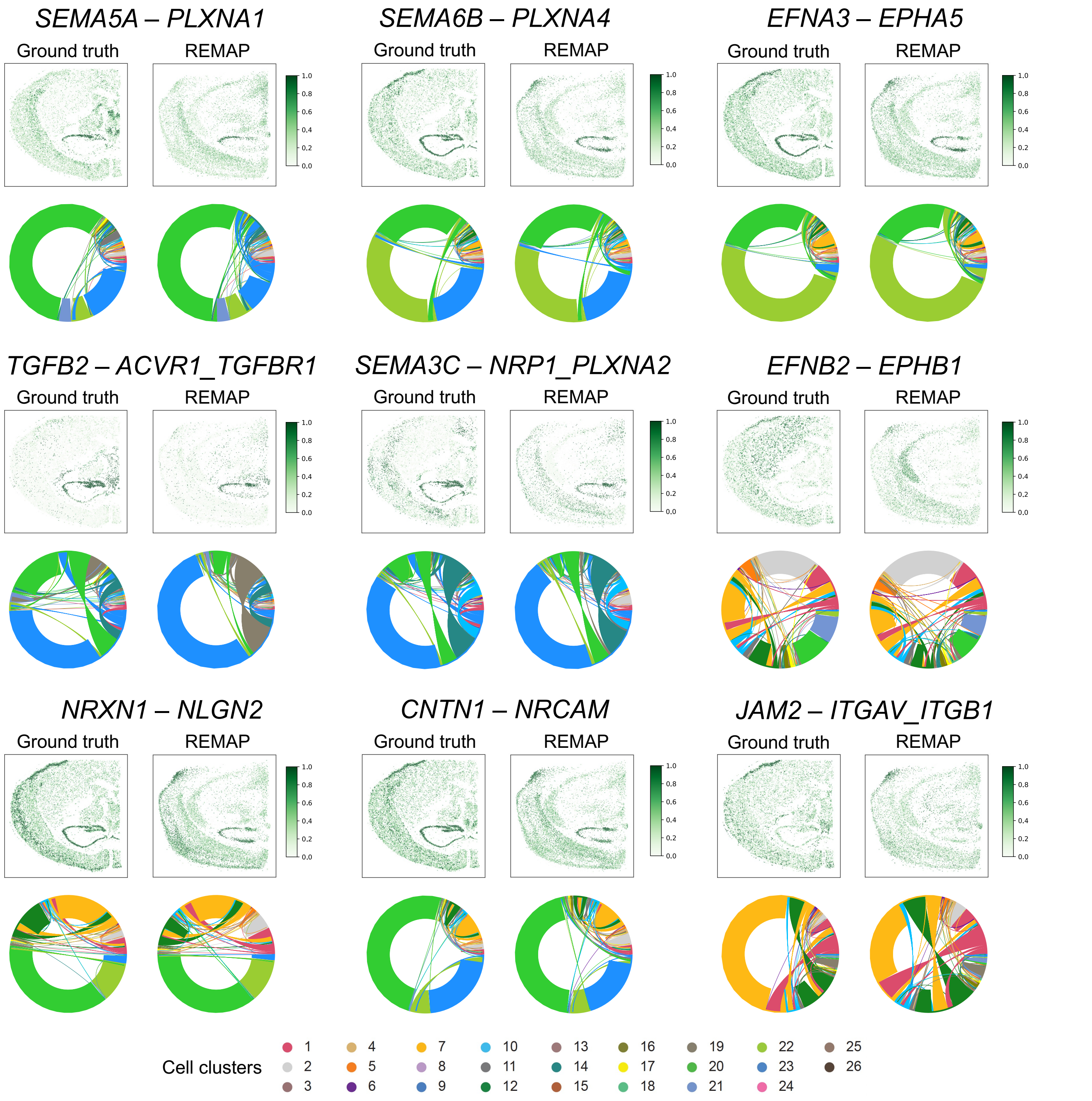

**Supplementary Fig. 5 | Application to mouse primary motor cortex data**. **a,** Two benchmarking scenarios for the mouse primary motor cortex data evaluation. Scenario 1: a single MERFISH slice was used as the reference, with a scRNA-seq data as the test dataset; Scenario 2: different numbers of MERFISH slices were used as the reference, with cortical layer regions from 10x Visium HD mouse brain data as the test. **b,** Predicted cell locations by REMAP and other methods in Scenario 1. **c,** Cortical depth distributions of different layers based on reference data and recovered locations in Scenario 1. **d,** Ground-truth and predicted cell locations by REMAP and LUNA in Scenario 2, using eight MERFISH slices as the reference. **e,** Pearson correlation between true and predicted pairwise distances across varying numbers of reference slices.

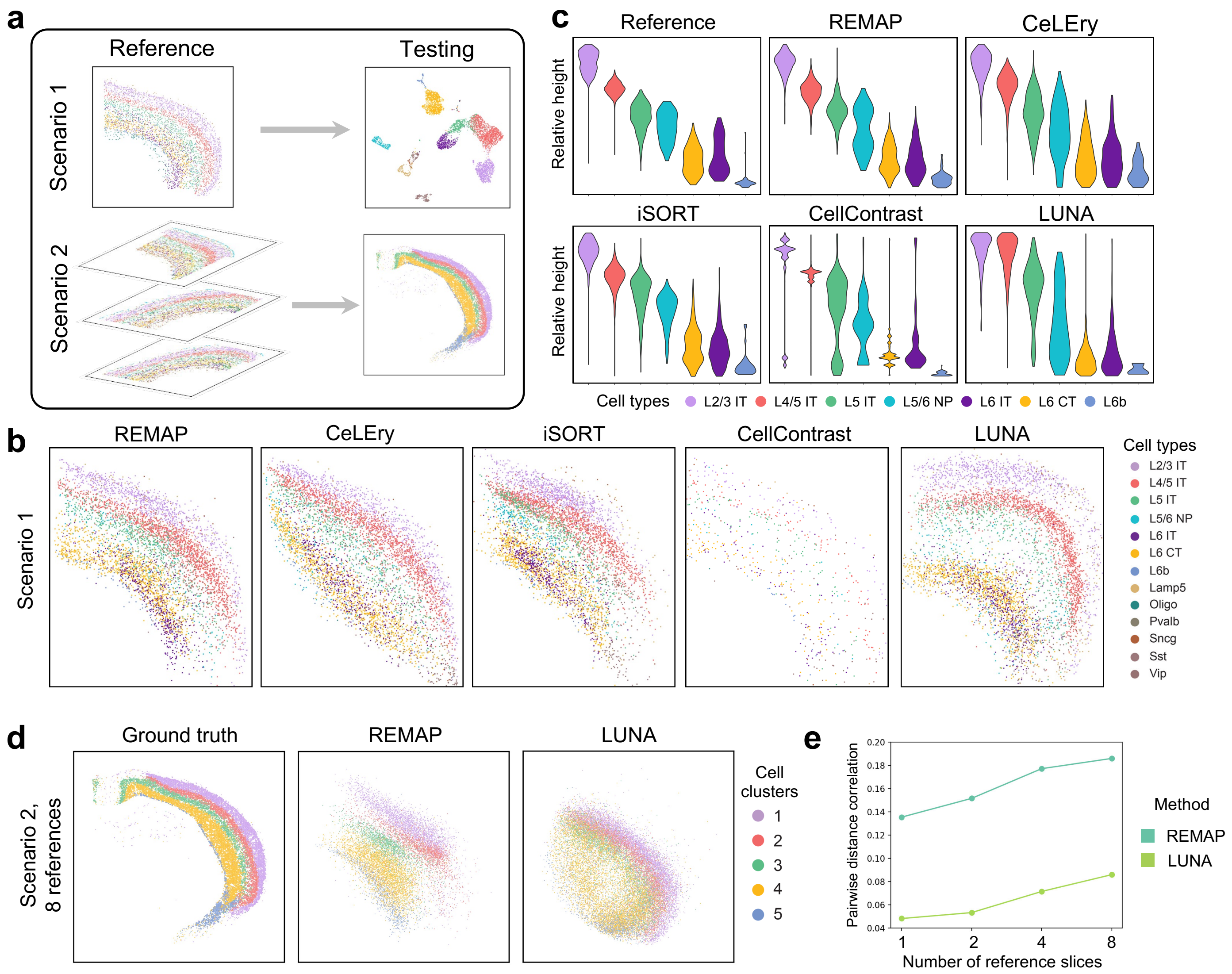

**Supplementary Fig. 6 | Recovered gene expression spatial heatmap of layer-specific marker genes in the mouse primary cortex data based on REMAP-recovered locations in scenario 1, with color indicating relative gene expression.**

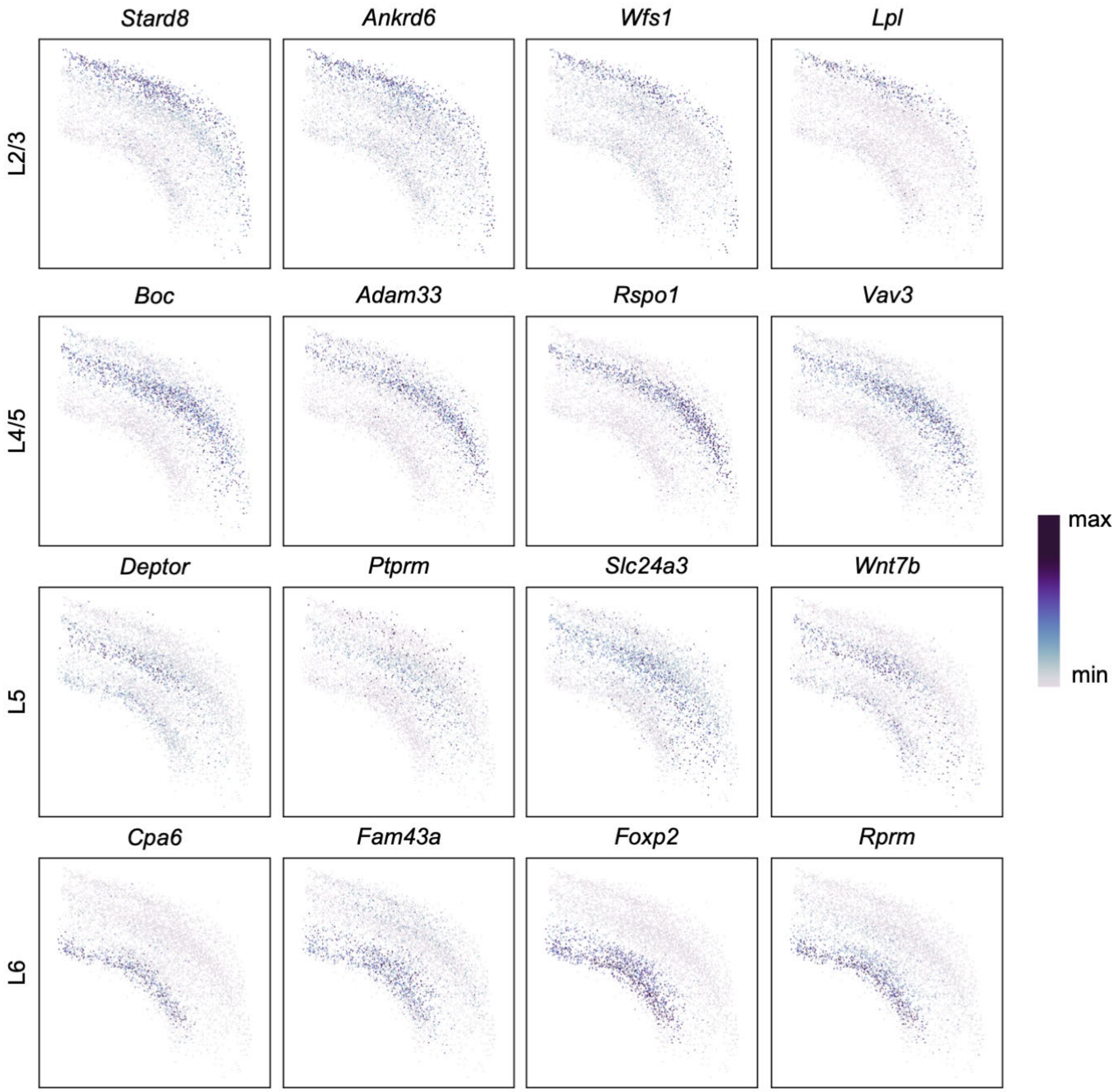

**Supplementary Fig. 7 | Evaluation for the setting with missing cell types in the ST reference for the mouse primary cortex data in scenario 1.** From the original ST reference, we performed leave-one-out experiments. One cell type was removed at a time, including L4/5, L5, L6 IT and L6 CT, and used as the reference input for all methods. Reference and predicted cell locations for each scenario and methods were shown.

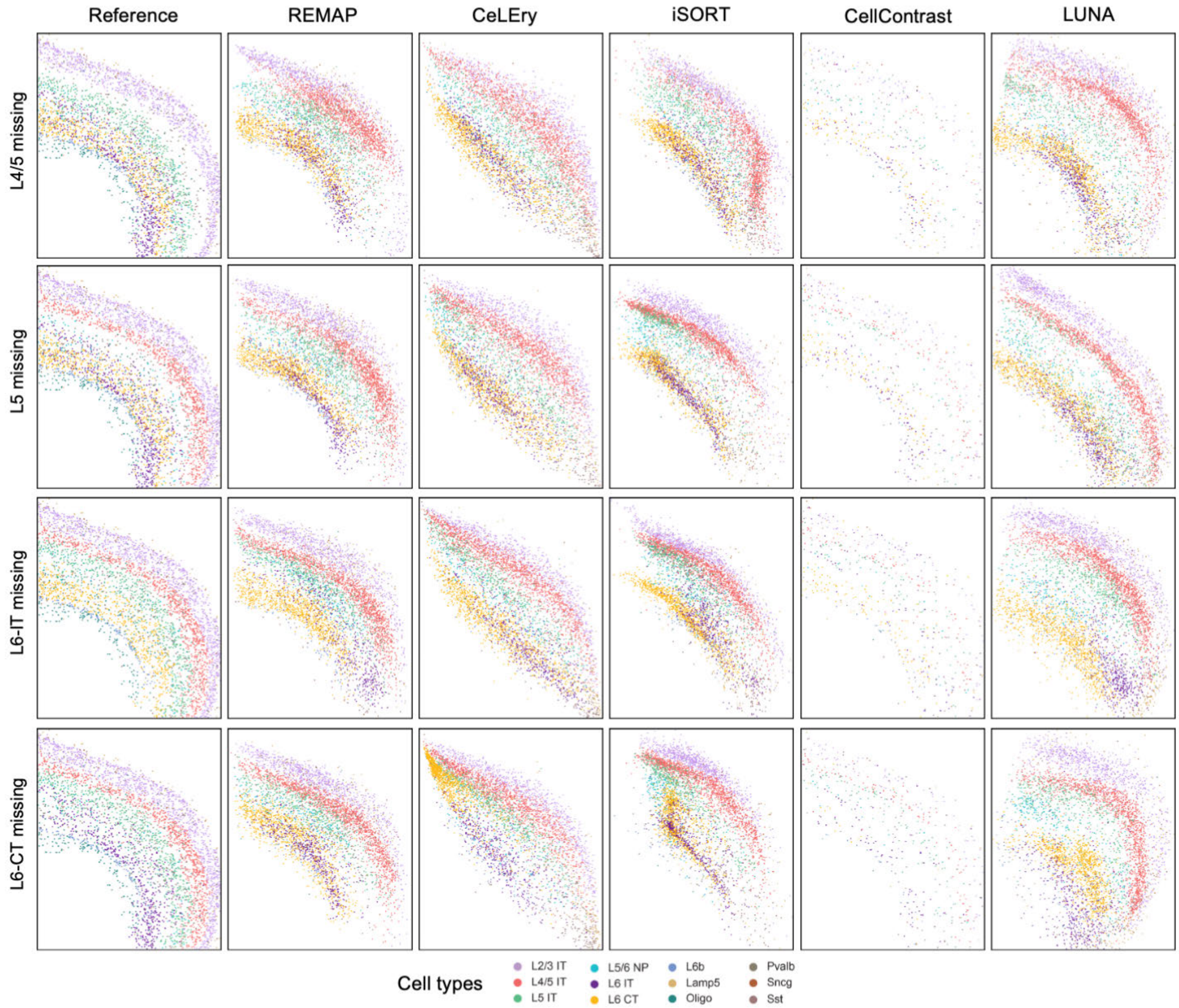

**Supplementary Fig. 8 | Location prediction plots in the mouse primary cortex data in scenario 2.** We progressively expanded the reference by incorporating 1, 2, 4, or 8 slices of varying orientations to predict cortical layer cell locations in the mouse brain Visium HD dataset. Ground-truth and predicted cell locations by REMAP and LUNA using different numbers of reference slices.

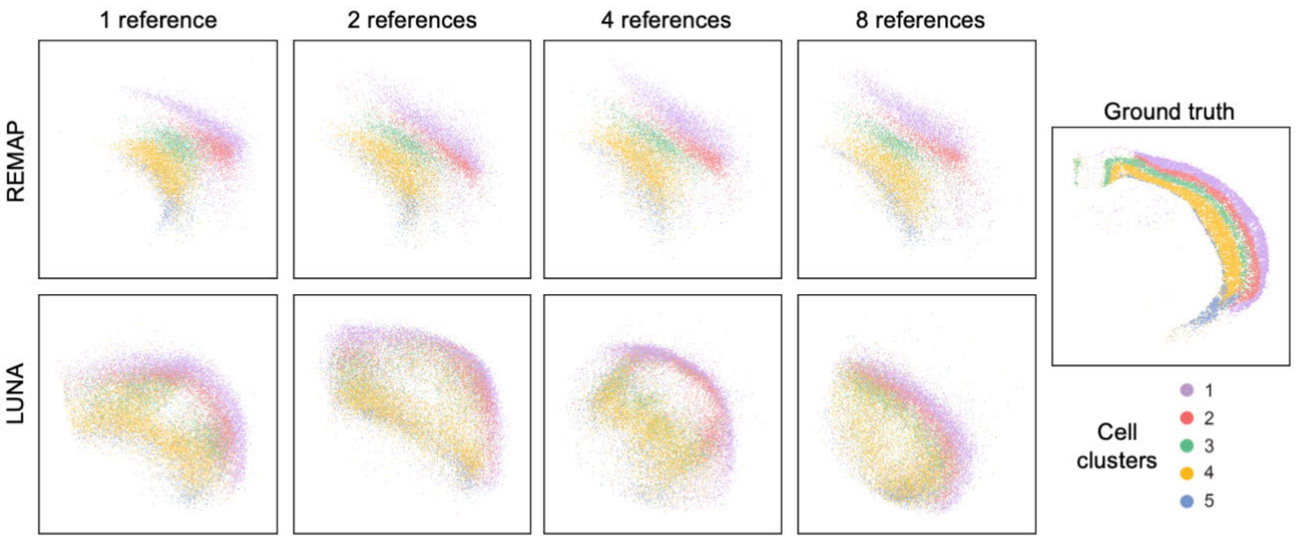

**Supplementary Fig. 9 | Location prediction plot for the human fetal cortex data in scenario 1. a**, Ground-truth and predicted cell locations for all clusters by REMAP and other methods. **b**, Ground-truth and predicted cell locations for V1/V2-specific clusters by REMAP and other methods. **c**, Cluster-specific location prediction plots for V1/V2 specific clusters by REMAP and other methods. **d**, Pearson correlation between true and predicted pairwise distances.

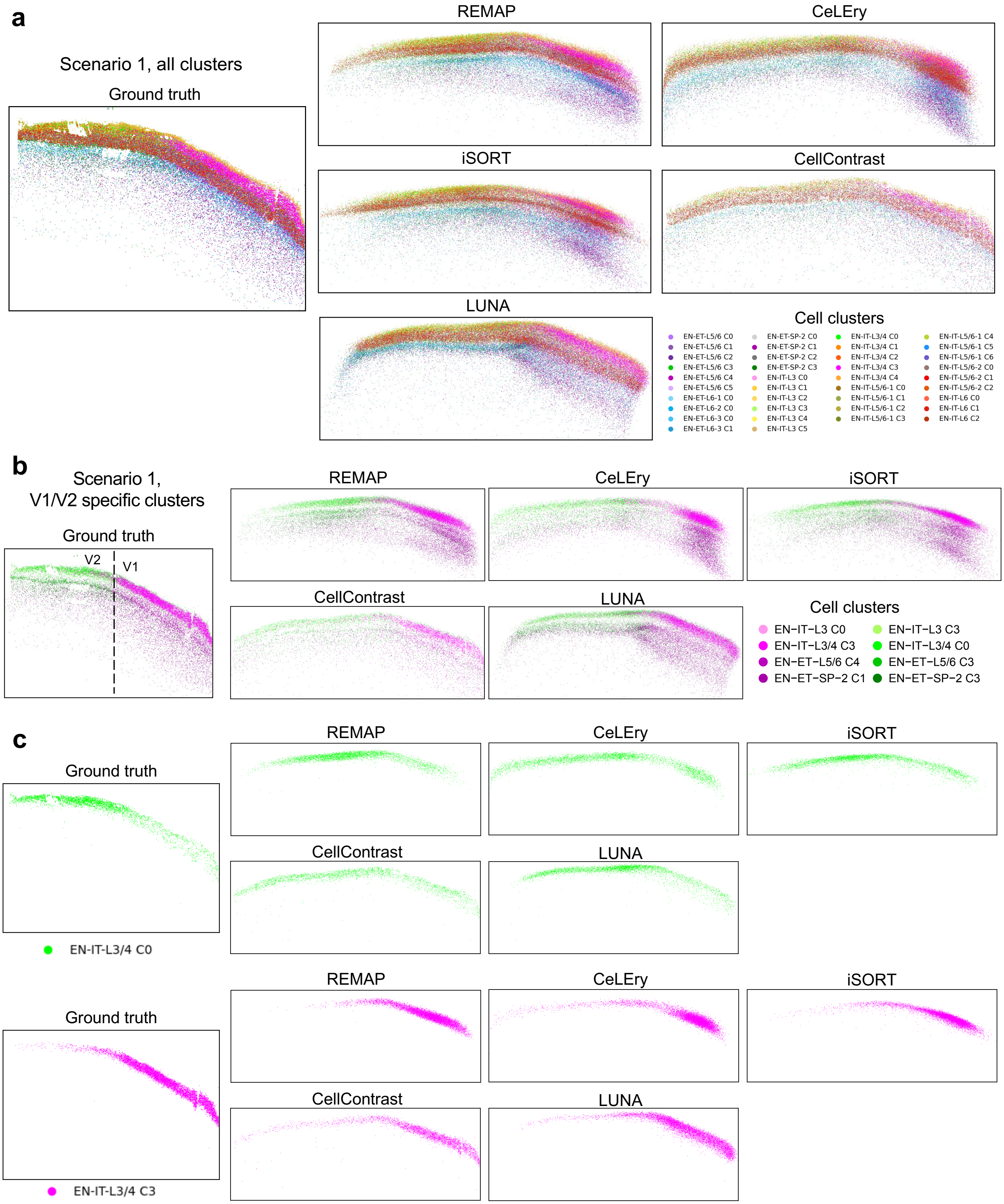

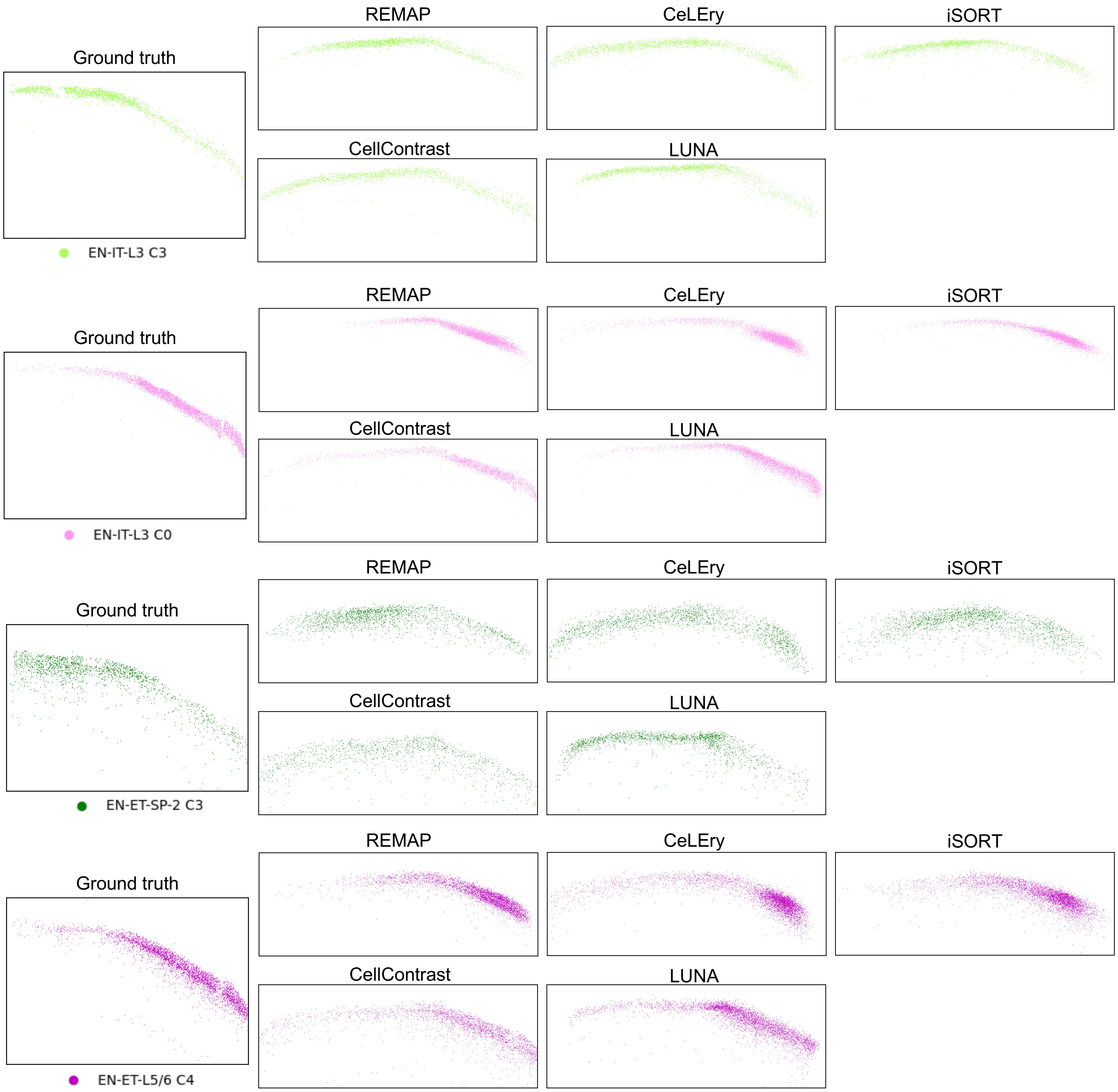
Ground truth
REMAP
CeLEry
iSORT
CellContrast
LUNA
EN-IT-L3 C3
Ground truth
REMAP
CeLEry
iSORT
CellContrast
LUNA
EN-IT-L3 C0
Ground truth
REMAP
CeLEry
iSORT
CellContrast
LUNA
EN-ET-SP-2 C3
Ground truth
REMAP
CeLEry
iSORT
CellContrast
LUNA
EN-ET-L5/6 C4

**d**

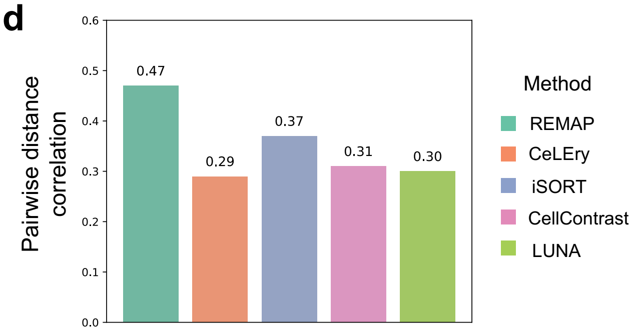
Pairwise distance correlation
0.0
0.1
0.2
0.3
0.4
0.5
0.6
0.47
0.29
0.37
0.31
0.30
Method
REMAP
CeLEry
iSORT
CellContrast
LUNA

**Supplementary Fig. 10 | CN clustering and spatial networks based on true CN clusters for the human fetal cortex data in scenario 1.** Here, we performed CN clustering based on ground-truth locations and constructed spatial networks based on ground-truth CN clusters. In the scatterplots, cells are plotted under predicted locations and colored by ground-truth CN clusters; in the spatial network, nodes represent the centroids of CN clusters and are connected based on their spatial proximity. Edge length corresponds to the Euclidean distance between centroids, and edge width was scaled inversely to distance, such that thicker edges indicate closer proximity. Node size is proportional to the standard deviation of within-cluster pairwise distances among cells.

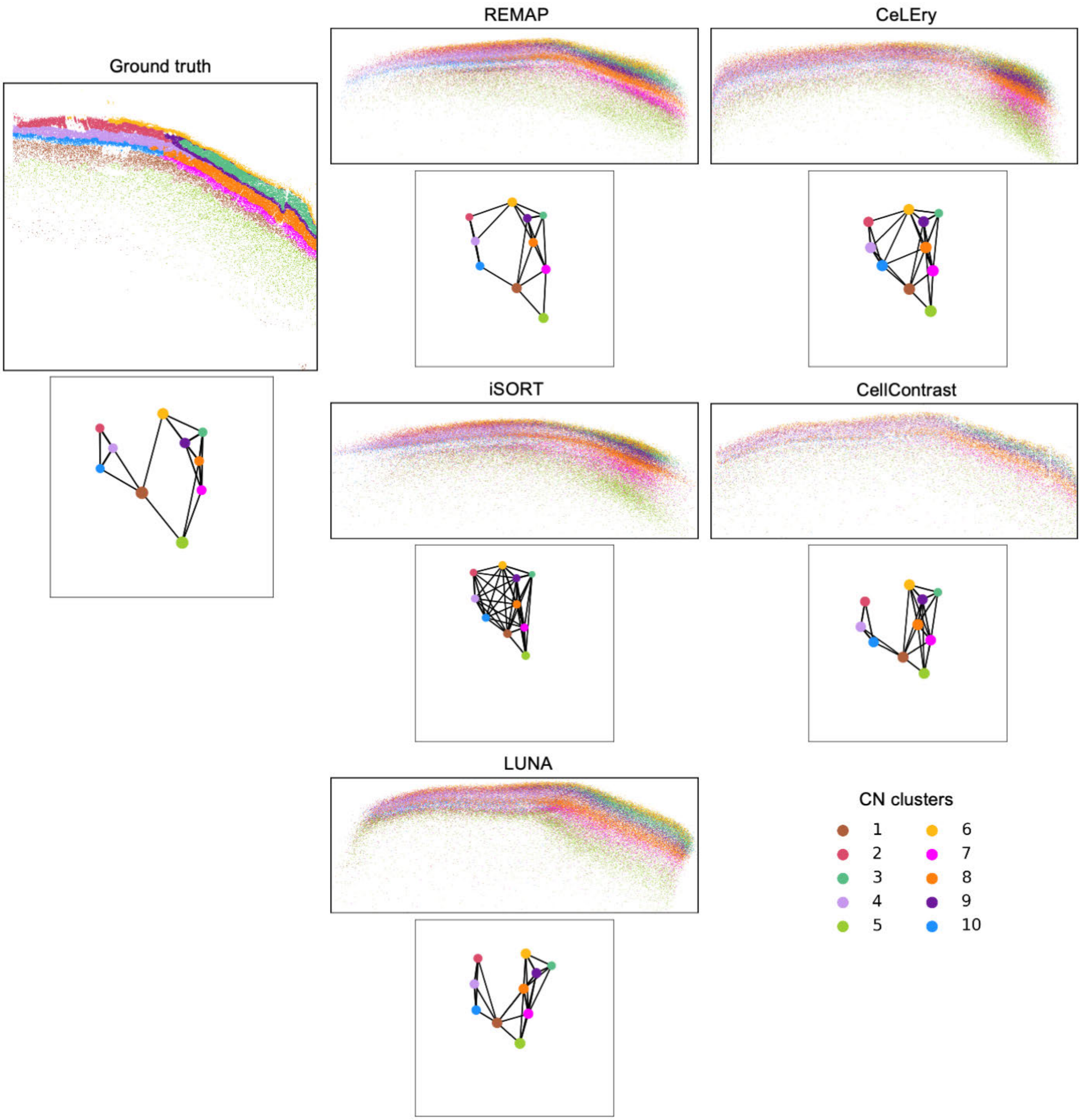

**Supplementary Fig. 11 | CN clustering evaluation metrics for the human fetal cortex data in scenario 1.** **a**, Mean error between true and predicted pairwise Euclidean distances among CN cluster centroids as a function of CN cluster number. **b**, NMI between true and predicted CN clusters as a function of CN cluster number.

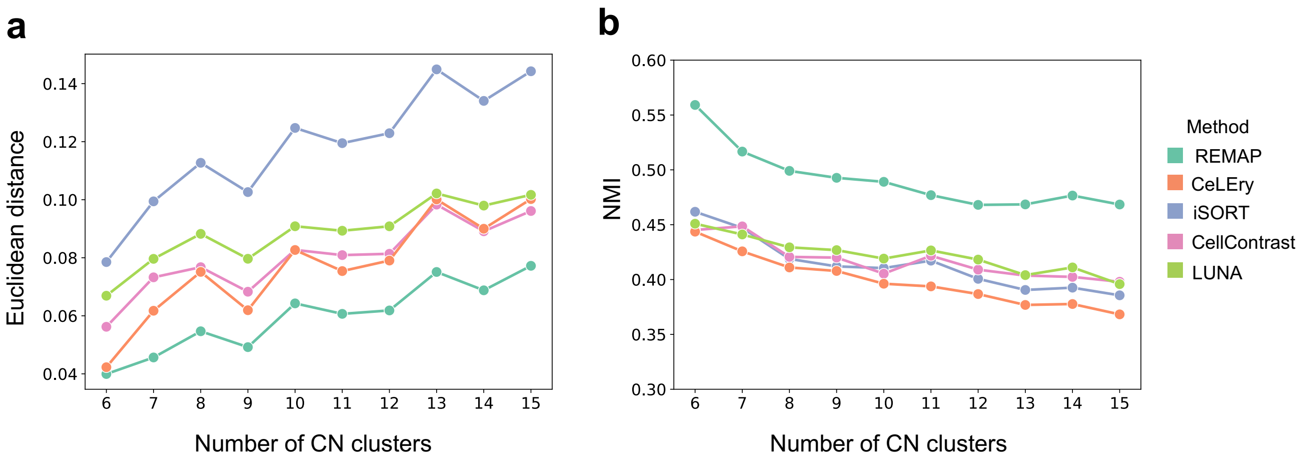

**Supplementary Fig. 12 | Cortical depth quantification for the human fetal cortex data in scenario 2. a**, Three annotated regions for cortical depth quantification based on ST reference**.** “Vertical 1” and “Vertical 2” were combined for vertical depth quantification on both sides of the cortex, while “Horizontal” was used for horizontal depth quantification. For each method, cells with predicted locations falling within the annotated region were identified, and their cortical depth was calculated. **b**, Violin plots showing vertical cortical depth distributions of all cell types and horizontal cortical depth distributions of V1/V2 specific cell types based on recovered locations by CeLEry, iSORT, CellContrast, and LUNA.

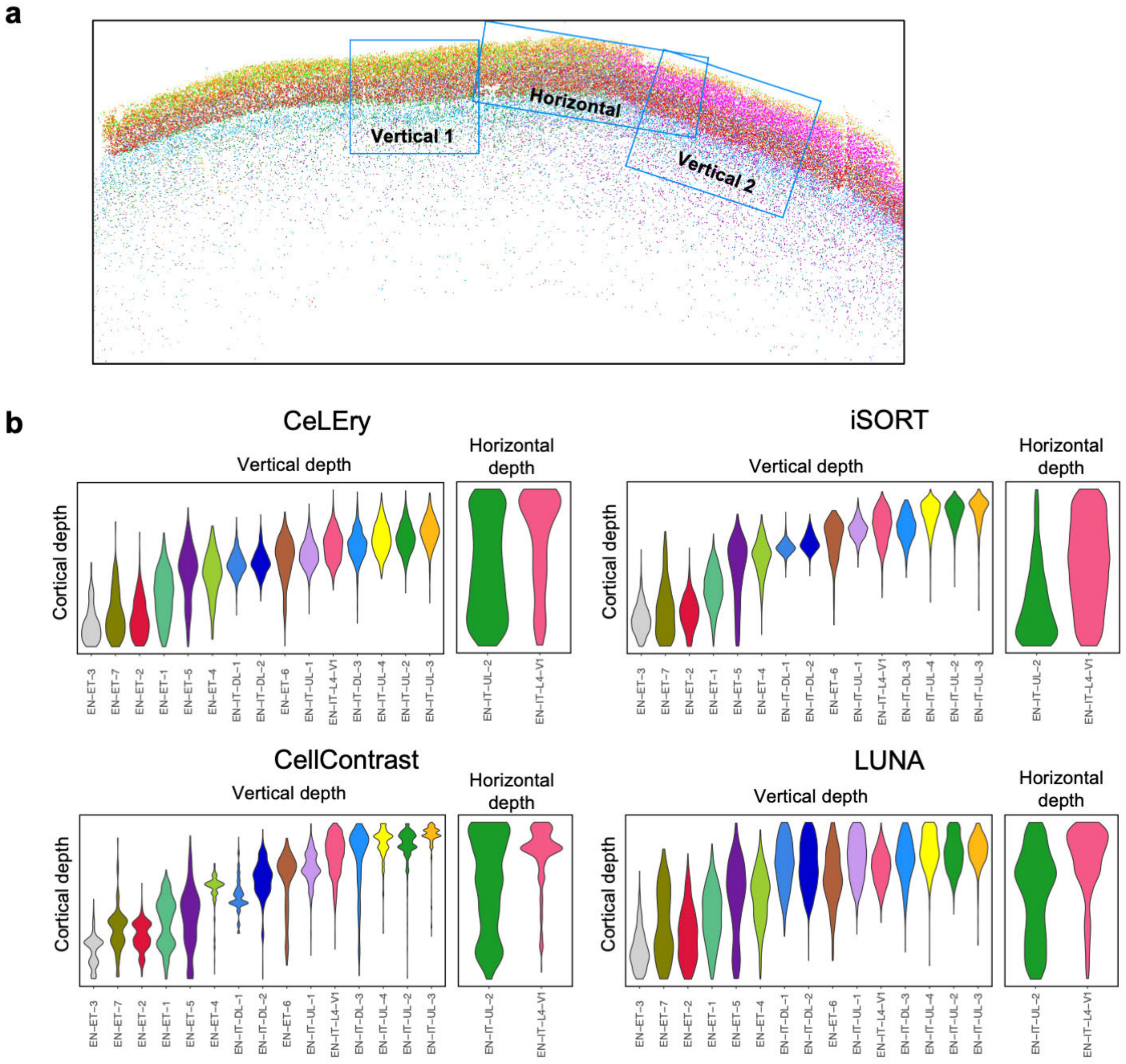

**Supplementary Fig. 13 | CN clustering and spatial networks for the human fetal cortex data in scenario 2.** Here, we performed separate CN clustering based on predicted locations for each method and constructed spatial networks based on predicted CN clusters. **a**, In the scatterplots, cells are plotted under predicted locations and colored by predicted CN clusters for each method; in the spatial network, nodes represent the centroids of CN clusters and are connected based on their spatial proximity. **b**, Top: lollipop plots displaying the mean expression of V1 and V2 marker genes identified in the original study in each CN cluster by REMAP. Bottom: bar plots displaying the fold change of each marker gene in the most enriched CN cluster versus other CN clusters.

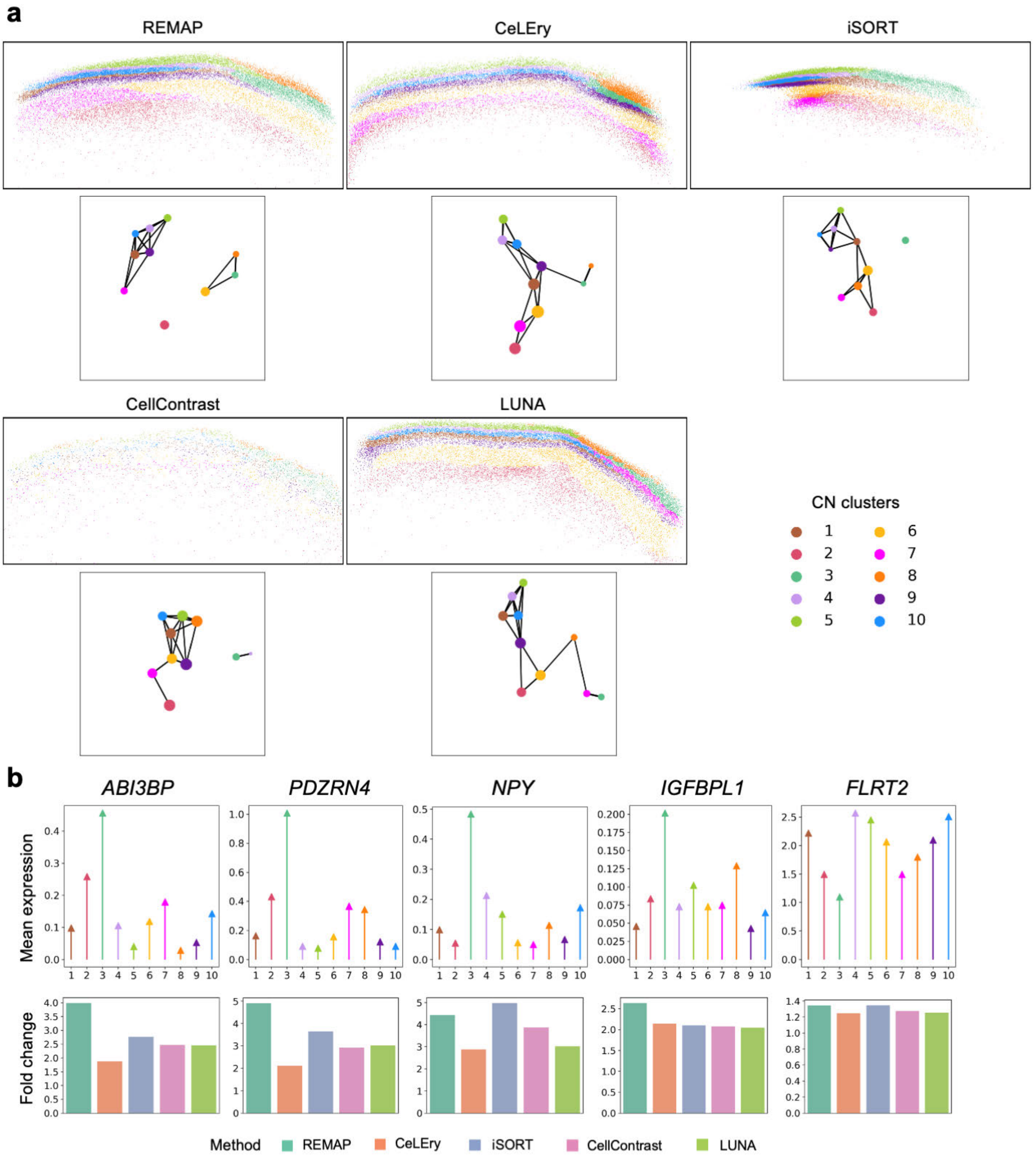

**Supplementary Fig. 14 | Recovered gene expression spatial heatmap of V1- and V2-enriched marker genes in the human fetal cortex data based on REMAP-recovered locations in scenario 2, with color indicating relative gene expression.**

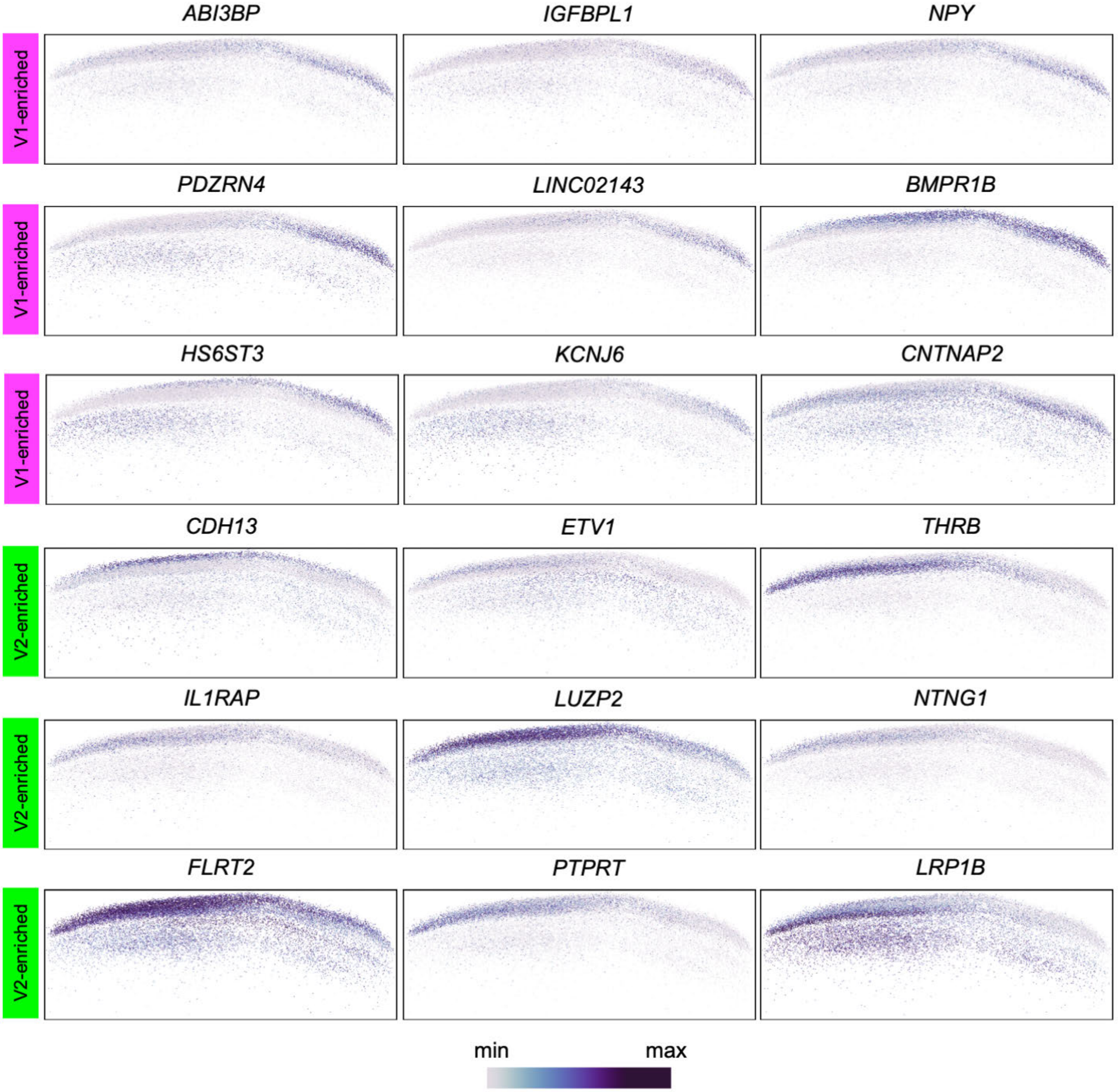

**Supplementary Fig. 15 | Cluster-specific location prediction plot for the human CRC data.** Scatterplot displaying the ground-truth and predicted locations by REMAP, iSORT, and LUNA for each cluster.

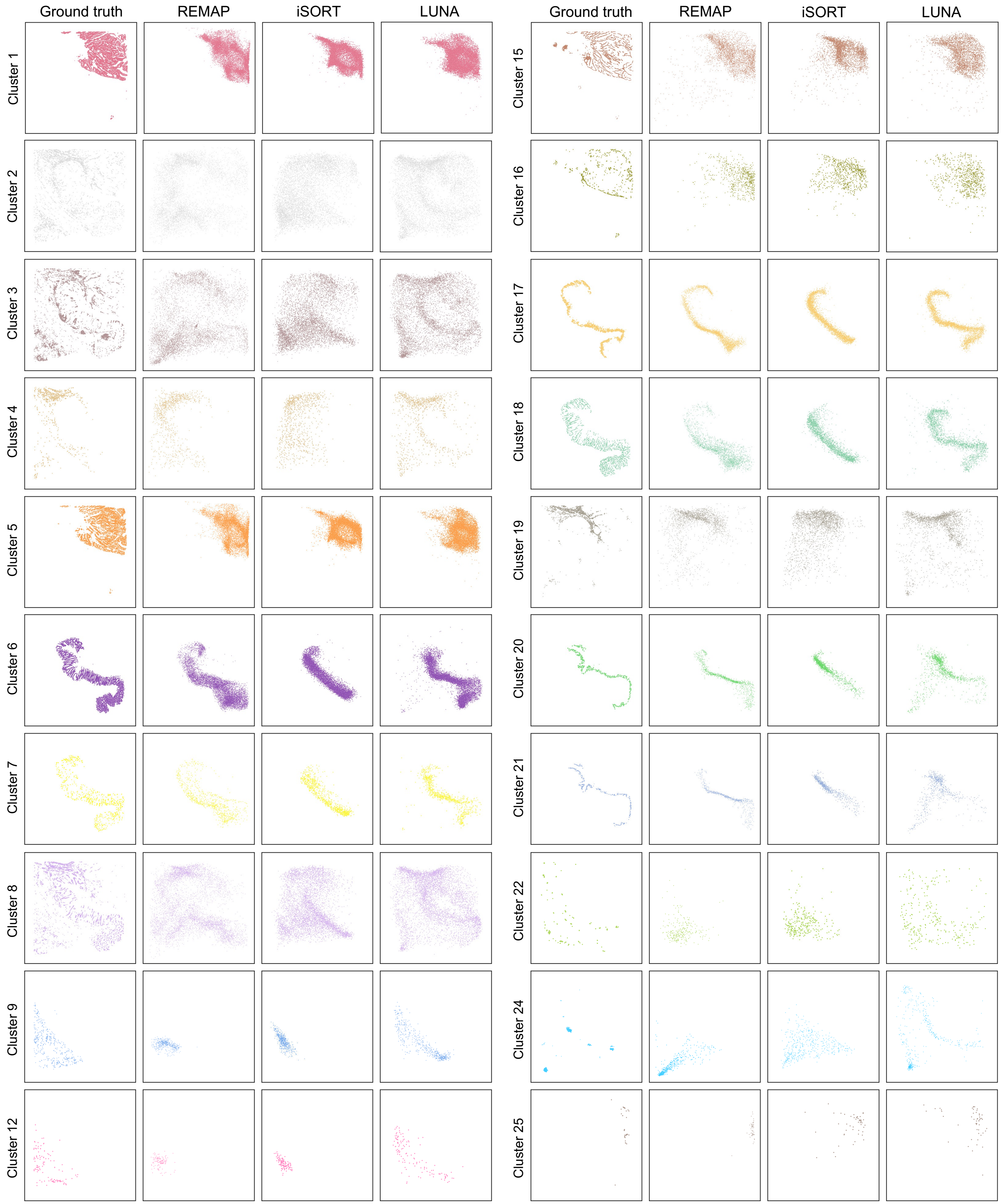

**Supplementary Fig. 16 | Cortical depth quantification for the human CRC data. a**, Annotated region within the goblet cell area for cortical depth quantification based on ST reference**.** For each method, cells with predicted locations falling within the annotated region were identified, and their cortical depth was calculated. **b**, Violin plots showing vertical cortical depth distributions of goblet cell-related clusters based on ground-truth and recovered locations by all methods.

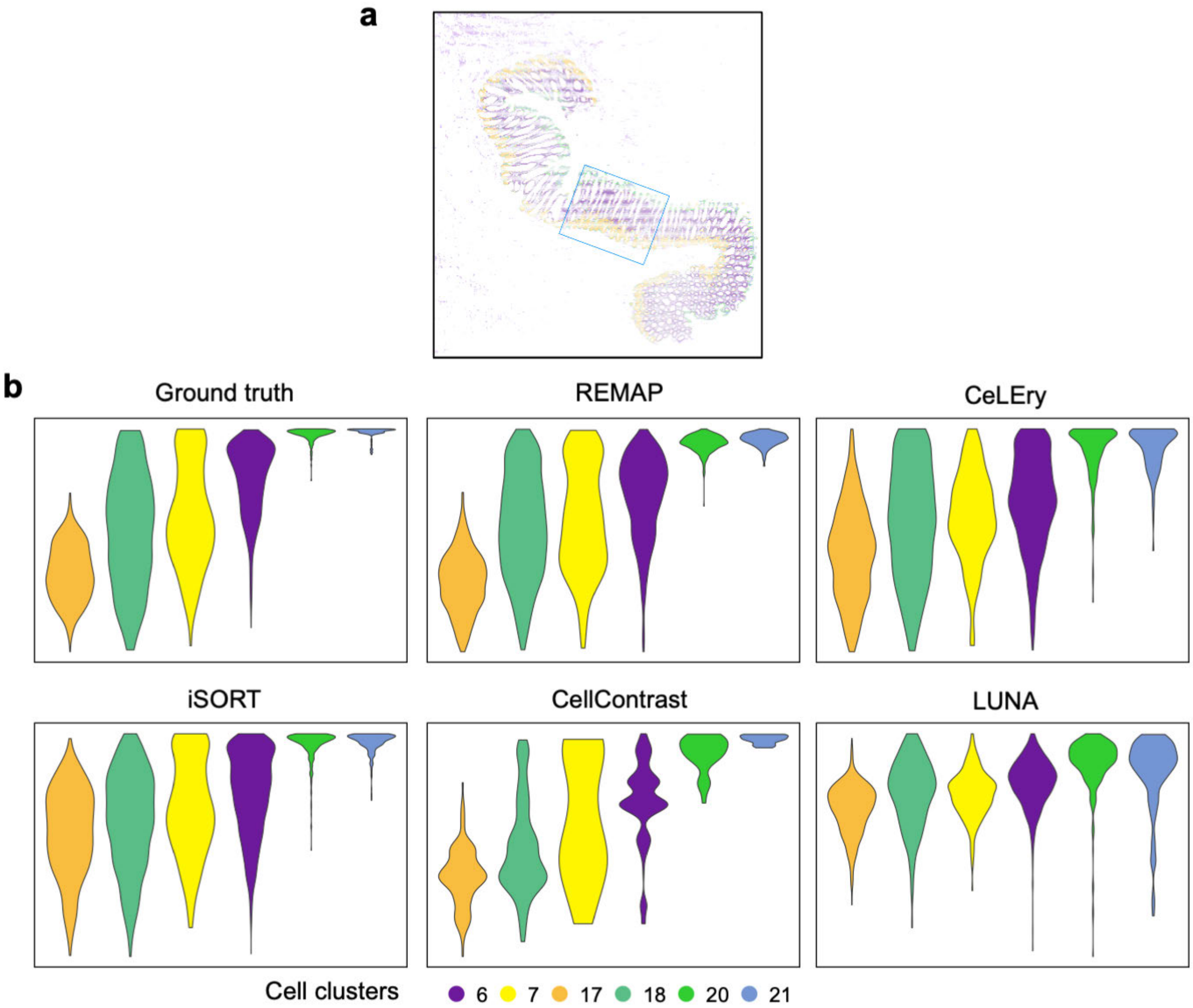

**Supplementary Fig. 17 | CN clustering and spatial networks based on predicted CN clusters for the human CRC data.** Here, we performed separate CN clustering based on predicted locations for each method and constructed spatial networks based on predicted CN clusters. **a**, In the top panel, cells are plotted under predicted locations and colored by predicted CN clusters; in the bottom panel, nodes represent the centroids of CN clusters and are connected based on their spatial proximity. Edge length corresponds to the Euclidean distance between centroids, and edge width was scaled inversely to distance, such that thicker edges indicate closer proximity. Node size is proportional to the standard deviation of within-cluster pairwise distances among cells. **b**, Sankey plots showing the correspondence between ground-truth and predicted CN clusters for each method, with NMI between two sets of cluster memberships indicated.

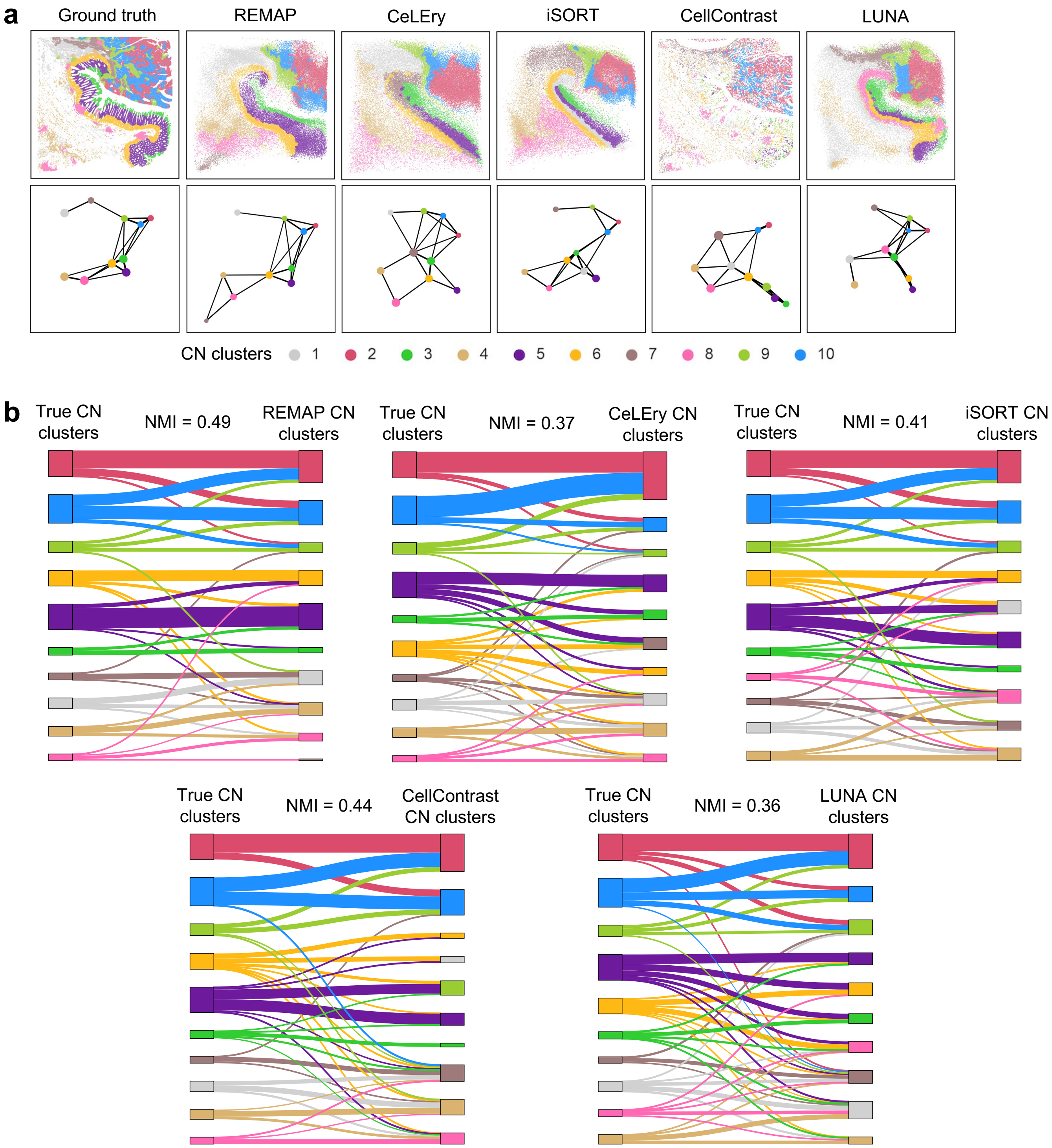

**Supplementary Fig. 18 | Additional examples of ligand-receptor pairs of REMAP-based cell-cell communication results in the human CRC data.** For each ligand-receptor pair, the scatterplot shows the interacting strengths (1 – *p*-value) for all cells, and the chord diagram represents the interaction patterns at the cell-cluster level, with node colors indicating cell types and edge colors denoting sender cell types.

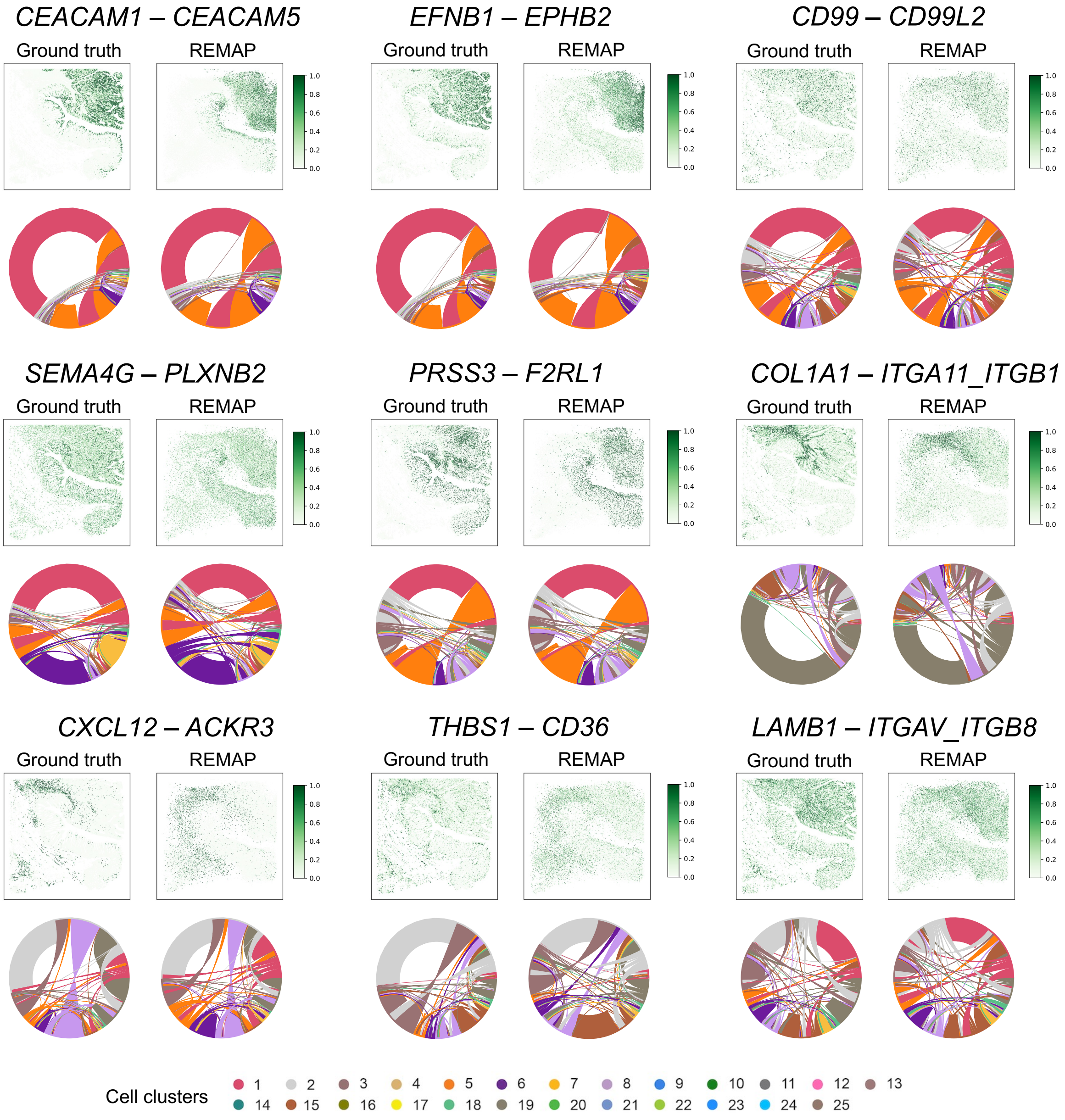

**Supplementary Fig. 19 | Application to human breast cancer data. a**, Study design for the 10x Xenium human breast cancer data benchmarking evaluation. One slice served as the reference, and an adjacent slice served as the test data. Two slices were roughly aligned based on their overlapping regions along the Y axis. **b**, Ground-truth and predicted cell locations by REMAP and other methods for all cell types and Myoepithelial ACTA2+. The region of interest was highlighted in a red circle. **c**, Pearson correlation between true and predicted pairwise distances. **d**, CN clusters and spatial networks constructed by REMAP and other methods. In the top panel, cells were plotted under predicted locations and colored by predicted CN clusters; in the bottom panel, nodes represent the centroids of CN clusters with edges drawn based on spatial proximity. Edge length corresponds to the Euclidean distance between centroids, and edge width was scaled inversely to distance, such that thicker edges indicate closer proximity. Node size is proportional to the standard deviation of within-cluster pairwise distances among cells. **e**, Mean error between true and predicted pairwise Euclidean distances among CN cluster centroids, as a function of CN cluster number. **f**, NMI between true and predicted CN clusters as a function of CN cluster number. **g**, Cell-cell communication results for REMAP. For each ligand-receptor pair, the scatterplot shows the interaction strengths (i.e., 1 – *p*-value) for all cells inferred by SpatialDM, and the chord diagram represents the interaction patterns at the cell-cluster level, with node colors indicating cell types and edge colors denoting sender cell types.

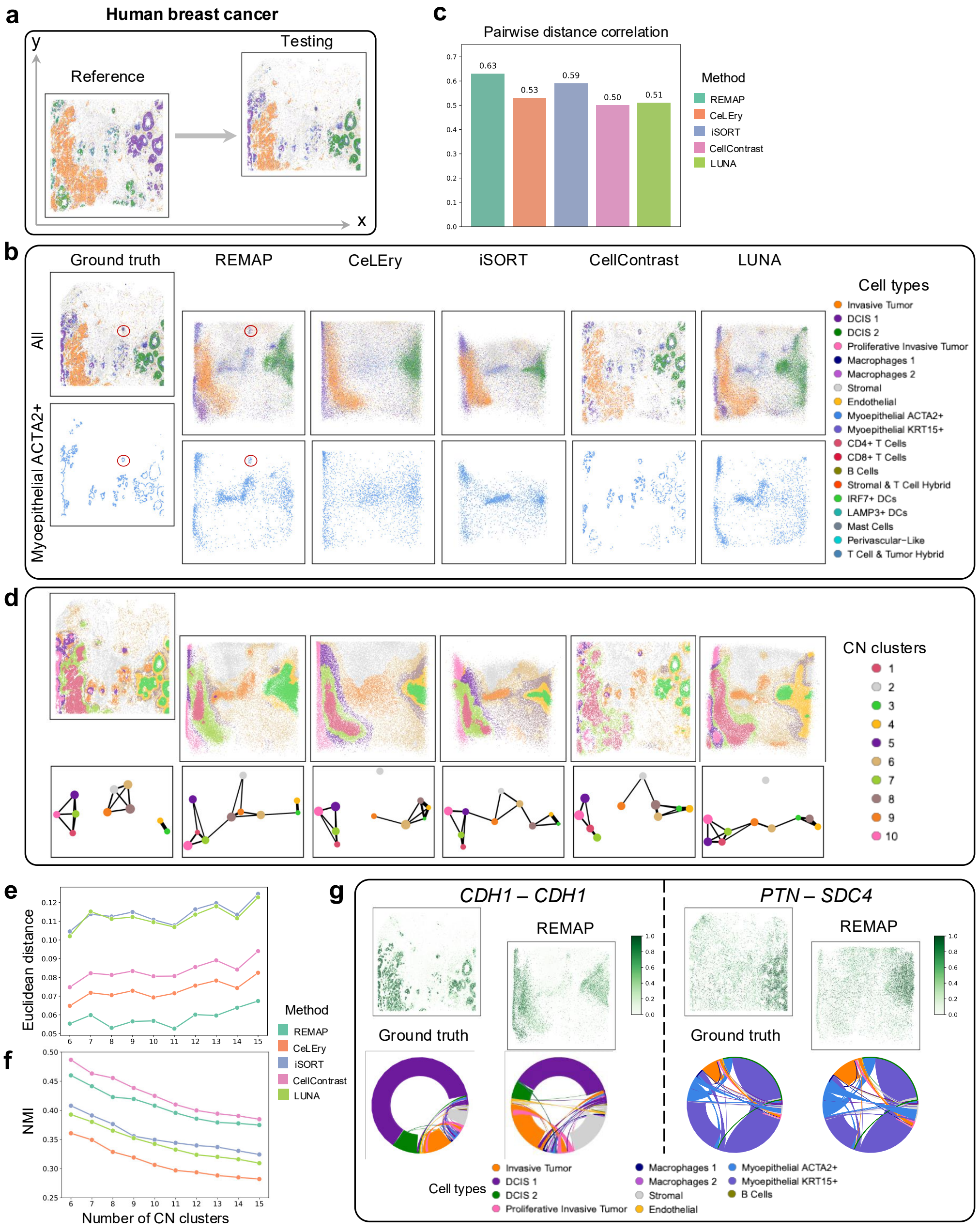

**Supplementary Fig. 20 | Cell-type-specific location prediction plot for the human breast cancer data.** Scatterplot displaying the ground-truth and predicted locations by REMAP, iSORT, and LUNA for each cell type.

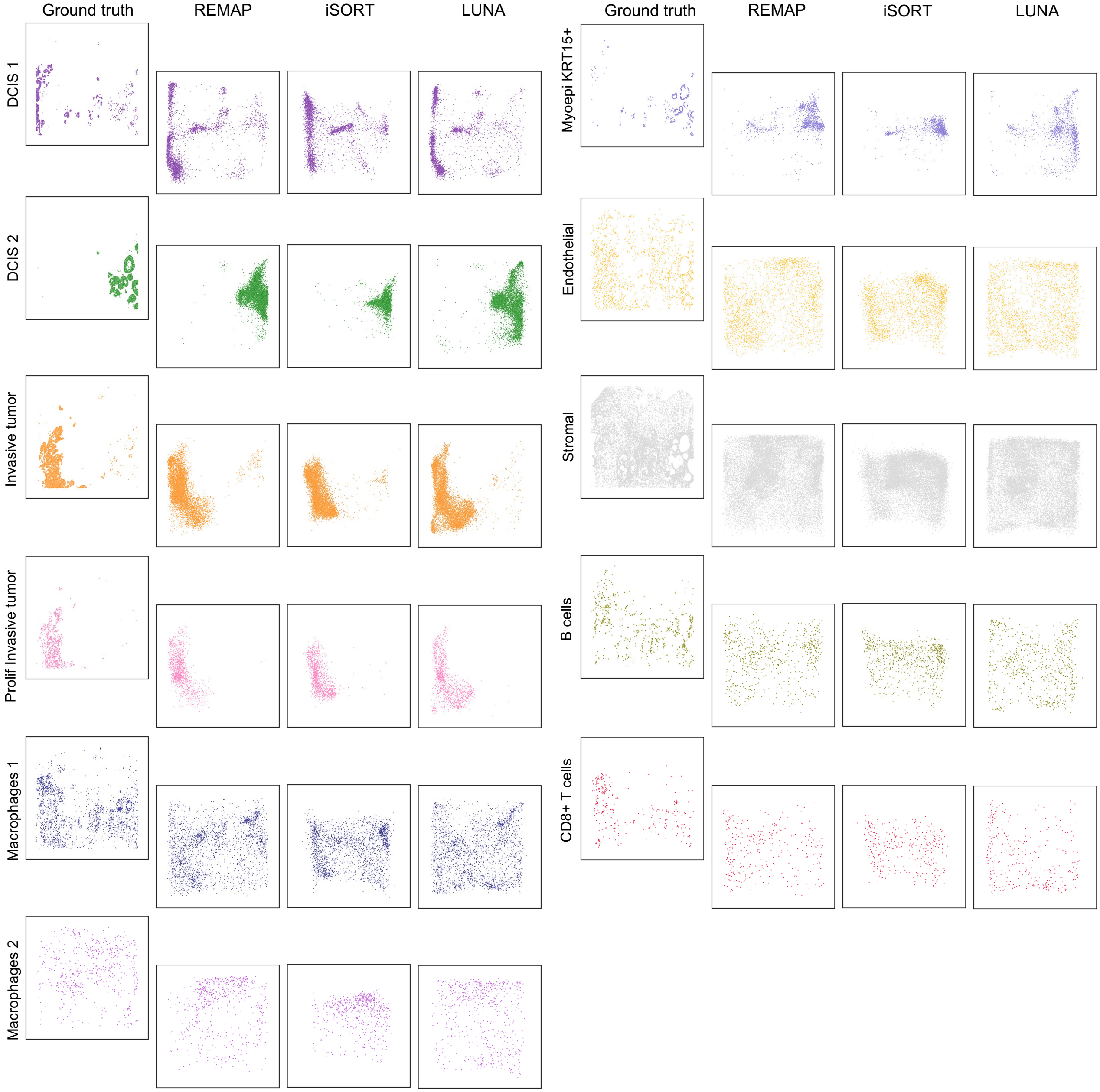

**Supplementary Fig. 21 | Additional examples of ligand-receptor pairs of REMAP-based cell-cell communication results in the human breast cancer data.** For each ligand-receptor pair, the scatterplot shows the interacting strengths (1 – *p*-value) for all cells, and the chord diagram represents the interaction patterns at the cell-type level, with node colors indicating cell types and edge colors denoting sender cell types.

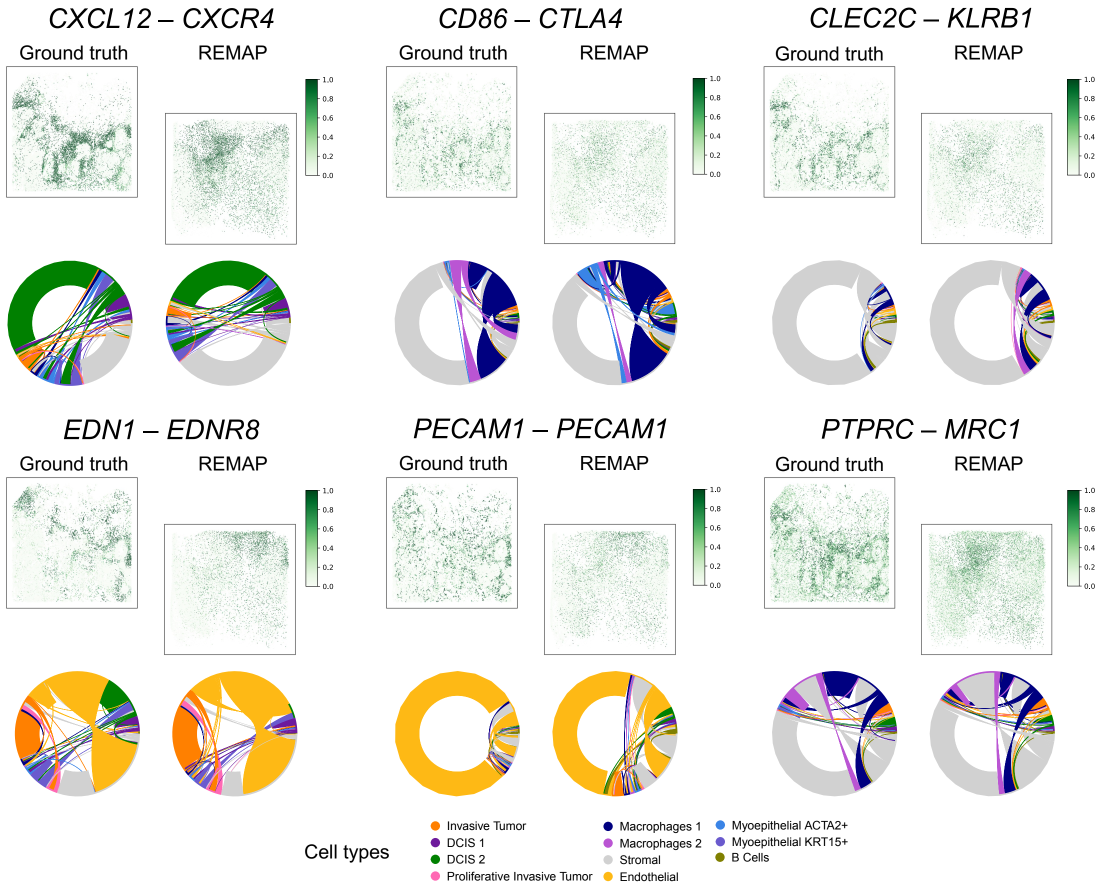

**Supplementary Fig. 22 |** Sankey plots showing the correspondence between ground-truth and predicted CN clusters for REMAP and LUNA in scenario 1 (**a**.) and 2 (**b**.) of the multi-capture ST analysis, with NMI between two sets of cluster memberships indicated.

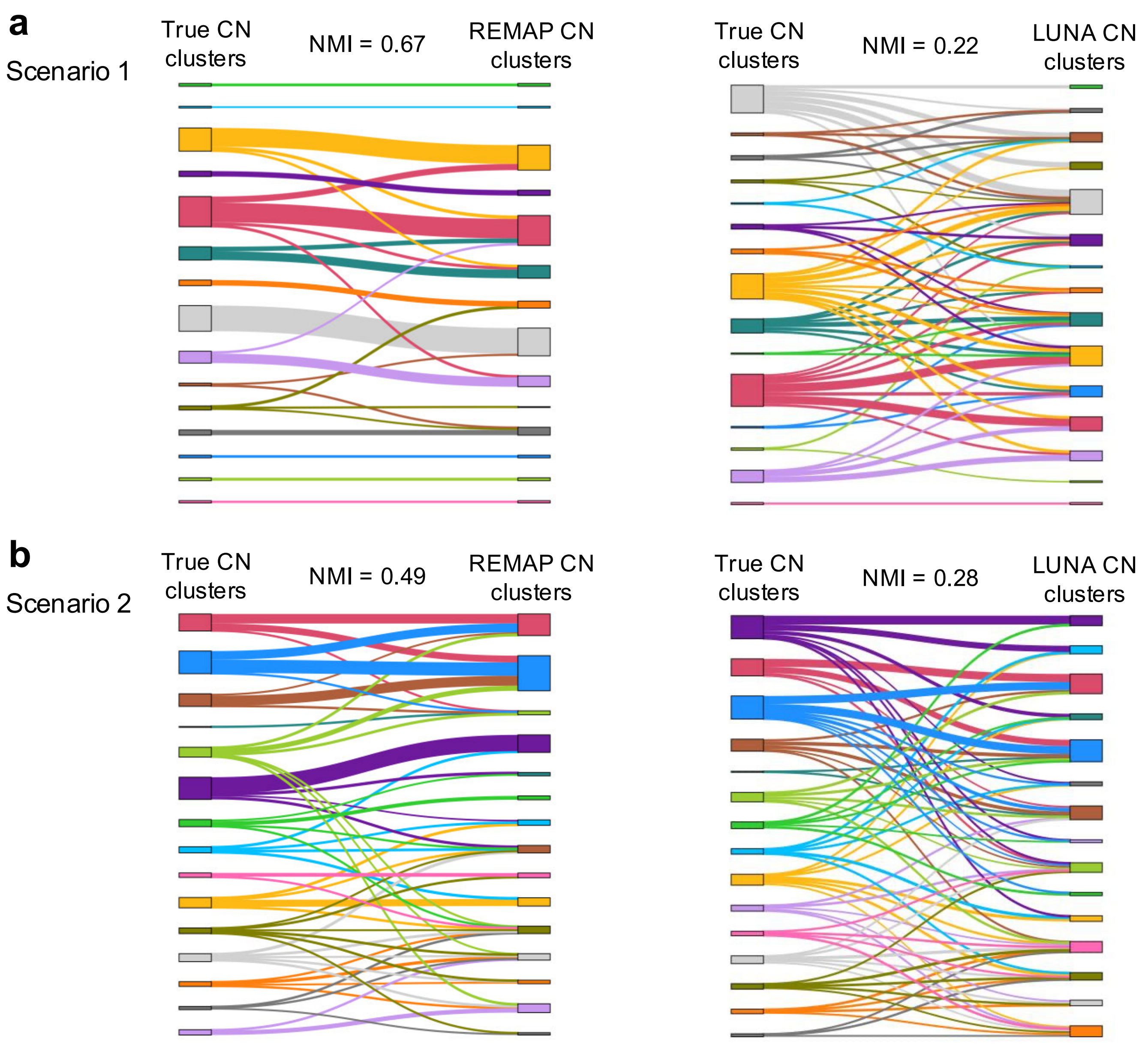

**Supplementary Fig. 23 | Benchmarking evaluations in the human gastric data. a**, Overview of study design. Multiple ROIs were selected, and within each ROI, half of the cells were selected as the reference, the other half and cells outside ROIs formed the testing set. **b**, Barplot of Pearson correlation between true and predicted pairwise distances. **c**, Line plots showing the NMI between true and predicted CN clusters across varying prespecified numbers of CN clusters. **d**, CN clusters and CN spatial proximity heatmaps constructed by REMAP and LUNA. In the top panel, cells were plotted under true locations and colored by predicted CN clusters; in the bottom panel, spatial proximities among CN clusters were shown as heatmaps. Clustering was performed on the predicted proximity matrices, and boxes indicated cluster memberships. **e**, Sankey plots showing the correspondence between ground-truth and predicted CN clusters for REMAP and LUNA, with NMI between two sets of cluster memberships indicated.

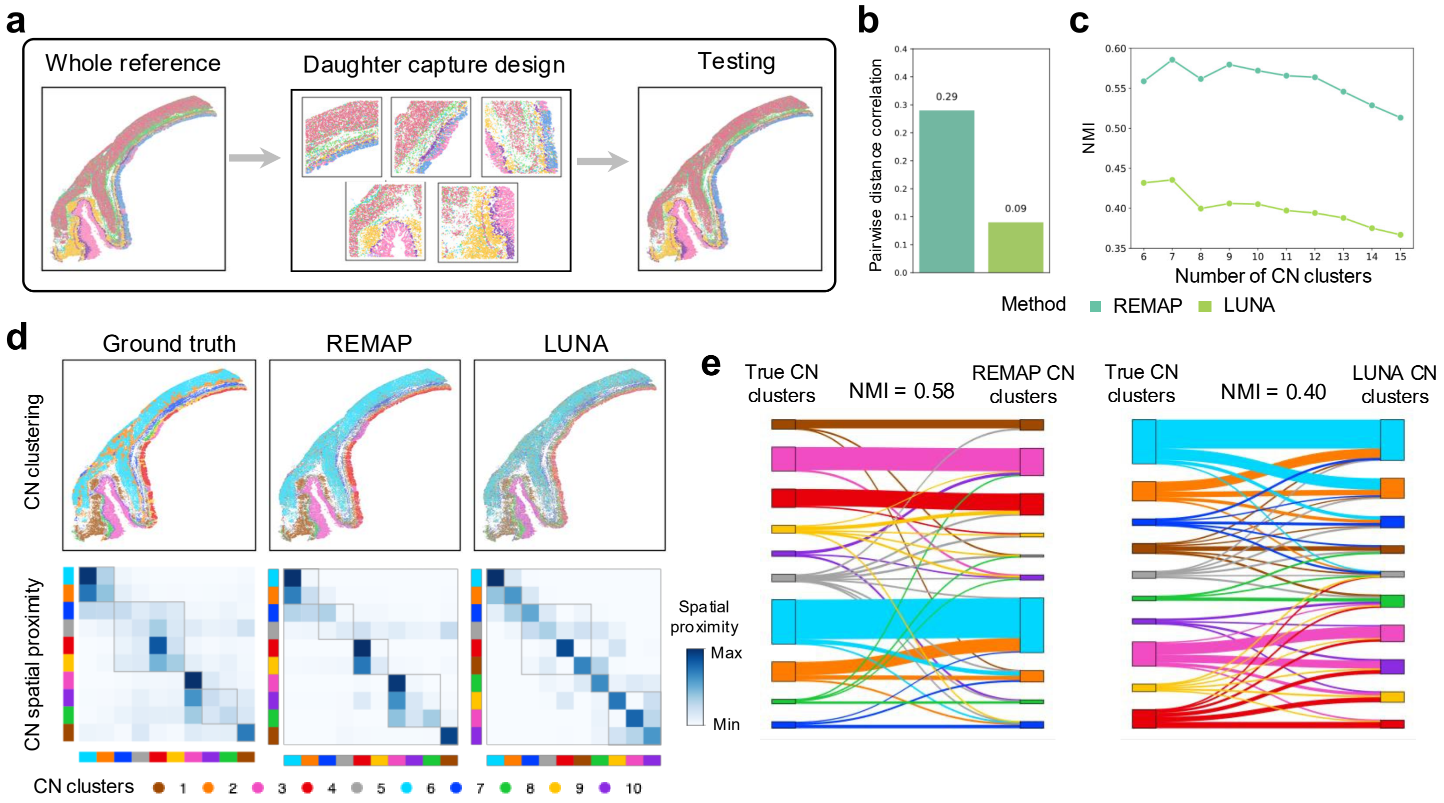

**Supplementary Fig. 24 | Benchmarking evaluations in the 3D whole mouse brain data. a**, Overview of study design. Mouse 1 with 147 consecutive 2D slices serves as the reference data, and Mouse 2 with 66 consecutive 2D slices serves as the testing data. **b**, Predicted 3D locations by REMAP, with a pairwise distance correlation of 0.73. **c**, Selected cell type-specific location prediction plots by REMAP.

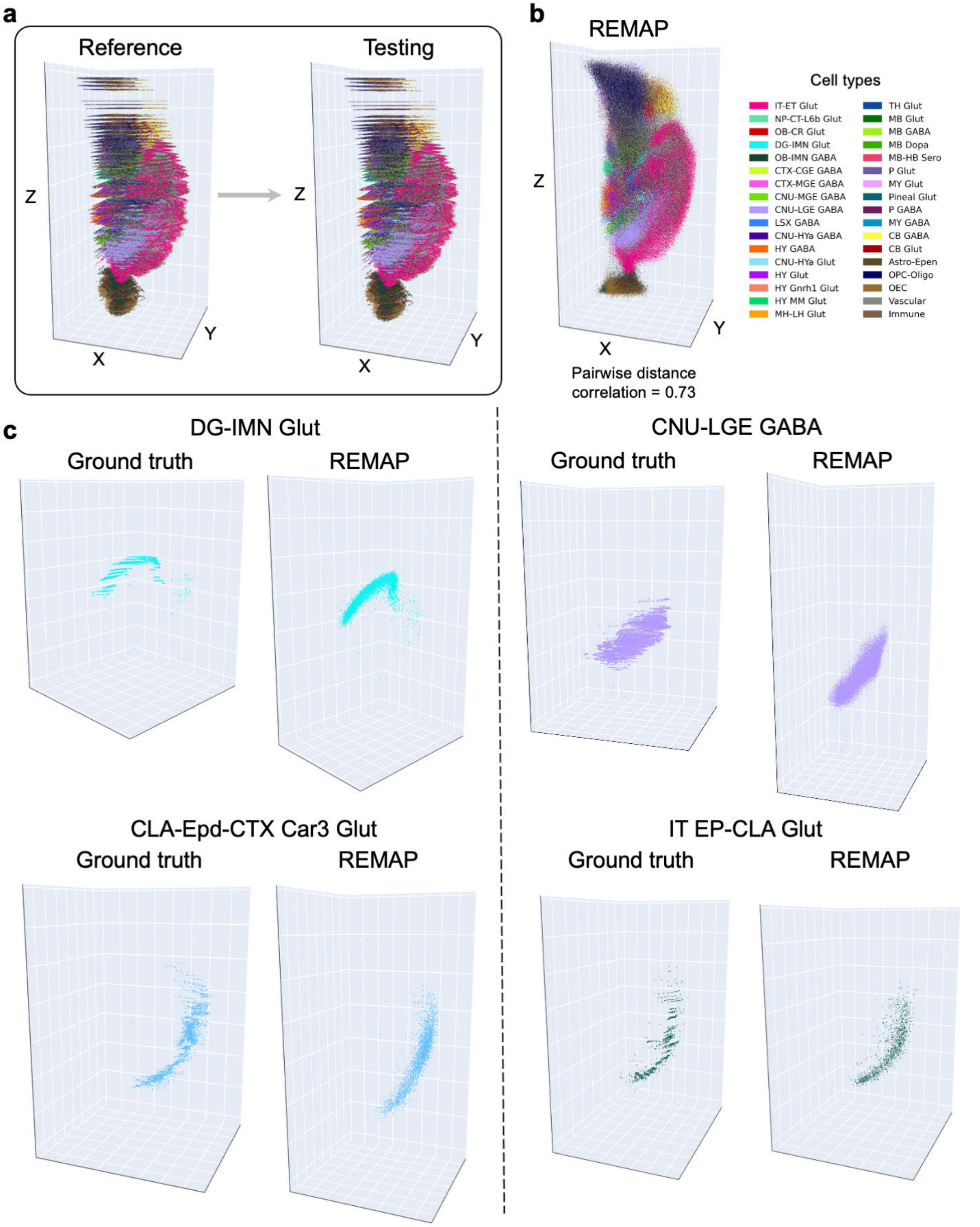

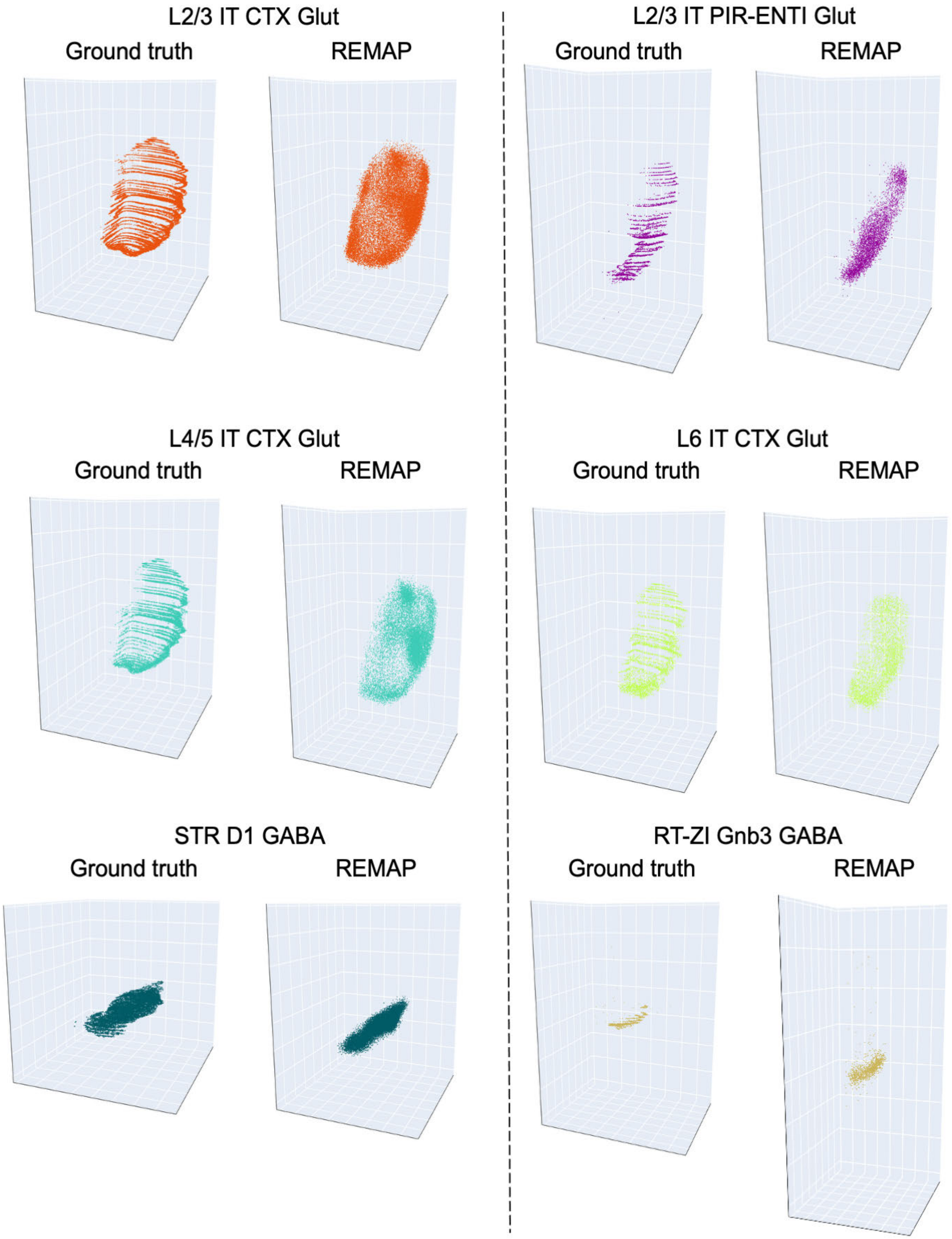
L2/3 IT CTX Glut
Ground truth
REMAP
L2/3 IT PIR-ENTI Glut
Ground truth
REMAP
L4/5 IT CTX Glut
Ground truth
REMAP
L6 IT CTX Glut
Ground truth
REMAP
STR D1 GABA
Ground truth
REMAP
RT-ZI Gnb3 GABA
Ground truth
REMAP

**Supplementary Fig. 25 | Location prediction plots by REMAP for additional samples in the human MS data.** Cell type colocalization matrices were then computed and compared with Visium deconvolution results, following the same procedure as in Fig. 5b.

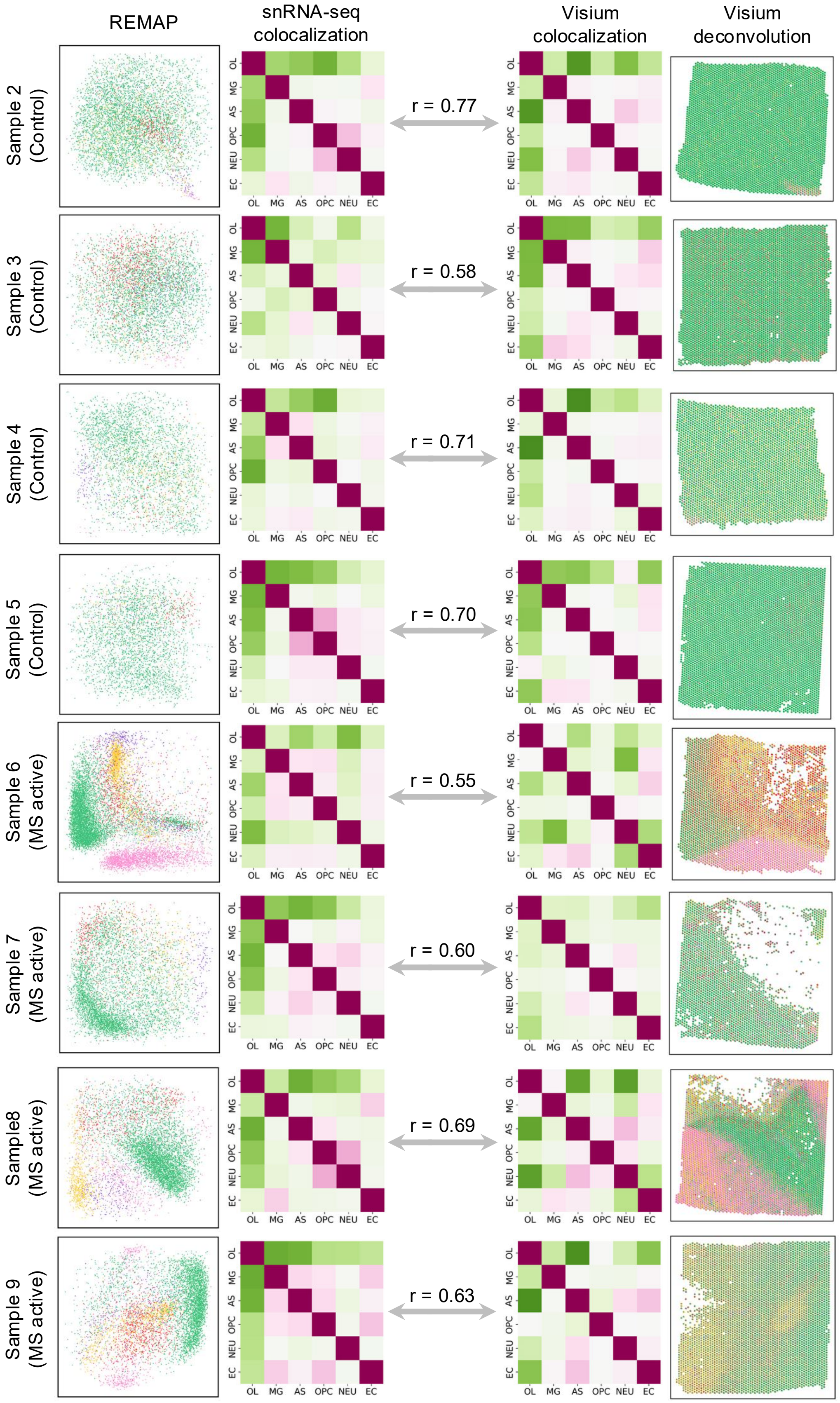

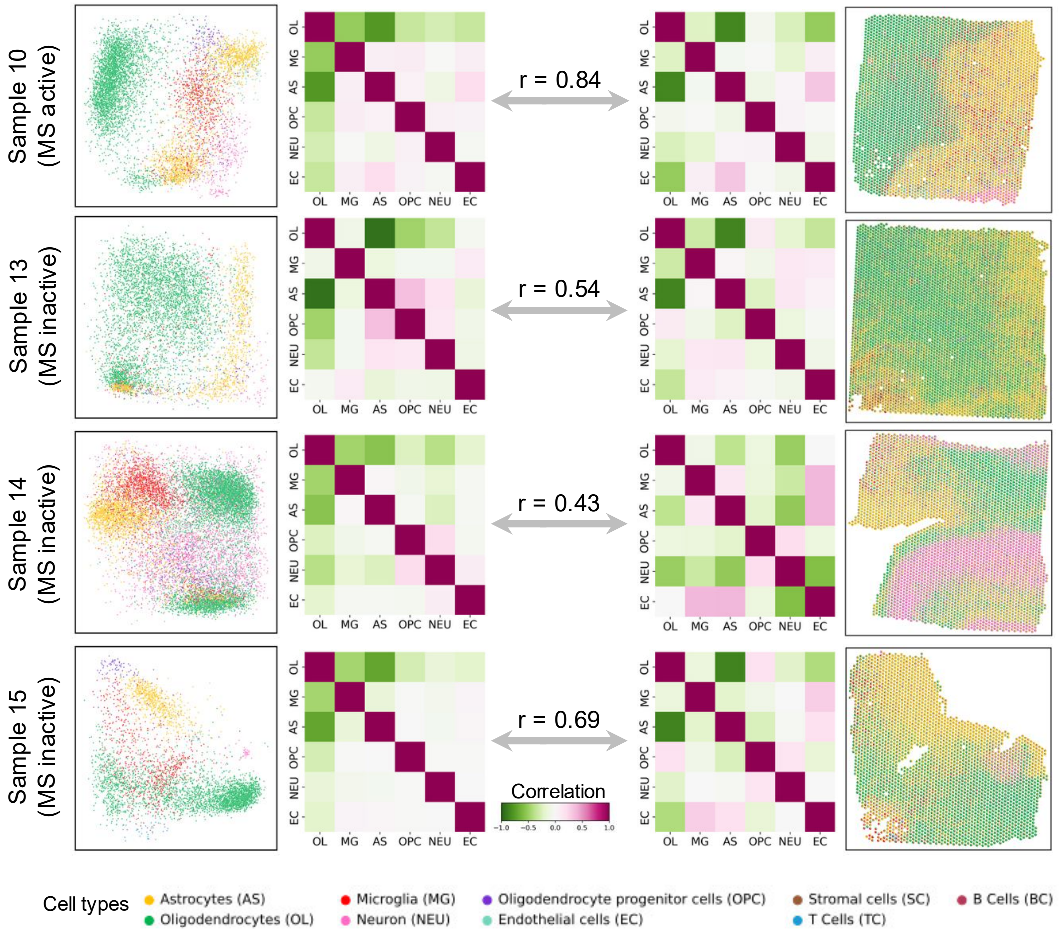
Sample 10
(MS active)
Sample 13
(MS inactive)
Sample 14
(MS inactive)
Sample 15
(MS inactive)
OL
MG
AS
OPC
NEU
EC
r = 0.84
r = 0.54
r = 0.43
r = 0.69
Correlation
−1.0
−0.5
0.0
0.5
1.0
Cell types
Astrocytes (AS)
Oligodendrocytes (OL)
Microglia (MG)
Neuron (NEU)
Oligodendrocyte progenitor cells (OPC)
Endothelial cells (EC)
Stromal cells (SC)
T Cells (TC)
B Cells (BC)

**Supplementary Fig. 26 | Bar plots comparing cell type composition between snRNA-seq and Visium data for each sample in the human MS data.**

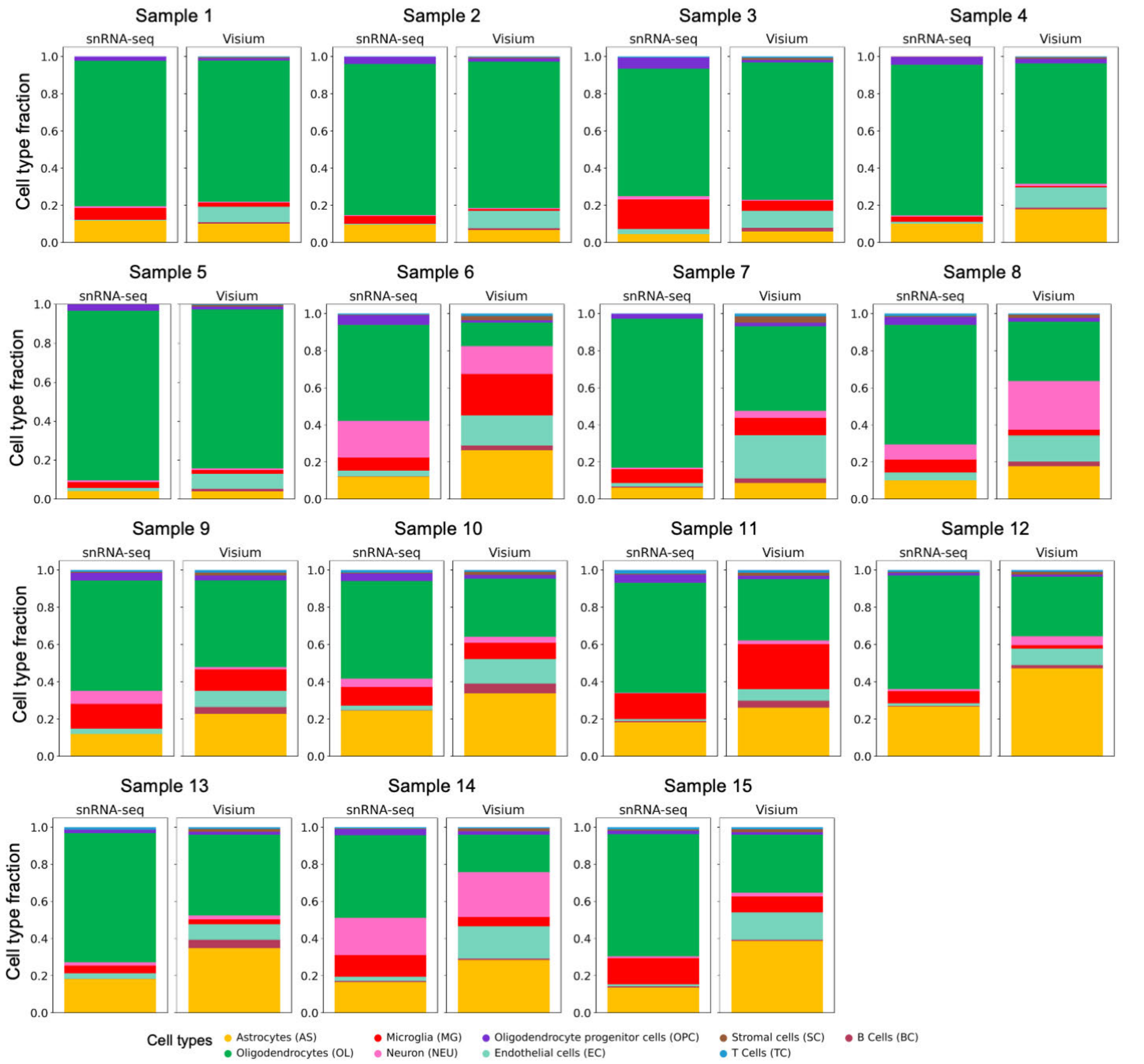

**Supplementary Fig. 27 | Ground-truth and predicted locations plots by REMAP and other methods for human cancers, including cervical cancer (a), ovarian cancer (b), melanoma (c), and lung cancer (d).** For cervical cancer, ovarian cancer, and melanoma datasets, besides REMAP, only CeLEry could process the full datasets.

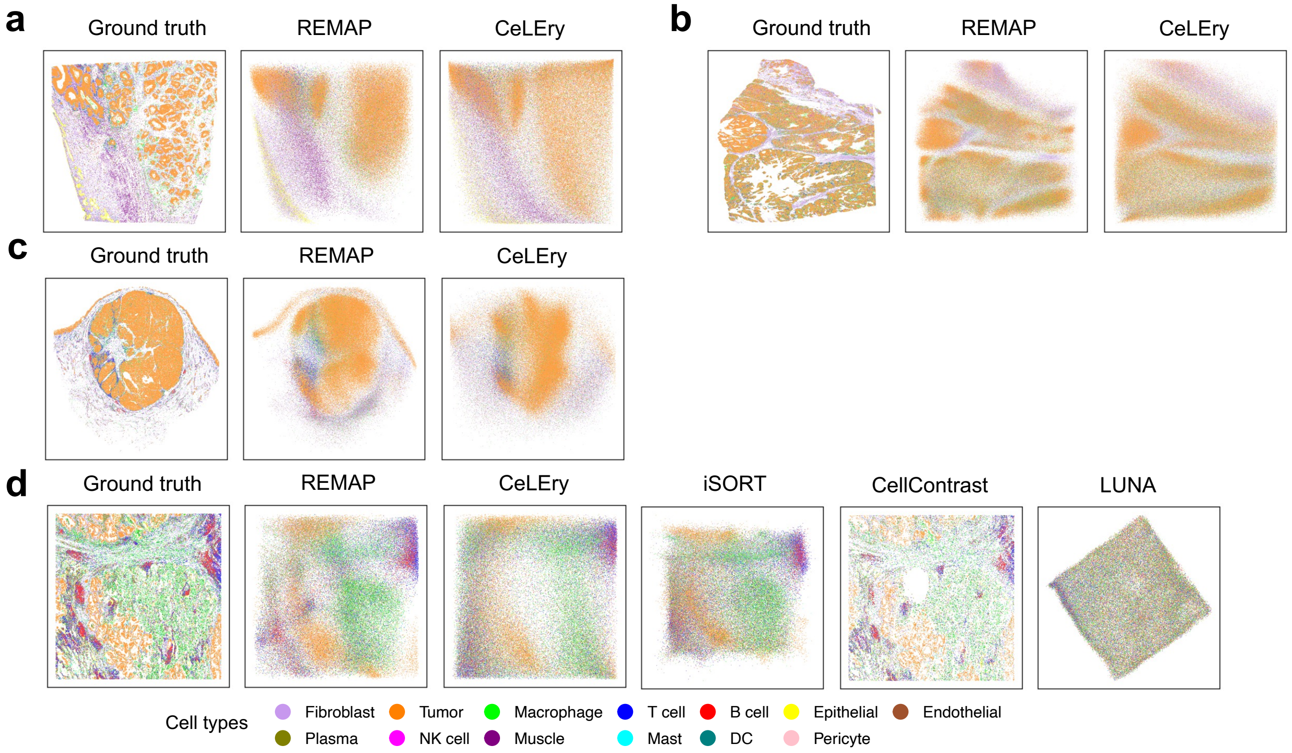

**Supplementary Fig. 28 | Bubble plots showing the expression of differentially expressed genes across NMF-derived CAF clusters for cross-subject location prediction in melanoma, using ground-truth (a) and REMAP-predicted locations (b).**

**a** Melanoma, cross patient, true locations

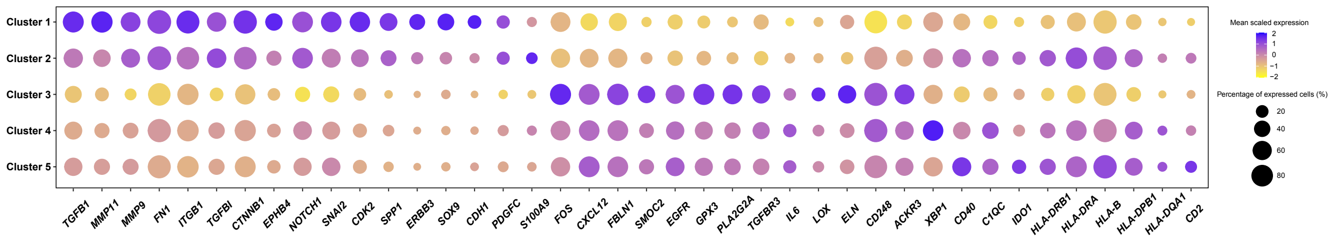


**b** Melanoma, cross patient, REMAP locations

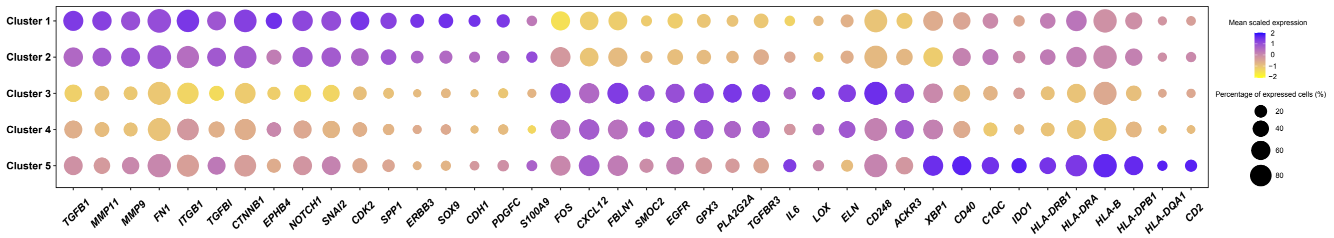

**Supplementary Fig. 29 | Bar plots of NMF-derived factors based on 3-hop neighborhood compositions of CAFs for within-subject location prediction under ground-truth and REMAP-predicted locations in cervical cancer (a) and ovarian cancer (b), melanoma (c), and lung cancer (d), with each region representing one hop.**

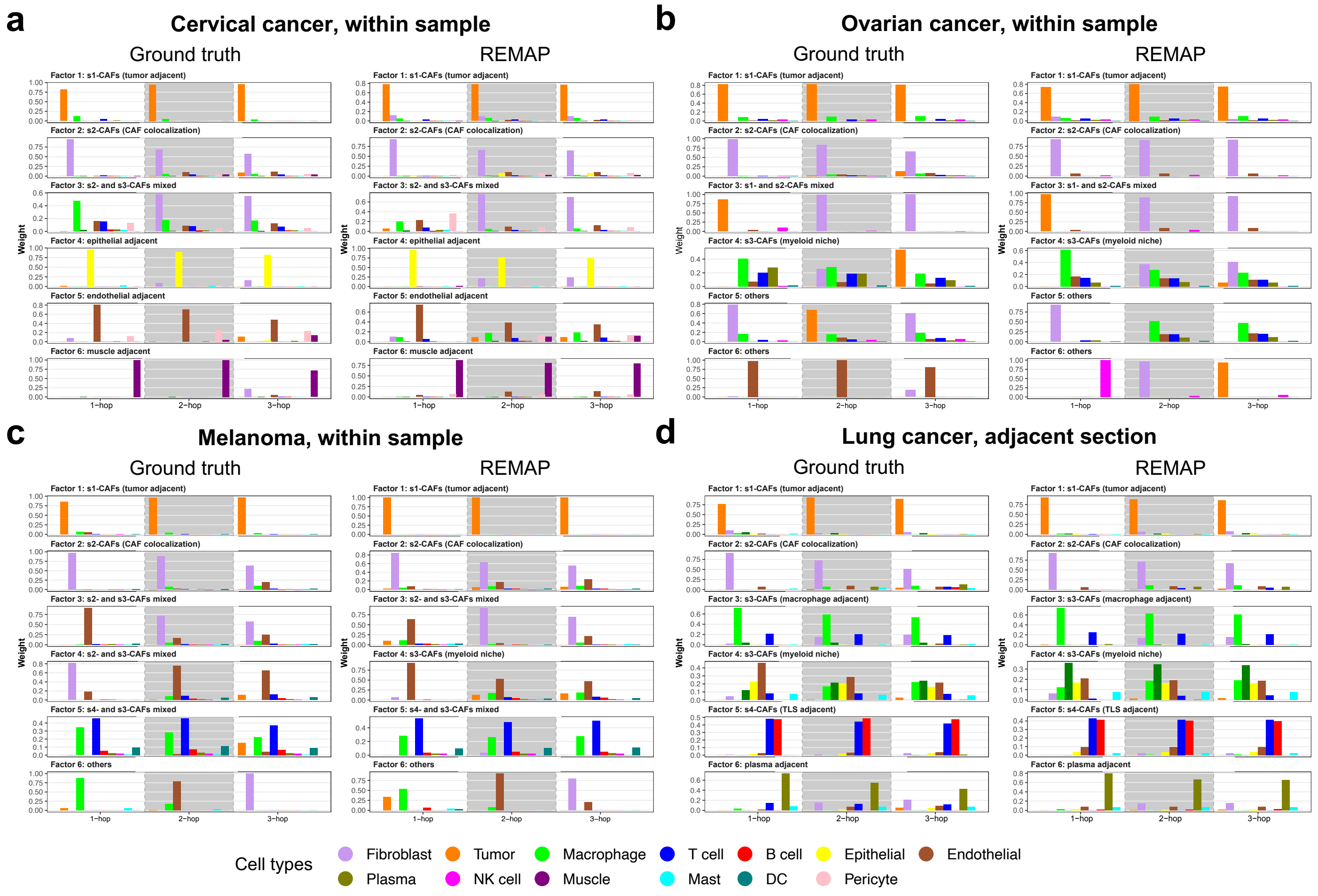

**Supplementary Fig. 30 | Applications to human prostate cancer. a-d**, Ground-truth and predicted cell locations by REMAP and CeLEry for within-subject location prediction. Other methods could not process the full dataset. **b**, Pearson correlation between true and predicted pairwise distances. **c**, Bar plots of NMF-derived factors based on 3-hop neighborhood compositions of CAFs for cross-subject location prediction using ground-truth and REMAP-predicted locations, with each region representing one hop. **d**, Bubble plots showing the expression of differentially expressed genes across NMF-derived CAF clusters for cross-subject location prediction, using ground-truth and REMAP-predicted locations.

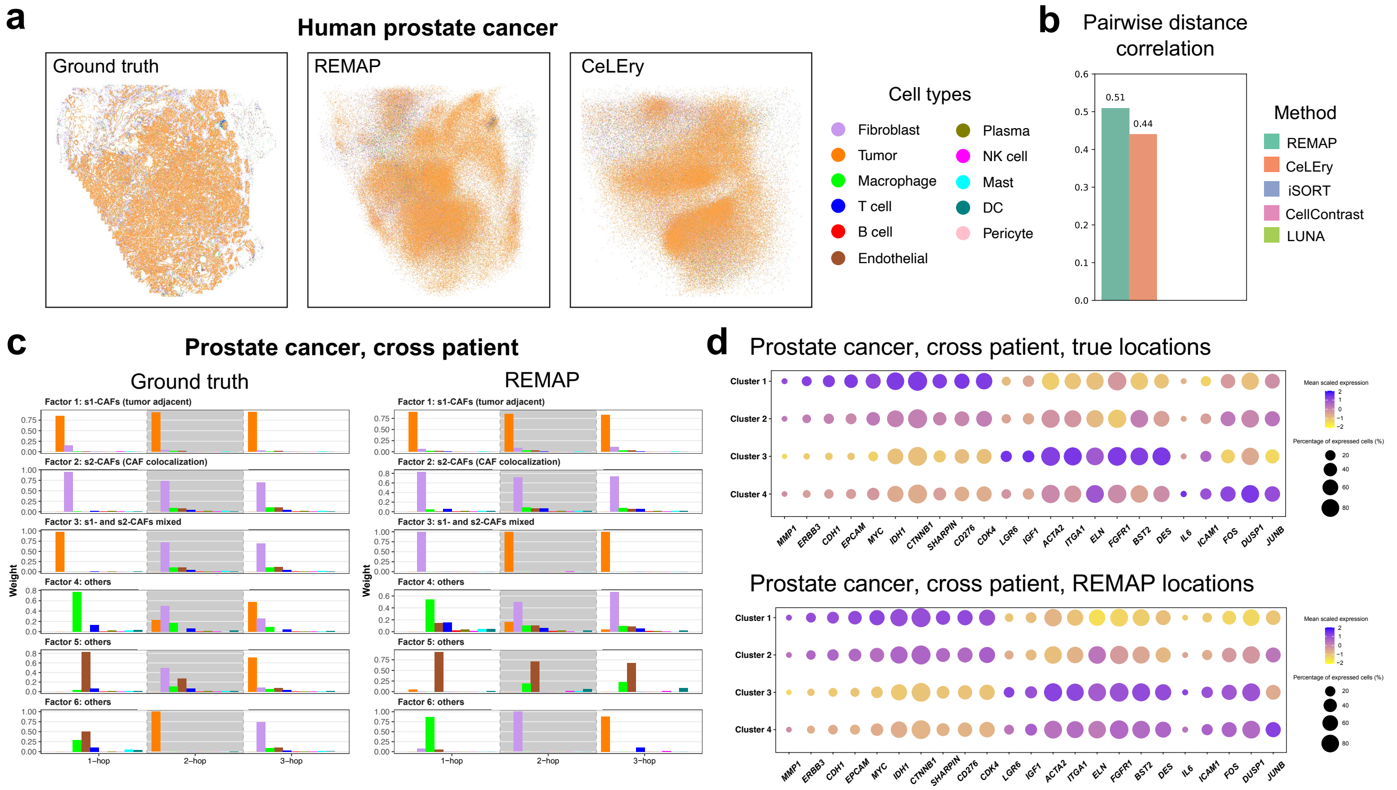

**Supplementary Fig. 31 | Running time comparison across REMAP, CeLEry, iSORT, CellContrast, and LUNA when evaluated on mouse primary cortex, mouse brain, and human melanoma datasets.** Due to the memory issue, iSORT, CellContrast, and LUNA failed to process the full human melanoma dataset.

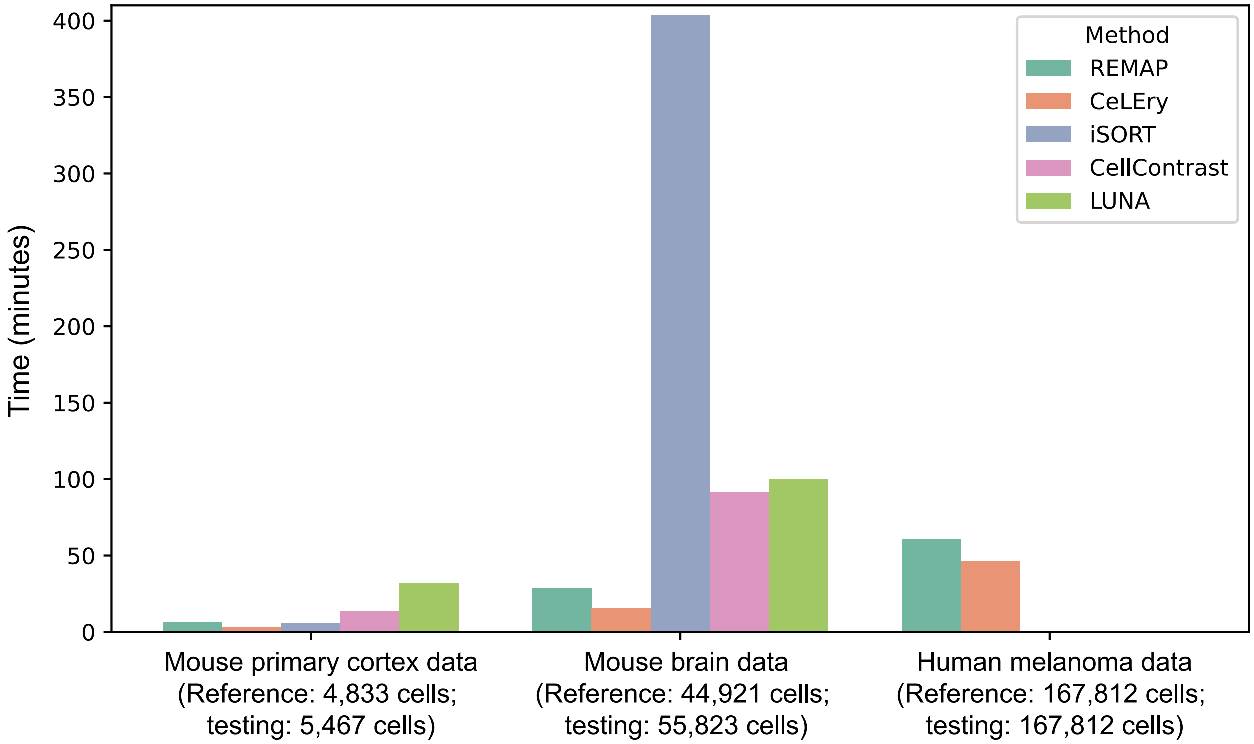
